# A generalized scaling law reveals synchrony-driven reorganization of brain dynamics across the human lifespan

**DOI:** 10.64898/2026.01.16.699865

**Authors:** Suman Saha, Chittaranjan Hens, Priyanka Chakraborty, Tomasz Kapitaniak, Gustavo Deco, Arpan Banerjee, Syamal Kumar Dana, Dipanjan Roy

## Abstract

Complex ecological systems obey universal scaling laws such as Taylor’s law, which links population variance to the sample mean. Whether other complex systems with integrated and segregated functional networks—such as the human brain—follow analogous scaling laws, and how these laws behave under synchronized activity, remains unknown. Progress has been hindered because neural signals at all measurement scales contain both positive and negative values, whereas Taylor’s law was developed for non-negative ecological data. Here we introduce a generalized spatial scaling law that replaces the sample mean with the root-mean-square (RMS) of detrended activity, extending Taylor-like scaling to signed multivariate time series in neural and other complex systems. Within this framework, we quantify synchrony using a dedicated metric of functional coordination and show analytically, and via numerical simulations using multivariate Poisson, binomial, uniform and gamma distributions, that the scaling exponent is inversely related to synchrony. Applying this approach to human fMRI data from three large lifespan cohorts (*N* = 840, ages 18–88) reveals distinct age-related trajectories of the scaling exponent, with healthy aging characterized by a substantial synchrony-induced reduction in the exponent. The synchrony–scaling relationship remains stable during resting-state activity but progressively shifts during naturalistic tasks across the lifespan, and is particularly strong in limbic, subcortical, and cerebellar subnetworks, indicating preservation at subnetwork levels. Finally, individuals with ADHD exhibit disrupted synchrony–scaling coupling, highlighting the potential clinical and translational utility of this generalized scaling metric.

## Introduction

Complex systems often exhibit scale-invariant fluctuations that reveal deep organizing principles [Munn bet al., 2024, D’Angelo and Jirsa, 2022,Tagliazucchi et al., 2012,He et al., 2010], yet we lack a unifying law that connects such variability to the coordinated dynamics of the human brain. Universal scaling relations such as Taylor’s law link mean and variance in non-negative ecological counts, but they do not directly apply to signed, detrended neural signals or explain how large-scale synchrony shapes brain-wide variability. This gap limits our ability to interpret ubiquitous power-law signatures in neuroimaging [Tomasi et al., 2017,Miller et al., 2009,Kello et al., 2010, within Tagliazucchi et al., 2012,Zimmern, 2020] a common theoretical framework and to relate them to healthy aging and brain disorders.

Here we develop a generalized spatial scaling law that extends Taylor’s law to multivariate signals with positive and negative fluctuations by replacing the sample mean with the spatial root-mean-square (RMS) of detrended activity. This simple reformulation yields a closed-form link between a spatial scaling exponent and emergent synchrony, allowing coordinated dynamics to be read out from scale-free variability in a principled way. We combine analytical derivations, simulations and large functional MRI cohorts spanning the adult lifespan to show that synchrony-driven changes in the scaling exponent track age-dependent reconfiguration of brain-wide coordination, remain stable at rest but reorganize during naturalistic tasks, and are selectively disrupted in neuropsychiatric conditions. By connecting coordination dynamics, fluctuation scaling and brain health within a single framework, our results position synchrony-induced scaling laws as a compact and general marker for complex neural systems. While examining brain-wide coactivation patterns, the multivariate analysis significantly enhances the sensitivity of human neuroimaging data [Kamitani and Tong, 2005,Haynes and Rees, 2005,Kriegeskorte et al., 2006] compared to conventional single-location-based approaches, e.g., temporal scaling law [Tomasi et al., 2017] or temporal autocorrelation [Shinn et al., 2023]. Coordinated cortical activity or collective dynamical behavior is often characterized as synchrony [Weistuch et al., 2021, Schölvinck et al., 2010, Zhao et al., 2019, Pikovsky et al., 2003, Sirovich et al., 2006, Banerjee et al., 2012, Zhang et al., 2019]. It is a tendency of neural population in distinct brain areas to be correlated through time [Sirovich et al., 2006,Banerjee et al., 2012,Zhang et al., 2019] . More generally, a coherent collective dynamics emerges in a network of dynamical units, as widely studied across disciplines, including physics [Pikovsky et al., 2003], ecology [Hanski and Woiwod, 1993,Post and Forchhammer, 2004,Liebhold et al., 2004, Cohen and Saitoh, 2016], and engineered systems [Witthaut et al., 2022]. One of the examples of universal scaling is Taylor’s law (TL), also known as fluctuation scaling, that has been verified with non-negative data in diverse complex systems, including ecology [Taylor, 1961,Smith, 1938,Maurer and Taper, 2002, Döring et al., 2015, Kuo et al., 2016, Xu et al., 2017], physics [Botet et al., 2001, Botet and P loszajczak, 2000,Vaughan and Uttley, 2007,Fronczak and Fronczak, 2010, Cohen and Xu, 2015], climatology [Jánosi and Gallas, 1999, Dahlstedt and Jensen, 2005], economics [Lee et al., 1998], real-world networks [De Menezes and Barabási, 2004, Argollo de Menezes and Barabási, 2004, Duch and Arenas, 2006, Eisler and Kertész, 2005,Yook and De Menezes, 2005, Šuvakov and Tadić, 2006], human population [Cohen et al., 2013,Yang et al., 2022], life science [Kendal, 2002,Woolhouse et al., 2001,Bar-Even et al., 2006], and neuroscience [Tomasi et al., 2017]. TL is described by a scalar relation between variance (*v*) and mean (*µ*) of non-negative data distributions, *v* = *aµ*^*b*^, where *a* is a constant and *b* is the exponent, predominantly existing in the range [1/2, 1] [Eisler et al., 2008]. Upon a logarithmic transformation, the equation becomes log(*v*) = *b* log(*µ*) + log(*a*). While obeying the power law, since there is no characteristic scale (size) for the phenomena under investigation, it is referred to as *scale-free* [Barabási and Albert, 1999, Marquet et al., 2005], *Among the two variants of TL, temporal* and *spatial*, we focus on the spatial TL that takes mean and variance of measurements over locations at different time instances, to explain the effect of coordinated brain activity on the scaling law.

Available evidence suggests that emergent coordinated dynamics in ecological population, measured as synchrony, can significantly affect the spatial TL exponent [Reuman et al., 2017]. A classic example in ecology is the Moran effect, where synchrony among spatially separated population alter the exponent [Reuman et al., 2017, Plank and Pitchford, 2017]. While theoretical and empirical links between synchrony and the scaling law (SL) have been realized [Eisler et al., 2008, Ballantyne Iv and Kerkhoff, 2005,Cohen and Saitoh, 2016,Engen et al., 2008,Reuman et al., 2017], prior works overlooked the bias introduced by the temporal mean and trends that is common in real-time or empirical data obtained from natural and experimental systems [Wu et al., 2007,Bunde et al., 2005]. We revisit this fundamental problem in data analysis that may have shadowed the underlying biophysical mechanism, and ask if the bias in measurement data is removed, how the presence and strength of the brain-wide coordinated activity influence the slope *b* of TL, and if so, how? A detrending operation [Wu et al., 2007] alleviates bias, ensuring stationarity and preserving spectral as well as correlation properties, while transforming data to span both negative and positive values; a methodological aspect overlooked in earlier works on fluctuation scaling. This development prompts an important question: Can the spatial scaling law be re-formulated to accommodate distributions containing both negative and positive values? This question becomes crucial when examining how synchrony governs universal scaling laws in complex systems, particularly, in the context of applying Taylor’s law in brain dynamics; this relationship remains largely unexplored. Extending Taylor’s law from its traditional domain of non-negative data distributions is therefore a crucial step forward to encompass the full range of values after detrending, while effectively isolating the effects of trends and temporal means in brain signals. Indeed, such an extension of the spatial Taylor’s scaling law makes it available for fluctuation analysis in other disciplines, including many physical and experimental systems. To implement this generic application of Taylor’s law, we simply replace the sample mean by spatial root-mean-square (RMS), and derive a revised scaling law between the spatial RMS and sample variance. Although detrended fluctuation analysis has been widely employed in climatology, extreme-event studies [Bunde et al., 2005], and tipping phenomena [Dekker et al., 2018], however the scaling behavior of spatial aggregation in detrended data with suppressed temporal mean and trends remains unexplored. More importantly, the effect of synchrony on spatial scaling was not widely explored earlier, except in the context of population segregation in ecology.

The brain-wide co-ordinated hemodynamics, in general, has been formalized as a temporal alignment or coactivation of distributed cortical regions during specific functional states [Sirovich et al., 2006,Banerjee et al., 2012, Zhang et al., 2019]. The coodinated hemodynamics or blood-oxygen-level-dependent (BOLD) activity, obtained from functional magnetic resonance imaging (fMRI), is quantified by a global metric, namely, synchrony by averaging functional connectivity (FC) matrix (in this work, it is similar to the variance of spatial mean derived from the multivariate time series). The FC is obtained from pairwise Pearson correlations between the BOLD signals (typically spanning 10 minutes) across all cortical regions [Greicius et al., 2003, Friston, 2011, Bressler and Menon, 2010, Friston, 2003, Beckmann et al., 2005,Damoiseaux et al., 2006,Power et al., 2011,Yeo et al., 2011,Chakraborty et al., 2024].

This synchrony measure captures the extent of neural coordination and large-scale coactivation patterns elicited in response to internal or external stimuli, and holds significant clinical relevance [Courtiol et al., 2020,Sahoo et al., 2020,Pathak et al., 2022,King et al., 2018]. Earlier studies investigated temporal correlations in brain oscillations as physiological correlates of behavior, highlighting scale-free 1*/f* ^*β*^ noise as a hallmark of complexity [Gilden et al., 1995], and reporting long-range correlations using the measure of synchronization errors during human coordination tasks [Chen et al., 1997]. Research utilizing EEG and MEG data has demonstrated self-organized criticality in brain oscillations through detrended fluctuation analysis and frequency–power scaling, suggesting that critical-state dynamics enable neural network reorganization [Linkenkaer-Hansen et al., 2001]. Furthermore, analyses of healthy human fMRI data revealed that temporal TL exponents change with age and vary across brain regions [Tomasi et al., 2017]. Yet, a substantial gap remains concerning the generic role of synchrony-induced spatial fluctuation scaling across diverse brain states, particularly, in relation to healthy aging and neural disorders. Building upon our revised scaling law, we examine how the coactivation patterns, as indicated by synchrony, modulate the fluctuation scaling exponent, and thereby uncover empirical and theoretical links between synchrony and the spatial SL in detrended multivariate time series, spanning negative to positive values. By quantifying coherent BOLD activity across multiple brain regions and subnetworks, we assess how aging affects synchrony-induced scaling across different task conditions throughout the lifespan, comprising both normative aging and neural disorders.

## Results

Our approach is illustrated in Fig. 1 using exemplary BOLD signals from a single participant. The multivariate time series data of size *T* × *n* (where *T* is total time length; *n* is brain regions or locations) is first detrended. Time series from four locations (*n*=4) are presented in green, red, blue and yellow. The detrending operation removes additional trends and sets the temporal mean to zero from each time series. The detrended data *Y*_*i*_(*t*) at each instant *t* in all locations follows a near-normal distribution. Next, amplitudes of the detrended BOLD signal of all the brain regions are squared individually and summed at a time instant *t* to obtain spatial RMS (*m*) at time *t*. Similarly, we store the spatial variance (*v*) at each time instant. The spatial SL exponent (*b*) is derived from the linear relationship between log_10_(*m*) and log_10_(*v*) taken at different time instants. Long-term correlation of measurements between a pair of locations is derived by Pearson correlation coefficient, giving a correlation matrix. Synchrony Ω is the var(*µ*(*t*)) (where *µ* is the sample mean) derived from the correlation matrix, see SI for detailed derivation. The association slope, *β* is finally obtained by a linear regression fit that describes the influence of synchrony Ω on the spatial SL exponent *b*. We rounded up to 1e-6 decimal values in all our simulations. For detailed derivations and descriptions, see Methodology section and supporting information (SI).

**Figure 1.**
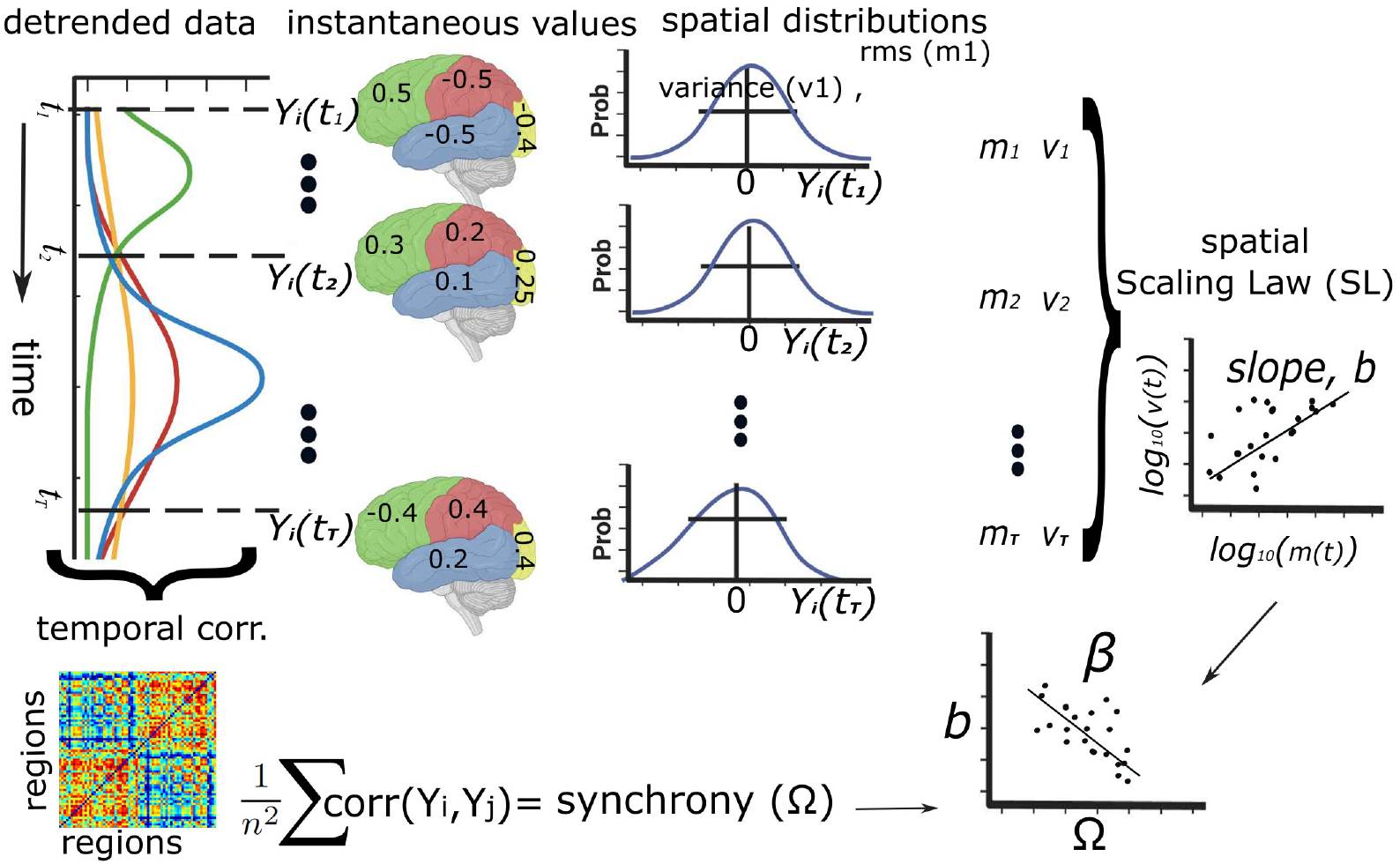
Schematics of sequential processing and analysis of data. This illustrates how synchrony-scaling association has been derived from the detrended spatio-temporal time series data.

### Numerical simulations: Effect of synchrony on the scaling law

For modeling correlated local dynamics, we generate multivariate time series data from a zero-mean Gaussian distribution with a specified covariance matrix. The time series dimension is *T* =260 time points and *n*=116 regions, matching empirical data as discussed below. Figures 2(a, b) show the log-normal distributions of spatial RMS log(*m*) and variance log(*v*), in logarithmic scale, respectively, as exemplary normal distributions N(0, 1). The spatial fluctuation scaling between *m* and *v* is shown in a log_10_-vs-log_10_ scatterplot in Figs. 2(c) in the case of i.i.d. data, when long-term correlations between any two locations, *ρ*=corr(*Y*_*i*_, *Y*_*j*_) ≃ 0 for *i* ≠ *j*. The exponent is estimated from semi-analytical and linear fitting as *b* = 1.988 and 1.991, respectively, shown in Fig. 2(c). The correlation between the spatial SL (*b*) and synchrony (Ω) is illustrated in Figs. 2(d). The spatial SL exponent remains almost around 2 (red line) when the strength of synchrony is very weak (*ρ* = 0), and the origin is a random process. For identically distributed but not necessarily independent data, the synchrony substantially influences the spatial SL exponent; see Figs. 2(e-g) for *ρ*=0.001, 0.01, and 0.1. The spatio-temporal time series data is constructed depending on the correlation value *ρ* using the *common element method* [Fischer, 1933]; see SI for details and other distributions. Figure 2(h) shows that an increasing degree of synchrony induces a monotonic decrease of the SL slope as obtained from analytical results using Eq. (3). The inset shows the fitting errors quantified by root mean squared errors (RMSE) between the actual log(*v*) values and *b* log(*m*) values from the linear regression of log(*m*)-vs-log(*v*). The spatial SL exponent, based on spatial RMS and variance, thus decreases monotonically with the degree of synchrony. This is independent of distributions of data as numerically confirmed with other distributions, namely, Poisson, negative binomial, uniform, and gamma distributions (see SI for detailed results in Figs. S2-S13). Note that irrespective of the original distribution of measurements, after the detrending operation, the data follow a normal distribution.

**Figure 2.**
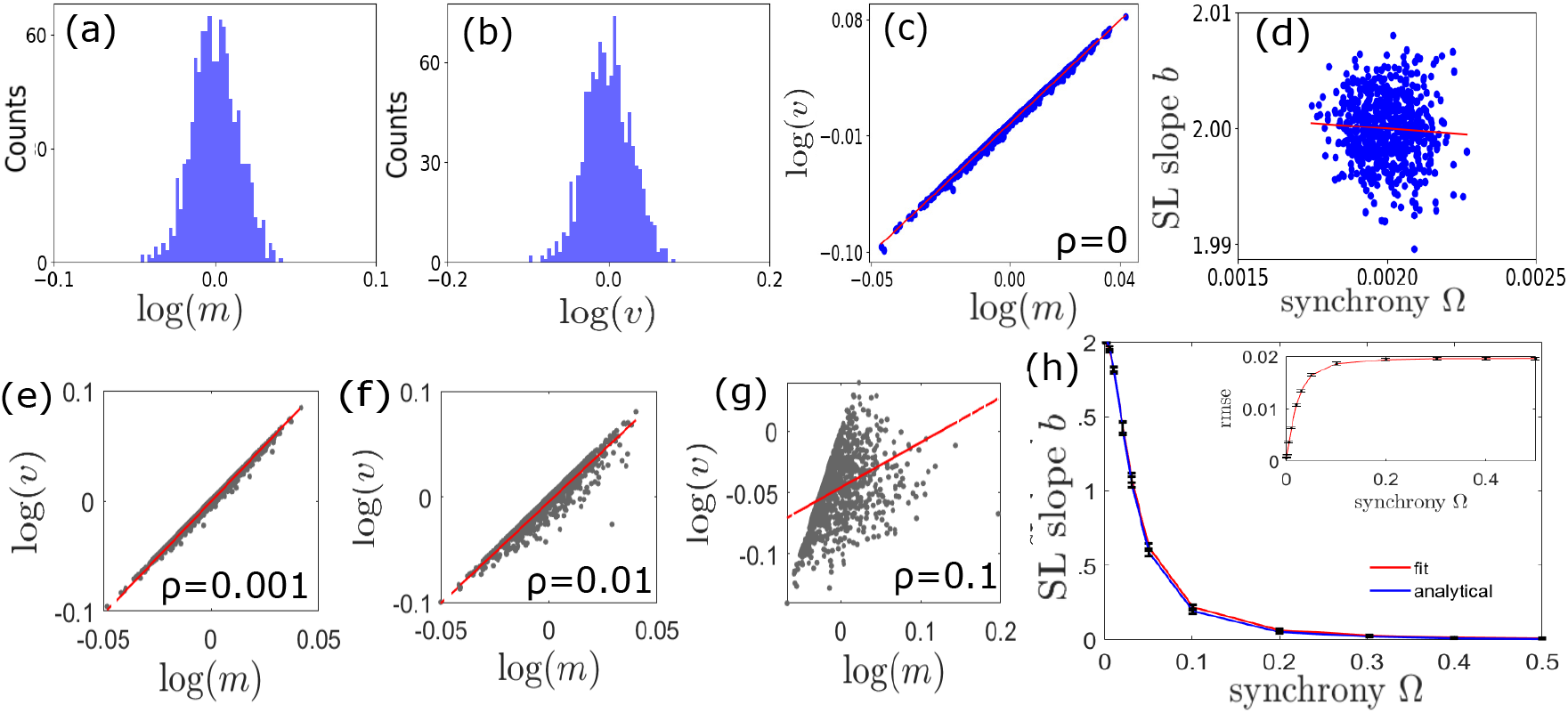
Numerical example: Analysis of spatio-temporal time series. Time series data at each location, generated by randomly drawn values from normal distribution. (a, b) Log-normal distributions of spatial RMS and variance. (c) Linear fit of spatial log(*m*) vs. log(*v*), the spatial SL slope *b* ≃ 2, when long-term correlation between any pair of locations is almost zero (*ρ* ≃ 0). (d) The SL slope plotted against synchrony for 645 independent realizations. (e-g) For identically distributed but not necessarily independent data, the effect of synchrony (Ω) on the SL for a model in different sampling locations in the multivariate time series drawn from normal distribution using the common element method [Fischer, 1933] given that *ρ*=0.001, 0.01, 0.1. The scaling exponent decreases with increasing correlation *ρ*. (h) The SL slope *b* against synchrony Ω is plotted using Eq. (3) that confirms the numerical trend in the scaling exponent. Inset shows the root mean squared error (RMSE) derived from the linear regression of log(*v*)-vs-log(*m*). Error bars depict standard deviation for 50 independent realizations.

### Empirical data distribution: Effect of detrending

Figure 3(a) Empirical BOLD signals from 116 brain regions of one young adult during a resting state. Non-zero temporal means and trends are prominent in the raw data of one location, as shown in

**Figure 3.**
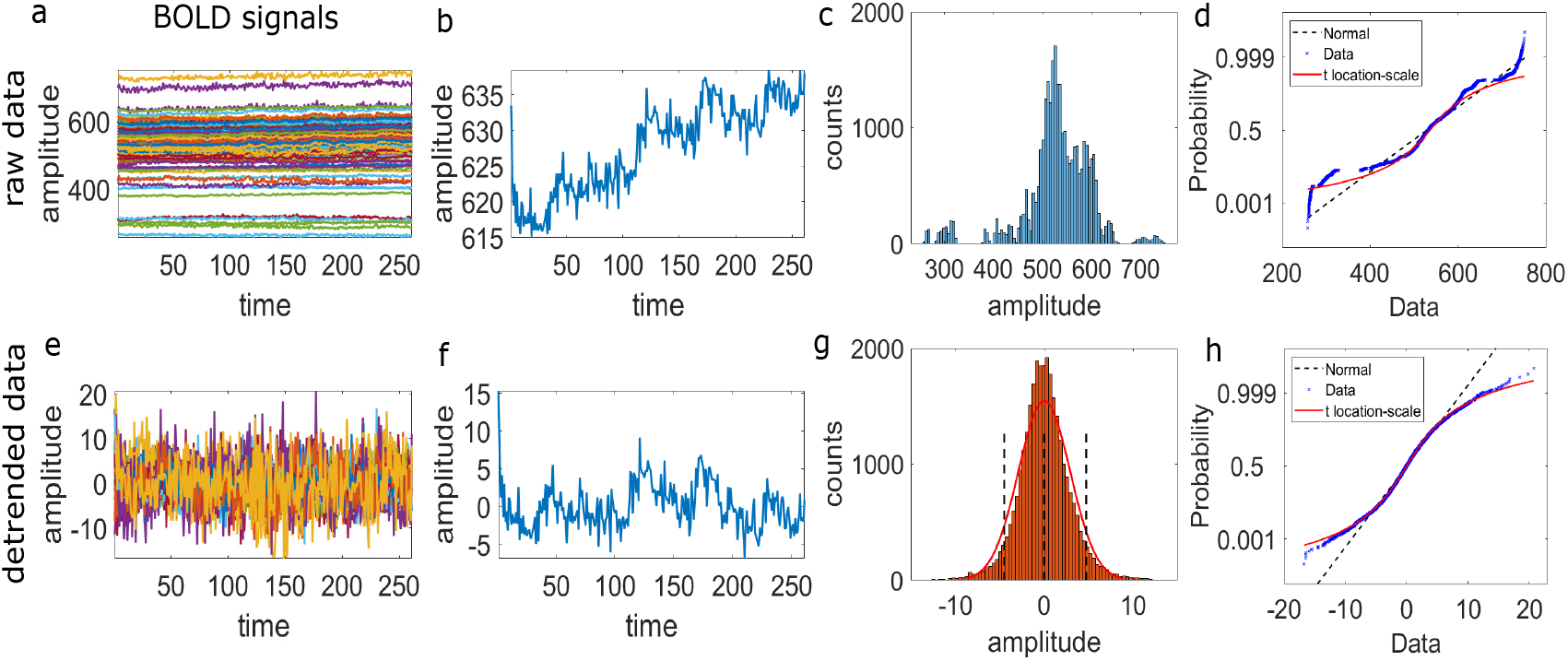
Empirical data distributions: Before and after detrending operation. (a) Empirical BOLD series in the resting state of one participant. Data size is 261 × 116; brain regions = 116 (AAL atlas), time points = 261. (b) BOLD signal of left cerebellum-10 (one location). (c) Histogram plot of the raw data. (d) The normal probability plot shows that neither the normal line (black dashed line) nor the location-scale curve (red) fits very well. (e) After detrending, the BOLD series of all the brain regions is shown to spread across zero. (f) Suppressed trend of the BOLD signal of left cerebellum-10, compare (b). (g) Data distribution with normal fit of the blue curve in (f). The vertical black dashed lines indicate 95% CI. (h) Normal probability plot of the detrended data shows that it fits well to the normal distribution for the data range within 95% CI.

Fig. 3(b). Figures 3(c,d) present a histogram and normal probability plot of the raw data. The data fit neither the normal line (black dashed line) nor the location-scale curve (red). Figure 3(e) illustrates that following a detrending operation, BOLD signals across all brain regions oscillate around a zero temporal mean with both positive and negative fluctuations. The detrending removes trends and temporal mean, as illustrated in Fig. 3(f) for one brain region. The detrended spatiotemporal BOLD data tend to follow a normal distribution, as confirmed by the plots in Figs. 3(g, h) for one exemplary brain region. The histogram and normal probability plots for the detrended data further confirm the normal distribution fit within a 95% confidence interval (CI). We perform the detrending operation on 116 brain regions of all the participants (*N* =840) using MATLAB and Python built-in functions. Results for the same subject during two naturalistic tasks are shown in Figs. S15 and S16 in SI. Also, subject-wise spatial SL exponents are computed and plotted for three brain states. See SI, Figs. S17 and S18 for log-normal plots of spatial RMS and spatial variance, quantile-quantile plots, and spatial RMS-versus-variance scatter plots.

### Task-driven coordinated dynamics and spatial scaling law

We examine the influence of any emergent coordinated dynamics or synchrony on the spatial SL exponent for both the resting state and active tasks using fMRI data from a large cross-section of adult lifespan (18-88 years), *N* =645 participants from CamCAN dataset (BOLD time series of size 260 time points×116 brain locations) [Taylor et al., 2017, Shafto et al., 2014]. The SL slope is plotted against synchrony for resting state (rest), sensorimotor task (smt), and movie watching (mov) task in Fig. 4, respectively, arranged in order from top to third row. We have considered two methods to derive the slopes, *b*, using Eq. (3), and linear regression fitting (shown in SI; Fig. S19). Factors other than synchrony that may have influenced the exponent are examined from the raw correlations between *b* and Ω for the three brain states shown in Figs. 4(A,D,G) for rest, smt, and mov, respectively. The relation between the scaling exponent and synchrony is significantly negative in all three brain states. Higher synchrony Ω is related to lower slope *b*, despite possible confounding influences. A higher association between the SL slope (*b*) and synchrony (Ω) may be attributed to contributions from marginal distribution.

**Figure 4.**
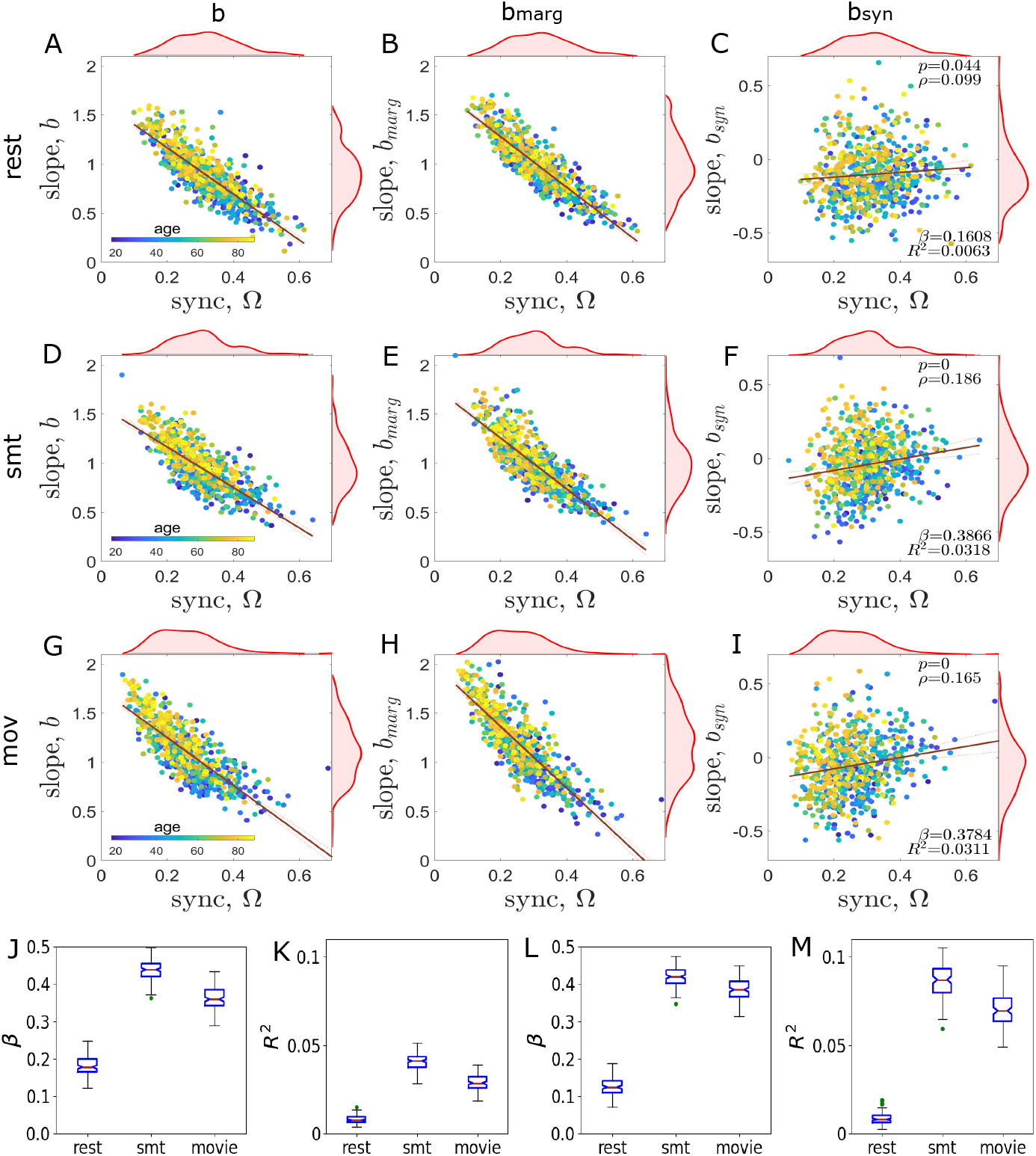
Spatial SL exponent against synchrony for different brain states. (A-C) resting state, (D-F) sensorimotor task, and (G-I) movie watch. (A,D,G) Scatter plots (colored circles) and linear regression fit (red lines) between exponent *b* and synchrony Ω in three brain states. (B,E,H) Contributions of marginal distributions, slope *b*_*marg*_ against synchrony Ω. (C,F,I) Synchrony-driven SL slope (*b*_*syn*_) after removing marginal effects. Statistical significance is considered when *p <* 0.005. (J,L) The association between *b*_*syn*_ and Ω is captured by *β*, together with goodness of fit *R*^2^ in (K,M) for 100 realizations. Results are shown for semi-analytical (J,K) and fitting-based analysis (L,M).

To account for the actual influence of synchrony, we remove the effect of marginal distributions that may have influenced the slope in conjunction with the changes due to synchrony. We decompose the scaling exponents, *b* = *b*_*marg*_ + *b*_*syn*_, where *b*_*marg*_ accounts for the contributions from time series marginals and *b*_*syn*_ due to synchrony. The scaling exponent *b*_*marg*_ is approximated from independently randomizing phases [Prichard and Theiler, 1994] of the empirical BOLD signals (refer to Section *Surrogate time series* in SI), and recomputing the log(*v*) vs. log(*m*) slope, which ensures that marginals do not contribute to the slope, and thereby to extract the influence of synchrony. Figures 4(B,E,H) show correlation between the spatial SL (*b*_*marg*_) vs. synchrony (Ω) in three brain states. Next, we derive *b*_*syn*_=*b* − *b*_*marg*_ to estimate the synchrony-induced SL exponent (*b*_*syn*_), which is only observed after masking the effects from the time series marginals and shown in Figs. 4(C, F, I). Changes in three different brain states delineate distinct patterns of association between the scaling exponent (*b*_*syn*_) and synchrony (Ω). At rest, we find no significant correlation between the two metrics (Fig. 4(C); *p >* 0.005). Whereas in sensorimotor (Fig. 4(F); *p <* 0.005) and movie-watching tasks (Fig. 4(I); *p <* 0.005), the scatterplot fit shows significantly positive correlations between *b*_*syn*_ and Ω, i.e., the SL exponent *b*_*syn*_ increases with increasing strength of synchrony. The Spearman correlation coefficient (*ρ*), and p-value (*p*), association slope (*β*), and adjusted *R*^2^ values are denoted in Figs. 4(C,F,I). The slope *β* is estimated by a least squares linear regression fit of the *b*_*syn*_ against the Ω plot. The parameter *β* in three brain states is verified with 100 independent realizations. Figures 4(J) and 4(L), obtained from semi-analytical and fitting-based results, respectively, show similar changing patterns; the correlation *β* is low in the resting state but significantly higher in movie-watching (mov) and sensorimotor tasks (smt). Figures 4(K) and 4(M) show corresponding goodness of fit (*R*^2^) for analytical and fitting-based results, respectively.

### Reduced synchrony-scaling association in aging brain

We have separated the CamCAN data into three age groups: (i) young adults (YA: 18-27 years), (ii) middle-aged adults (MA: 46-55 years), and (iii) older adults (OA: 75-84 years). The correlations between synchrony and the spatial scaling exponent are presented for three age groups in three columns; see Figs. 5(A,D,G) for YA, Figs. 5(B,E,H) for MA, and Figs. 5(C,F,I) for OA. The three brain states, rest, smt and mov, are in order from top to bottom rows, respectively. The association between the exponent *b*_*syn*_ and synchrony Ω remains significantly positive in all three mental states in YA; see Figs. 5(A,D,G). In MA group, *β* values significantly decrease in advanced age compared to YA, as seen in Figs. 5(B,E,H), which further decreases in OA group; see Figs. 5(C,D,I). The Spearman correlation (*ρ*) and p-value (*p*), association slopes *β*, and adjusted-R^2^ derived from a linear fit between synchrony Ω and *b*_*syn*_ are mentioned in the figures. The strength of correlation, measured by the associated slope *β*, has a tendency to rotate clockwise with increasing age in all three states.

**Figure 5.**
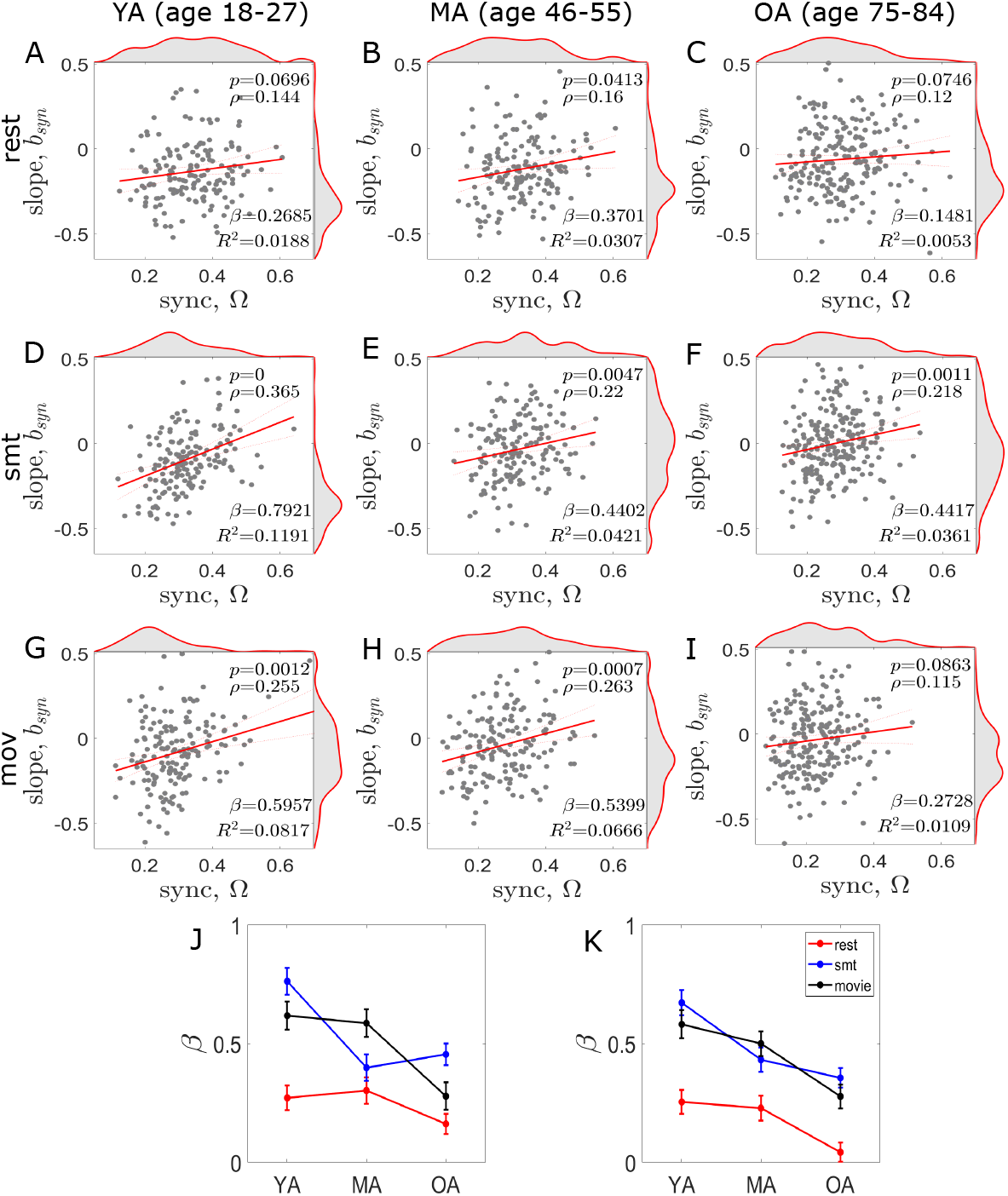
Group-wise age-related alterations in the spatial exponent. Associations between the spatial SL (*b*_*syn*_) and synchrony (Ω) are shown for three age groups in *rest* (A-C), *mov* (D-F) and smt (G-I) states. Three age groups are created using 10-year age bins. (A,D,G) In young adults (YA), the slope (*β*) remains positive and significant in all three conditions. In middle-aged adults (shown in (B,E,H)), *β* rotates in clock-wise direction compared to young adults, and (C,F,I) rotates further in clock-wise direction in older adults. The slope *β* shows a tendency to rotate clockwise with increasing age in all three brain states. (J,K) Association between Ω and *b*_*syn*_ is monitored by the slope *β* for three age groups in three brain states, where left and right plots are obtained from the semi-analytically derived and fitting-based SL slopes, respectively. Consistency in *β* is checked by 100 independent realizations. The error bars indicate variations over the trials. *p<*0.005 implies statistical significance.

Furthermore, we present the results for 100 independent realizations by randomizing BOLD phases of individual subjects to check consistency in the association (measured by the slope *β*) between synchrony (Ω) and SL slope (*b*_*syn*_) for the three age groups in Fig. 5(J, K), where left (J) and right (K) figures are plotted using semi-analytical approach in Eq.(3) and a numerical fitting-based SL exponent, respectively. The association of *β* remains almost similar for both the YA and MA adults in the resting state and movie-watching task (red and black lines, respectively), but largely declines in older adults. It seems that *β*, in the sensorimotor task (blue line), significantly decreases across the adult lifespan. For fitting-based results, see Fig. S21 in SI.

### Preserved synchrony-scaling relationship in subnetworks

To delineate the contributions of the subnetworks in age associated changes on spatial SL against synchrony in empirical data, we separate seven functional subnetworks [Krzemiński et al., 2020] for the AAL atlas [Tzourio-Mazoyer et al., 2002], such as: default mode network (DMN), fronto-parietal network (FPN), sensorimotor network (SMN), visual network (VN), limbic, subcortical, and cerebellum regions. DMN includes 12 ROIs: middle orbitofrontal (ORBmid L/R), inferior orbitofrontal (ORBinf L/R), anterior cingulate (ACG L/R), posterior cingulate (PCG L/R), angular gyrus (ANG L/R), precuneus (PCUN L/R) [McGill et al., 2012, Krzemiński et al., 2020]. FPN includes middle frontal (MFG L/R), frontal inferior triangularis (IFGtriang L/R), superior parietal (SPG L/R), inferior parietal (IPL L/R), supramarginal (SPG L/R), and angular gyrus (ANG L/R) [Wolf et al., 2015, Krzemiński et al., 2020]. SMN includes precentral (PreCG L/R), rolandic operculum (ROL L/R), supplementary motor area (SMA L/R), superior temporal (STG L/R), and postcentral gyrus (PoCG L/R) [Clemens et al., 2013, Krzemiński et al., 2020]. VN includes calcarine (CAL L/R), cuneus (CUN L/R), lingual (LING L/R), middle occipital (MOG L/R), inferior occipital (IOG L/R), superior occipital (SOG L/R), fusiform (FFG L/R) [Liu et al., 2007]. Limbic regions include anterior cingulum (ACG L/R), posterior cingulum (PCG L/R), dorosal cingulum (DCG L/R), hippocampus (HIP L/R), parahippocampal (PHG L/R), superior temporal pole(TPsup), and mid temporal pole (TPmid L/R) [Tzourio-Mazoyer et al., 2002]. Subcortical regions include amygdala (AMYG L/R), caudate (CAU L/R), putamen (PUT L/R), pallidum (PAL L/R), thalamus (THA L/R)) [Tzourio-Mazoyer et al., 2002]. The subnetworks are shown in the first and second rows in Fig. 6. In the resting state, the synchrony-induced SL exponent (*b*_*syn*_), captured by the slope *β*, is found to be more prominent in limbic, subcortical, and cerebellum regions compared to the other four subnetworks, . The visual network (fourth column) shows a significant correlation between the synchrony and SL exponent in the movie watching task compared to the other two brain states. In the DMN and SMN, movie watching is profoundly observable, compared to the resting state and smt, whereas in FPN, the resting state condition is more dominating. Overall, the limbic, subcortical, and cerebellum regions show higher correlation between the synchrony and SL exponent.

**Figure 6.**
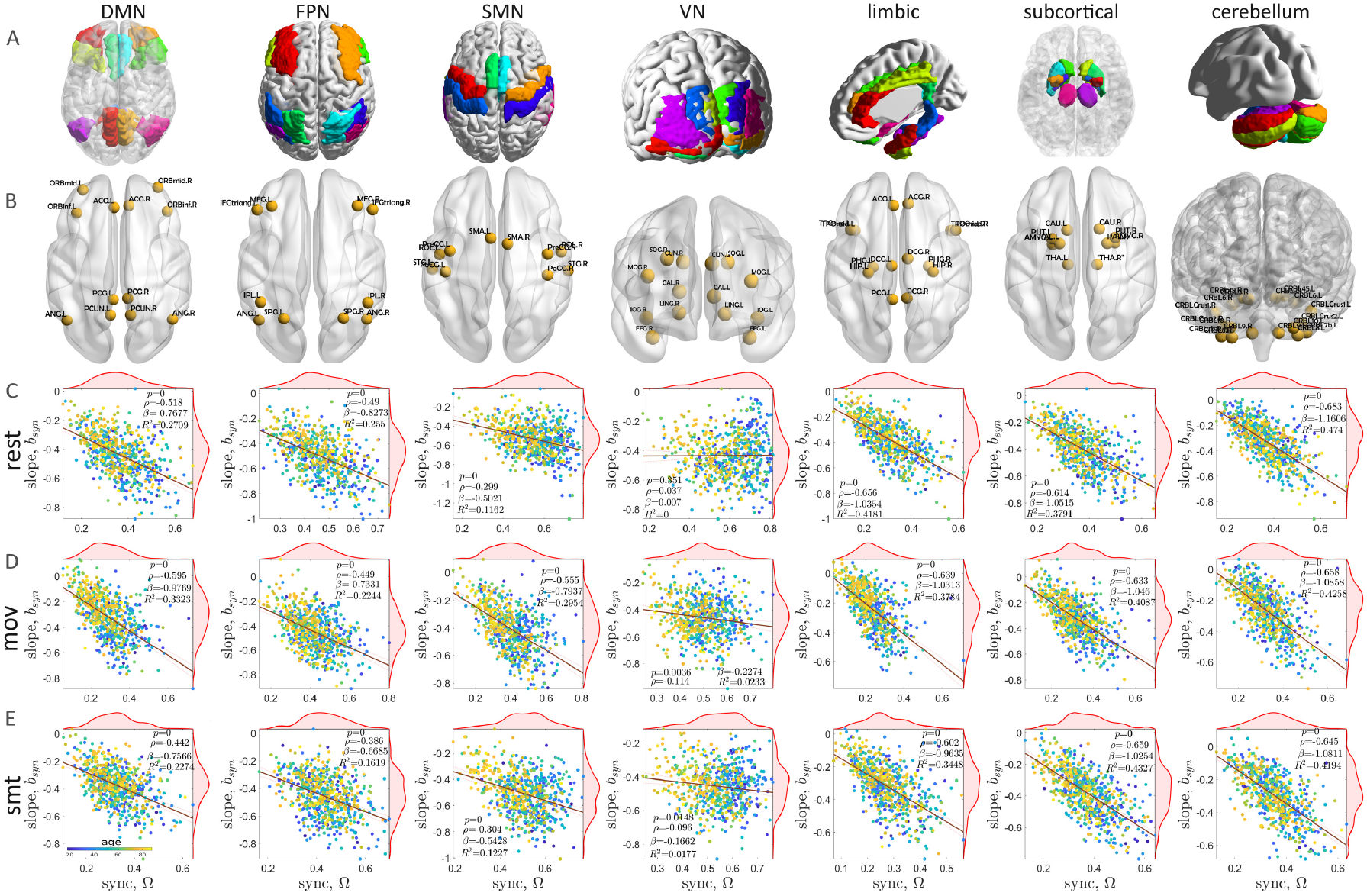
Spatial SL slope against synchrony is shown for seven subnetworks for the three brain states. (A, B) Visualization of the seven subnetworks in the brain, including cortical and subcortical regions involved in neurodevelopment and aging. Subnetworks are defined based on the AAL atlas. (C-E) The scatterplot (colors represent age) of spatial SL slope, *b*_*syn*_, against synchrony, Ω, during resting state, movie watching, and sensorimotor task are shown.

### Cross-validation, clinical relevance and comparison with other metrics

We test generalization across independent datasets and clinical populations using two independent cohorts: NKI/Rockland and UCLA datasets. Cross-validation in the NKI/Rockland dataset (*N* =89) replicates the negative relationship between synchrony and spatial scaling exponents with aging (Fig. S22 in SI). Comparing young adults (ages 18-30) versus older adults (ages 59-83), the association slope (*β*) between synchrony and scaling exponents shows a significant clockwise rotation with age, consistent with CamCAN findings. Clinical validation in the UCLA-CNP dataset (*N* =106) reveals that individuals with ADHD (*N* =40) compared to healthy controls (*N* =66) shows a non-significant synchrony-induced spatial scaling relationship (*p>*0.05; Fig. S23 in SI). The association slope *β* is profoundly reduced in ADHD, confirmed across 1000 independent realizations, suggesting that neural disorders disrupt the coupling between brain-wide coordinated activity and fluctuation scaling patterns observed in healthy adults. These validations demonstrate that the synchrony-scaling relationship is robust across independent healthy aging cohorts and sensitive to neuropsychiatric conditions, supporting its utility as a biomarker of brain dynamics.

Finally, a comparison with the traditional metrics for brain-wide dynamics shows consistent patterns. Metastability [Deco and Kringelbach, 2016,Shanahan, 2010], entropy of functional connectivity dynamics [Hutchison et al., 2013, Sastry et al., 2022], and phase synchrony using Kuramoto order parameter [Kuramoto, 1984] all show minimal age-related changes in resting state but significant alterations during movie-watching and sensorimotor tasks (Fig. S24 in SI). These findings agree with the synchrony-induced spatial SL results [cf. Fig. S20 in SI], confirming that aging effects are stronger in task-evoked than resting-state activity across multiple measures of brain dynamics.

## Discussion

We have developed a generalized spatial scaling law that extends Taylor’s Law beyond its traditional domain of non-negative data, enabling applications to multivariate time series spanning negative to positive values. This methodological advance reveals fundamental connections between emergent synchrony and scale-invariant patterns in complex systems, with a particular relevance in understanding the coordinated brain dynamics across the human lifespan.

Our approach reformulates the spatial scaling law analysis by replacing the sample mean with the spatial root-mean-square (RMS) in the power-law relationship against spatial variance, thereby extending the existing framework to a broader class of data across various disciplines. This modification enables the analysis of spatial variance against RMS scaling even when observations exhibit negative-to-positive spreads that are not amenable to traditional Taylor’s law formulations. A special care of data is adopted using a detrending operation, to remove temporal trends and means before computing spatial moments, which mitigates nonstationarity and isolates spatial structure. This operation transforms a wide range of marginal distributions, including Poisson, negative binomial, gamma, and uniform, into approximately normal fluctuations with both negative and positive deviations, allowing them to be treated within a unified Gaussian-like framework. By working with detrended and RMS instead of raw means, the resulting spatial scaling law accommodates signed fluctuations that arise naturally in many empirical examples (e.g., environmental, ecological, or socio-economic fields). This generalization overcomes a key limitation of the classic spatial Taylor’s law, which implicitly assumes nonnegative counts and thus cannot capture variability patterns around zero or changing baselines.

The analytical derivation in Eq. 3 establishes that the spatial SL exponent *b* is inversely related to synchrony (Ω), with synchrony appearing in the denominator and thus directly modulating the slope. This relationship holds when marginal distributions remain fixed while correlations or coordinated dynamics vary, suggesting that long-range spatial correlations, rather than local statistical properties, drive deviations from the baseline exponent of *b*≃2. Numerical simulations across diverse probability distributions confirm the prediction that the exponent monotonically decreases as synchrony strengthens [see Figs. S2-S13 in SI].

We empirically validated the generalized scaling law using BOLD fMRI data from healthy adults aged 18-84 years during resting-state and naturalistic tasks in two independent cohorts (CamCAN, *N* = 645; NKI/Rockland, *N* = 89). Healthy aging was associated with a pronounced reduction in brain-wide average synchrony across three brain states (rest, movie watching, and sensorimotor task), consistent with prior reports of age-related disruption in neural coordination [Hinault et al., 2020,Weistuch et al., 2021, D’Esposito et al., 2003]. To mechanistically interpret these effects, we decomposed the observed spatial scaling exponent *b* into a marginal component *b*_marg_ and a synchrony-driven component *b*_syn_ using phase randomization, thereby isolating contributions from coordinated neural activity versus intrinsic signal statistics. This decomposition allowed effects of aging on large-scale coordination to be quantified independently of changes in local variance structure.

The impact of aging on spatial SL differed markedly between spontaneous activity and naturalistic tasks (SI, Fig. S20). During rest, aging exerted minimal influence on *b*_syn_ (SI Fig. S20, second column, first row), suggesting either that spontaneous hemodynamics are relatively resistant to synchrony-mediated scaling changes in later life, or that resting-state dynamics are less sensitive to age-related decline. In contrast, during tasks, aging significantly increased the spatial SL exponent *b*_syn_ (SI, Fig. S20, second column, second and third rows). Across all three brain states, average whole-brain synchrony decreased significantly with advancing age (SI, Fig. S20, third column). At rest, we observed no significant association between *b*_*syn*_ and synchrony, suggesting that spontaneous brain dynamics maintains relatively stable scaling relationships independent of brain-wide coordinated activity. In contrast, both sensorimotor and movie-watching tasks exhibited significant positive correlations between *b*_*syn*_ and synchrony, with association slope (*β*) markedly higher than resting values [cf. Fig. 4]. This task-dependent modulation indicates that externally driven or goal-directed brain states recruit coordinated activity patterns that fundamentally alter spatial scaling properties, a signature absent during unconstrained spontaneous activity.

Group-wise analyses in the CamCAN cohort revealed that young (18–27 years), middle-aged (46– 55 years), and elderly adults (75–84 years) exhibited a progressive clockwise rotation of the linear association slope *β*, which quantifies the influence of synchrony Ω on *b*_syn_, across the adult lifespan. We further replicated this pattern in the NKI/Rockland sample, comprising *N* = 45 young adults (18–30 years) and *N* = 44 older adults (59–83 years). The association slope *β* remained stable in young adults across all conditions but declined markedly in middle-aged participants and decreased further in older adults. This age-related attenuation of synchrony’s impact on spatial scaling was strongest during sensorimotor tasks, moderate during movie watching, and weakest at rest. Together, these findings suggest that aging selectively impairs the brain’s ability to regulate spatial scaling via coordinated activity in cognitively demanding or stimulus-driven states, while largely preserving the intrinsic scaling characteristics of spontaneous dynamics.

Network-specific analyses of seven major functional subnetworks showed that limbic, subcortical, and cerebellar regions exhibited the strongest synchrony–scaling associations across all three brain states, consistent with their roles in integrating distributed neural activity. The visual network showed minimal effects at rest but robust associations during movie watching, in line with the strong visual demands of the task. Default mode and sensorimotor networks demonstrated enhanced synchrony-induced scaling during movie watching relative to the other conditions, whereas frontoparietal networks showed comparatively stronger effects at rest, potentially reflecting domain-general control processes engaged during unconstrained cognition.

In the absence of biophysical coordination mechanisms, the spatial SL exponent approaches 2 under near-zero synchrony, reflecting the purely statistical aggregation of independent fluctuations. As coordinated cortical activity emerges through neural encoding and stimulus processing, synchrony increases and the exponent decreases monotonically. This monotonic dependence provides a quantitative bridge between collective dynamics and universal scaling behavior, positioning spatial scaling exponents as compact summary statistics for distributed coordination across brain states. Moreover, the contrast between rest and task points to distinct organizing principles. Resting-state dynamics primarily reflect intrinsic network architectures that are largely invariant to coordination strength, whereas task-evoked dynamics recruit flexible cortical coordination patterns that substantially reshape scaling relationships. Age-related effects predominantly impact task-evoked dynamics, suggesting that healthy aging preserves intrinsic network organization while compromising the adaptive coordination capacities required to meet external cognitive demands.

Synchrony-induced spatial scaling analysis yields quantitative biomarkers for conditions marked by disrupted neural coordination, as illustrated by our comparison of individuals with ADHD to healthy controls (*N* = 106; UCLA/CNP data), which revealed altered synchrony–scaling relationships (SI Fig. S23). This demonstrates that the framework can be extended to disorders characterized by atypical large-scale synchrony, including autism spectrum disorder, epilepsy, schizophrenia, and neurodegenerative diseases [King et al., 2018, Mathalon and Sohal, 2015, Ye et al., 2014], enabling quantification of how coordination deficits reshape spatial scaling properties and providing mechanistic insights beyond those afforded by conventional functional connectivity measures.

In summary, this work introduces a robust, data-driven generic framework for characterizing how coordinated dynamics shape universal scaling properties in complex systems such as the human brain. Age-related effects are shown to act primarily on synchrony-induced scaling during tasks, while leaving resting-state scaling relationships largely preserved. By explicitly removing marginal contributions, the framework isolates biophysical coordination mechanisms, positioning spatial scaling laws as a powerful tool for probing the multiscale organization of brain dynamics across the lifespan and in neurological or psychiatric conditions. Alternative synchrony metrics, such as the Kuramoto order parameter [Kuramoto, 1984] or Ising-spin–based [Weistuch et al., 2021] measures, may provide effective strategies to refine both theoretical predictions and empirical quantification in future studies. Beyond neuroscience, the proposed framework naturally extends to any complex system exhibiting scale-invariant spatial organization. The generalized RMS–variance formulation overcomes prior limitations by mitigating the confounding effects of temporal means and trends, thereby revealing coordination-driven scaling laws that would otherwise remain obscured. By stripping away marginal structure, it exposes fundamental biophysical (or mechanistic) principles governing collective dynamics, providing a unified analytical approach for measurements across ecological, financial, climate, and engineered systems.

### Methodology

#### Spatial scaling law

Our general methodology is elaborated here using a realistic example of spatiotemporal data from the brain. Consider spatio-temporal time series data *y*_*i*_(*t*), where *i* = 1, 2, …, *n*; *n* is the number of nodes or locations or brain regions, that represent the BOLD signals of four exemplary colored cortical regions (green, red, blue, and yellow) in Fig. 1. The multivariate data may follow any distribution (Poisson, negative binomial, gamma, uniform, and others). After a detrending operation, the additional trends and temporal mean are removed from such multivariate spatio-temporal time series. The spatial distribution of the detrended data *Y*_*i*_(*t*) across the entire location then follows a normal-like distribution, having negative to positive data spreading.

Exemplary values of the variable *Y*_*i*_(*t*_1_) at time *t*_1_, *Y*_*i*_(*t*_2_) at time *t*_2_ and so on, *Y*_*i*_(*t*_*T*_) at time *t*_*T*_ of the detrended data shown in Fig. 1 are taken from the four colored brain regions. The spatial variance *v*_1_ and spatial root-mean-square (RMS) *m*_1_ values are derived from spatial distributions over the entire space at each time instant. Next, the exponent *b* is obtained by fitting the least-squares linear regression to the log(m) scatterplot against log(v) at various time points, which here is referred to as the spatial scaling law (SL). Please refer to Fig. 1 for visual interpretation.

We assume that the detrended multivariate stochastic process represented by *Y*_*i*_(*t*) = (*Y*_1_(*t*), …, *Y*_*n*_(*t*)) is assumed to be stationary and ergodic [Brillinger, 2001, Reuman et al., 2017]. The spatial SL is then formulated as a linear relation between the spatial RMS (*m*(*t*)) and spatial variance (*v*(*t*)) after logarithmic transformation at time point *t*. The spatial RMS is defined as the square root of the quadratic mean over the entire locations (all brain regions) at time instant *t* as, 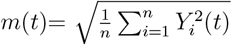. Thus, the spatial RMS considers squared values of all the data points directly, indicating the effective magnitude of the signal, averaged over the space at each time instant. The sample mean over the whole location at time *t* is calculated by 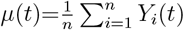, which measures the central tendency of the data at time *t*. The spatial variance at time *t* is usually expressed as 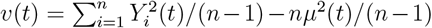, which measures the dispersion of data points around the sample mean. Here, the spatial variance (*v*) is expressed as a function of spatial RMS (*m*) as *v*(*t*)=*n* (*m*^2^(*t*) − *µ*^2^(*t*)) */*(*n* − 1). The scaler relationship between RMS and variance is defined by, *v*(*t*) = *am*^*b*^(*t*), or after a logarithmic transformation, we have,

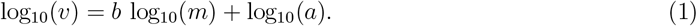

#### Coordinated dynamics quantified by synchrony (Ω)

Accordingly, the long-time correlation between measurements in any pair of locations is derived using pair-wise correlation coefficients, yielding a region-vs-region correlation matrix, as shown in Fig. 1. We derive average synchrony (Ω) from this correlation matrix to quantify brain-wide coordinated hemodynamics, by defining 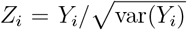, the standard measure of average synchrony [Reuman et al.,2017, Liebhold et al., 2004] is given by,

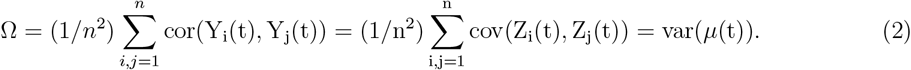

Detailed derivations are presented in SI. The synchrony Ω can be 0 ≤ Ω ≤ 1, which is a global metric capturing overall coordinated neural responses underlying interal and external stimuli driven brain activity from the pair-wise correlation matrix. Ω is 1 when the all regions are highly correlated, could be in- or, anti-phase correlation, and Ω=0 represents complete desynchrony across all regions. The pair-wise correlation matrix is also referred to as functional connectivity (FC) in neuroscience, [Friston, 2011, Bressler and Menon, 2010, Friston, 2003, Beckmann et al., 2005, Damoiseaux et al., 2006, Power et al., 2011, Yeo et al., 2011, Chakraborty et al., 2024], which varies from -1 to 1, where FC_*ij*_ = 1 implies complete coherence between *i*^th^ and *j*^th^ brain regions; FC_*ij*_ = −1 implies antisynchrony, and FC_*ij*_ = 0 depicts complete desynchrony between two brain regions. While FC can be quantified by correlation, covariance, or mutual information [Friston, 2011, Bullmore and Sporns, 2009, Bressler and Menon, 2010], it primarily serves as an empirical characterization of interregional spatial relationships without revealing the underlying mechanisms driving the covariation [Friston, 2011,Friston, 2003] .

#### Analytical formulation: Synchrony-induced spatial scaling law

The traditional way to test the scaling law is a linear fit between log(*m*(*t*)) and log(*v*(*t*)) for a finite realization of the processes. The linear regression exponent is *b* = cov(log(*m*), log(*v*))*/*var(log(*m*)) [Reuman et al., 2017]. Applying the delta method [Oehlert, 1992], we derive,

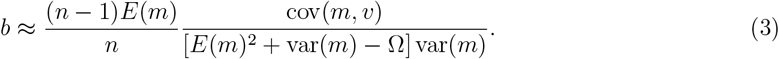

On the right-hand side, the first factor and the expected value *E*(*m*) depend on marginal distributions, *Y*_*i*_, but not on the correlations cor(*Y*_*i*_, *Y*_*j*_). Please refer to the SI for the detailed derivation of Eq. 3. This provides an idea behind our subsequent analyses: if the brain-wide coordinated activity measured by the average synchrony (Ω or var(*µ*)) changes, but the marginals *Y*_*i*_ remain fixed, we expect the slope *b* will change. We assume that cov(*m, v*) and var(*m*) have minimal effect on the slope. As synchrony (Ω) appears in the denominator of Eq. (3), the slope *b* is inversely proportional to the variations in synchrony, thereby encapsulating the long-term correlations across the spatial domain. In our empirical analysis, we restrict our examination to the linear association between synchrony (Ω) and the spatial SL exponent (*b*) captured by the association slope *β*, described in the section below.

In our empirical examples, we extracted the slope *b*_*syn*_ attributable to coordinated dynamics by removing the contribution of marginal distribution (*b*_*marg*_, driven by random processes) from the spatial SL exponent (*b*) using

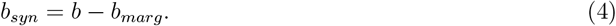

The slope *b*_*marg*_ is derived from the surrogate time series obtained after phase randomizing the empirical BOLD signal at each location; see SI “Section 6.3. Surrogate time series: phase-randomization.

#### Synchrony-scaling association (*β*)

Finally, we assessed the impact of synchrony on spatial scaling through the synchrony-scaling association slope (*β*), obtained by regressing synchrony strength (Ω) against the slope *b*_*syn*_, while removing the marginal contributions from random processes. We derive an approximate analytical relation between spatial SL and synchrony, then validate the framework using empirical fMRI data from healthy aging and developmental cohorts to demonstrate generalizability. The slope *β* is utilized to capture alterations in resting-state and task-evoked BOLD activity across brain regions and subnetworks along the lifespan trajectory. We have obtained the degree of impact of synchrony on the spatial SL using least-square linear regression fit between *b* and Ω. We obtained *β* using two methods: (i) Analytical approach: We analytically derived *b* from Eq. 3, calculated *b*_*syn*_ using Eq. 4, and then obtained *β* [Figs. 4(J,K) and 5(J)]. (ii) Fitting-based approach: We estimated *b* via least-squares linear regression of log_10_(*m*)-versus-log_10_(*v*), calculated *b*_*syn*_, and then obtained *β* [Figs. 4(L,M) and 5(K)].

#### Empirical Datasets

We used three independent cohorts from publicly available datasets. The Cambridge Centre for Aging and Neuroscience (CamCAN) *N* =645 healthy participants, http://www.mrc-cbu.cam.ac.uk/datasets/camcan [Taylor et al., 2017, Shafto et al., 2014]. Total *N* =89 healthy participants from the Nathan Kline Institute (NKI)/Rockland dataset, available in the UCLA Multimodal Connectivity Database (UMCD) [Brown et al., 2012]. We have randomly selected *N* =66 healthy controls (HC) and *N* =40 Attention-Deficit Hyperactivity Disorder (ADHD) participants from the Consortium for Neuropsychiatric Phenomics, https://f1000research.com/articles/6-1262/v2 [Poldrack et al., 2016].

#### Supporting Information Appendix (SI)

Supplemental Information (SI) contains definitions of all the metrics, analytical derivations, and algorithms to construct spatio-temporal time series data, numerical examples for different case studies using randomly drawn values from various distributions, results on empirical data (participant-wise, group-level), and the phase randomization process applied to the empirical BOLD signals. Full details of the analytic results are presented in SI, subsection S3. Full details of numerical simulations are provided in SI Appendix subsection S4, Simulation-based results

#### Code availability

All code for analyzing scaling laws and synchrony, estimating brain synchrony, and running the simulations used to generate the figures is available at https://github.com/ecesuman06/scaling-law-syn-aging

## Acknowledgement

We acknowledge online simulation support from the Neuroscience Gateway (NSG) [Sivagnanam et al., 2013]. We have no competing interests and did not use AI-assisted technologies in creating this work. AB and DR were supported by the Dementia Science Programme: Incidence/Prevalence/Risk/Intervention analysis of Dementia and Basic research by the Department of Biotechnology, Ministry of Science and Technology, Government of India, BT/HRD/Dementia/2017.

## Supplemental Material

### 1 Visualization of the synchrony-induced scaling law

Lets consider an arbitrary spatio-temporal time series data, *y*_*i*_(*t*), where, *i* = 1, 2, …, *N* ; *N* is the total number of nodes/sites/locations/brain areas. Figure S1 shows the schematic of the pipeline. The four colours represent resting state BOLD activity from four cortical regions. First, we apply detrending operation on the multivariate spatio-temporal time series. Thereafter, in the detrended data *Y*_*i*_(*t*), the additional trends and bias are removed, and temporal mean is set to zero. The bias can be described by taking an example from signal processing. Suppose, a DC voltage or a ramp like signal is added to a sinusoidal signal. Now, the signal starts oscillating around a positive voltage above the zero temporal mean. The detrending process removes the trend and DC bias. Thereafter, the signal starts oscillating around the zero mean voltage or zero temporal mean. We rounded up to 1e-6 decimal for zero value in numerical simulation. In second column in Fig. S1, the instantaneous values (*Y*_*i*_(*t*_1_) at time *t*_1_, or *Y*_*i*_(*t*_2_) at time *t*_2_ or *Y*_*i*_(*t*_*n*_) at time *t*_*n*_) of the detrended data from all the four brain regions are shown on the brain plot. The spatial variance (*v*_1_) and spatial root-mean-square (RMS, *m*_1_) value are then derived from the spatial distribution over the entire space for each time instant. Next, the exponent *b* is obtained from the log-log relationship between the RMS and variance, which is referred to as spatial scaling law (spatial SL). We derive the long-time correlation using Pearson correlation coefficient between any pair of regions, which gives a region-vs-region (space-vs-space) square matrix. Averaging over the absolute values of the entire matrix the spatial synchrony (Ω) is obtained. Finally, we examine the impact of synchrony on the spatial SL exponent from a linear fitting using common least square regression, considering Ω and *b* as independent and dependent variables, respectively. From the regression model, we estimate the slope (*β*), that implies the degree of influence of synchrony on the SL. In addition, we analytically derive the approximate formula for the spatial SL as a function of synchrony. We have tested the method on empirical examples using fMRI data from CamCAN healthy aging cohorts; see Empirical results section. The metric *β* is used to examine the changes in rest and task evoked BOLD activity across the adulthood aging trajectory.

**Figure S1.**
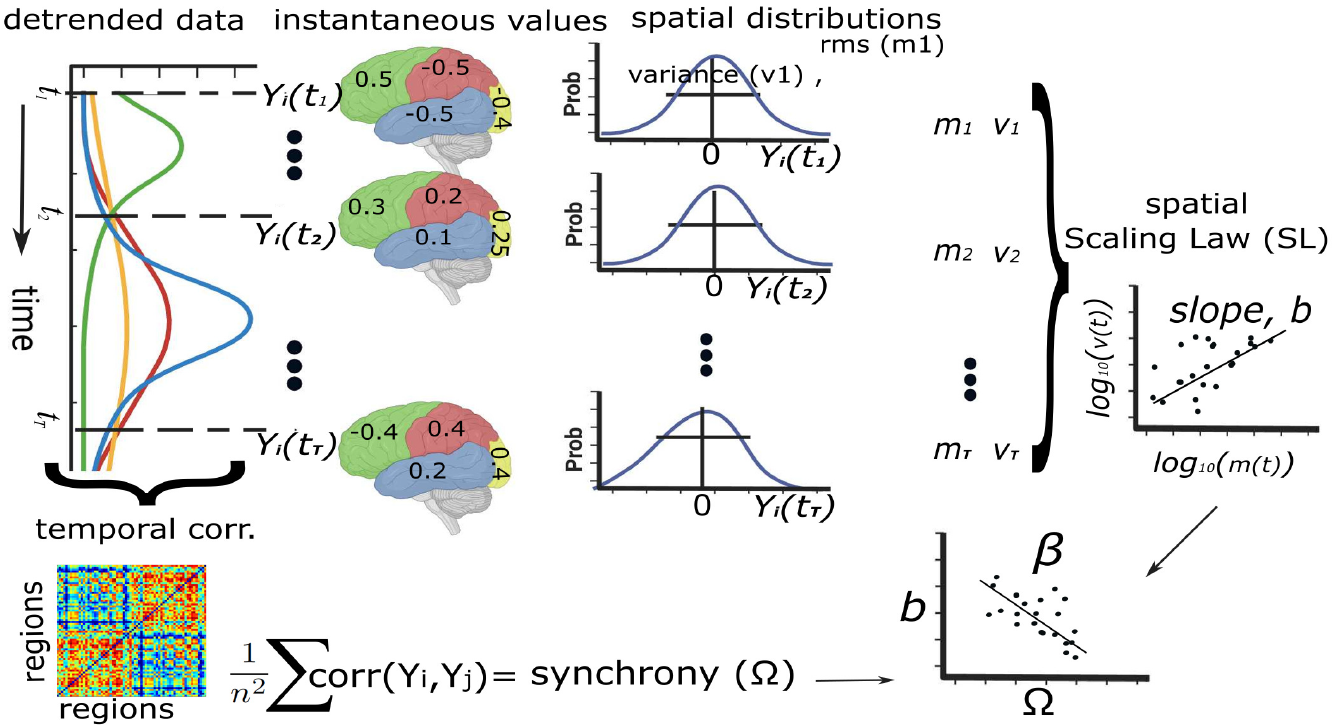
Schematic diagram of the proposed pipeline has been illustrated using a BOLD series from a participant. First step of our method involves, detrending process, where we remove the trends and the temporal mean is set to zero. Next, the amplitude of the detrended BOLD signal of all the brain regions are squared and summed at a time instant *t* to obtain spatial RMS at time *t*, similarly, we store spatial variance at each time point. The exponent (*b*) of the least-square linear fit between the spatial RMS (log_10_(*v*)) and variance (log_10_(*v*)) is derived, besides the average spatial synchrony (Ω) is also calculated for each participants. Finally, we make a scatterplot to examine the relationship between synchrony (Ω) and SL exponent (*b*) using a linear regression model. The slope *β* and *p*-value describe the influence of the synchrony on the emergence of the spatial SL in both the empirical and numerical examples.

### 2 Definitions

Suppose the amplitude (e.g., population density, voltage, amplitude of blood-oxygen level dependent activity) of the spatio-temporal time series data in location *i* at time *t* is modeled by the random variable *Y*_*i*_(*t*), *i* = 1, …, *n* and *n* = total number of sites (e.g., nodes in a network, number of patches in ecological network, number of brain areas or region of interests). We assume that the multivariate stochastic process *Y* (*t*) is stationary and ergodic [1, 2].

A wide range of stochastic processes have the property that their average statistical properties are independent of where they are formed along the time [3]. For a stationary signal, the basic signal properties of mean and variance do not change over time [4]. Roughly speaking, a “stationary” process is one whose joint probability distribution is not affected by shifts in time, so that mechanisms of the dynamics are not shifting in time [2]. For such a signal, the measurement of the mean or variance over only one segment is sufficient to estimate the signal’s true mean [3, 4]. An “ergodic” process is one for which statistical characteristics can be deduced from a single, sufficiently long realization of the process, so that, for instant, long-term statistical outcomes are independent of initial conditions [2]. Ergodicity is a more restrictive concept compared with that of wide-sense stationarity, or even of stationarity in the strict sense [5]. More precisely, for an ergodic process, time averages coincide with ensemble averages [5]. In neurophysiological experiments, the validity of the ergodicity assumption, i.e., replacing ensemble means by time means, is often implicitly assumed [3]. Precise definitions of stationary and ergodic processes can be found in Ref. [1]. The definition of the spatial RMS, sample mean, spatial variance and synchrony of a given multivariate time series data are given below.

#### 2.1 Temporal mean (*µ*_*t*_)

Temporal mean [6] for a given raw data, i.e., spatio-temporal time series, *Y*_*i*_(*t*) is computed for individual site *i*, as

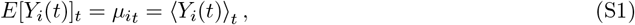

where, *i* represents index of the space. ⟨.⟩ denotes long-time average, or average over one cycle when *Y*_*i*_(*t*) is periodic. After detrend operation, the temporal mean *µ*_*t*_ becomes 0.

#### 2.2 Temporal variance (*v*_*t*_)

Temporal variance [2] of location *i* over the entire time length is expressed as,

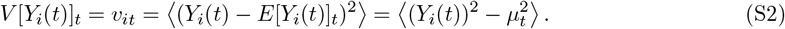

This spatial quantity measures dispersion or spread of the data points around the temporal mean (*µ*_*t*_).

#### 2.3 Spatial mean (*µ*)

The arithmetic mean or sample mean over all regions at time *t* is calculated by

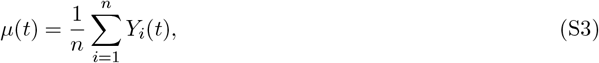

which measures the central tendency of the sample data at time *t*. The spatial mean contains negative and positive values *µ* ∈ [−1, 1], where var(*µ*)*>*0, and E(*µ*)= 0.

#### 2.4 Spatial variance (*v*)

Spatial variance [2] over the space at a time instant *t* is expressed by,

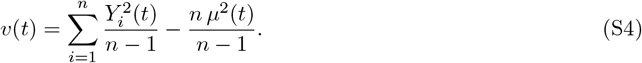

This spatial quantity measures dispersion or spread of the data points around the sample/ensemble/spatial mean (*µ*).

#### 2.5 Spatial RMS (*m*)

The spatial root-mean-square (RMS) is defined as square root of the quadratic mean, i.e., RMS over the entire space (e.g., over all brain regions) at time *t* as,

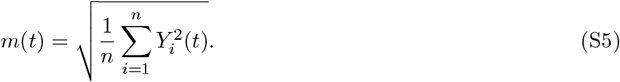

Thus, the spatial RMS of the signals takes into account the squared values of all the data points directly, indicating effective magnitude of the signal, averaged across space at each time instant. The values of *m* are always positive, i.e., *m >* 0.

#### 2.6 Average Synchrony (Ω)

The term ‘synchrony’, used in this study, is referred to as a global metric that quantifies the strength of coordinated cortical activity or brain-wide coactivation pattern. To define the metric synchrony, we have first selected a very fundamental measure, functional connectivity (FC), frequently used in neuroscience to study resting state fMRI data. The FC matrix is computed using Pearson correlation coefficient that captures the long-term correlation or degree of coherence between two BOLD signals from each pair of brain regions. We get a symmetric square matrix of size=regions X regions, where the off-diagonal entries can vary between [-1,1]. Finally, we averaged the entire matrix which is referred to as the spatial synchrony (Ω), that measures the strength of coherence in BOLD activity of individual adults. The standard measure of average spatial synchrony is also used earlier in ecological population [2, 7]. Expression for the global metric synchrony is,

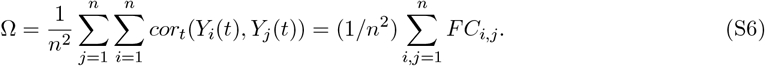

where, *FC*_*i,j*_ = *cor*(*Y*_*i*_, *Y*_*j*_). *FC* is the static functional connectivity matrix with a size (*n*×*n*), measured by Pearson correlation coefficient between the time series of a pair of locations. *n* is the number of locations, sites, space, ensembles, nodes in a network, or, brain regions. For example, *n* is the total number of brain regions for a given brain atlas, such as, automated anatomical labelling (AAL) atlas [8], where *n* = 116.

### 3 Analytical results

#### 3.1 Mathematical derivation of the spatial scaling exponent

Lets assume, the amplitude of a spatio-temporal time series data at time *t* in *i*^*th*^ location is modeled by a real valued random variable *Y*_*i*_(*t*) for *i* = 1, …, *n*, where *Y* (*t*) = (*Y*_1_, …, *Y*_*n*_) is stationary and ergodic. We are interested in the log(*v*)-log(*m*) scatter plot for a finite duration realization of these processes (ignoring the times *t*, when *m*(*t*) or *v*(*t*) was zero), and particularly in the slope and intercept of an ordinary linear regression through such a plot,

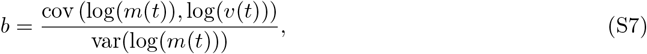

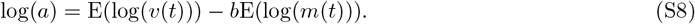

These are the two basic quantities in the spatial scaling law (SL). The covariance, variance, and RMS are sample quantities computed over time for the finite-duration realization. Given the ergodicity of *Y* and consistency of the spatial RMS, mean, variance and covariance, we consider the previous equations as a general form in the sequel for large samples, valid for both temporal and spatial cases. We compute the covariance, variance, RMS, and expected values for the marginal distributions of the processes *m* and *v* when these quantities are positive, and likewise we get the same results from the positive- and negative-valued data distributions. Furthermore, we assume that both the scaling parameters (*m* and *v*) are positive and finite.

Suppose the long-term temporal correlations *ρ*_*ij*_ = *cor*(*Y*_*i*_, *Y*_*j*_) exist for all *i, j* = 1, …, *n*. These temporal correlations between a pair of sites represent spatial synchrony between the regions *i* and *j* (again making use of ergodicity, and consistency of the sample covariance and variance). The synchrony, 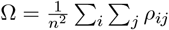 is an average correlation across all pairs of sampling locations (including comparisons of a location with itself, which give *ρ*_*ij*_ = 1, and is commonly used in empirical studies. We will analyze the influence of Ω on *b*. Ω is naturally interpretable as a measure of average coherence in anti- and in-phase state for the following additional reasons. First, letting, 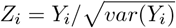, we have

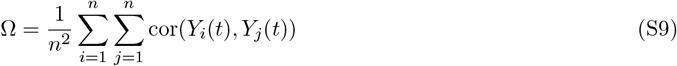

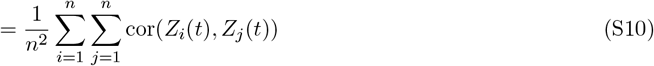

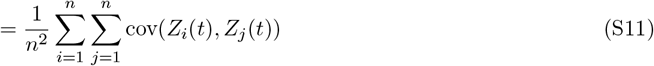

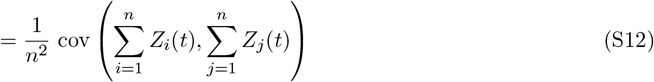

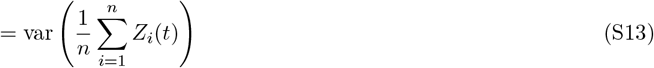

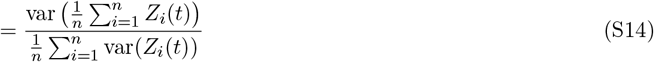

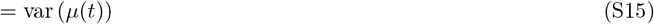

where, equation (S15) is the variance of the spatial mean *µ*, and is readily interpretable as the extent to which oscillations in local time series reinforce each other or cancel in the average time series. In all our examples for the detrended data, var(*Z*_*i*_(*t*)) = 1.

Second, Eq. (S15) and the definition of Ω show that 0 ≤ Ω ≤ 1. The value 0 occurs when the variability in the *Z*_*i*_ exactly cancels, i.e., 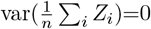 and perfect asynchrony. The value 1 occurs for perfect synchrony, when *Z*_*i*_ = *Z*_*j*_ for all *i, j*. An alternative measure for synchrony is the summation of 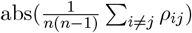. The correlation coefficient *ρ* can take negative values. In case of perfect anti-synchrony and synchrony, the Ω can be 1, except, for perfect asynchrony.

To understand the relationship between synchrony and scaling law for large samples under assumptions of stationarity and ergodicity, it is sufficient to work with the marginal distribution *Y*, which follow a normal distribution in our case, instead of with the process *Y* (*t*). We therefore consider families of distributions of the form *Y* = (*Y*_1_, …, *Y*_*n*_) such that for each *i*, all members of the family have the same marginals *Y*_*i*_, and we consider how Ω and *b* covary through the family before and after the detrending operation. After detrending operation, the underlying distribution of the marginal started following Gaussian normal distribution. In the theoretical development, marginals are kept constant to understand the influence of synchrony Ω in isolation from other potential influences on *b*. To describe the intuition of the mathematical analyses that will be described in details in the next section, we need a lemma, taken directly from [2]. They cited [9, 10], and [11] pp. 355-358.

**Lemma 1**. Let *x* be a real valued random variable with finite mean E(*x*) and variance var(*x*). If a real-valued function *f* of real *x* is twice differentiable at *E*(*x*), then the *delta method* [10] gives

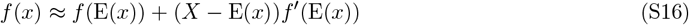

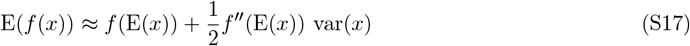

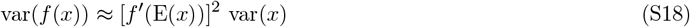

We now use the Lemma 1 with the real-valued function *f* → log, and for two random variables, *x* → *m, v* (RMS and variance), which are ergodic stationary stochastic processes.

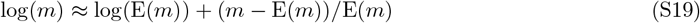

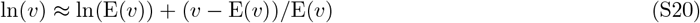

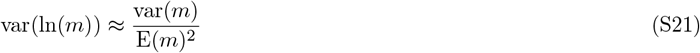

where the approximation is good as long as *m* and *v* do not vary too much around *E*(*m*) and *E*(*v*), relative to the range of *x* for which first- and second-order Taylor expansions of the log(*x*) at *E*(*m*) and *E*(*v*) are good approximation. Therefore,

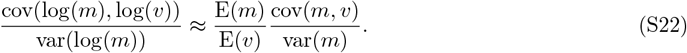

Now, using Eqs. (S19) and (S20), we can write

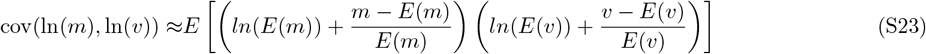

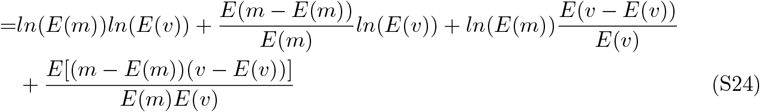

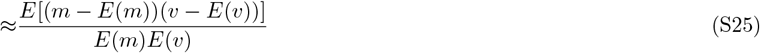

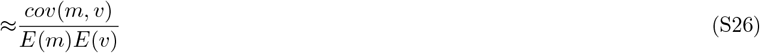

Using equation Eqs. (S21) and (S26) we have,

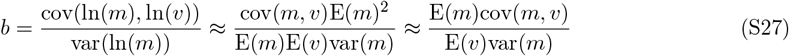

Now, var(*m*) = E(*m*^2^) − E(*m*)^2^, or, E(*m*^2^) = var(*m*) + E(*m*)^2^, and var(*µ*) = E(*µ*^2^) − E(*µ*)^2^ or, E(*µ*^2^) = var(*µ*) + E(*µ*) .

The RMS is defined as, 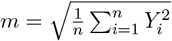 or 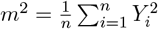 . The variance is defined as, 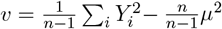, or, 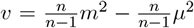. Taking expectation to the both side, 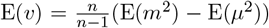. Now we can rewrite,

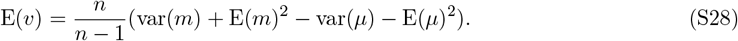

Next, putting the results from Eq. (S28) into Eq. (S27), we can write,

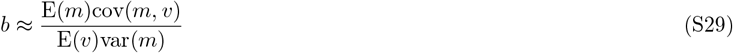

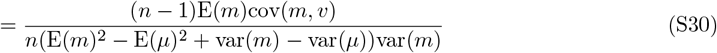

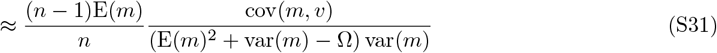

The first factor on the right-hand side of Eq. (S31) solely depends on the number of location or brain areas, *n*, and the marginals *Y*_*i*_, but not on their correlations. The quantities E(*m*) and var(*m*) also depend only on marginal distribution structure. The quantity var(*µ*), equals to 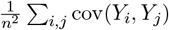, is directly related and similar in form to the measure of coherence, i.e., the average spatial synchrony Ω. Therefore, Eq. (S31) provides the intuition behind our analysis: only if synchrony (Ω or var(*µ*)) changes but the marginal *Y*_*i*_ remain fixed, the slope *b* will be affected. In other words, if the long-term correlation among the spatio-temporal time series data varies, but the complexity in the time series remain fixed, the slope *b* will be altered. From a logical assumption, the time series complexity [12] can be represented by a specific dynamical nature, such as periodic, two periodic, quasiperiodic, chaotic and stochastic.

Since, the synchrony Ω is in the denominator, the slope *b* is inversely proportional to the changes in the synchrony, which captures long-term correlations across the entire space.

#### 3.2 Linear Fitting

Dropping the time dimension, the spatial RMS at time *t* is expressed as,

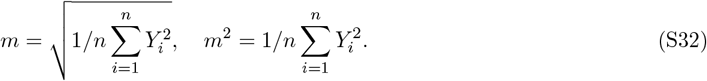

Sample mean at time *t* is 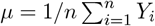.

Spatial variance at time *t* is,

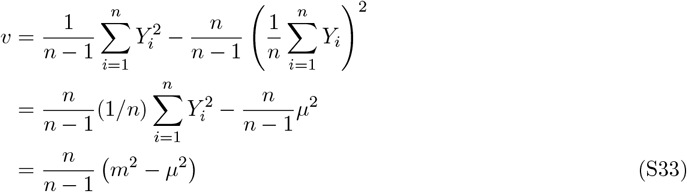

Taking ‘log’ to both sides of Eq. S33, we get,

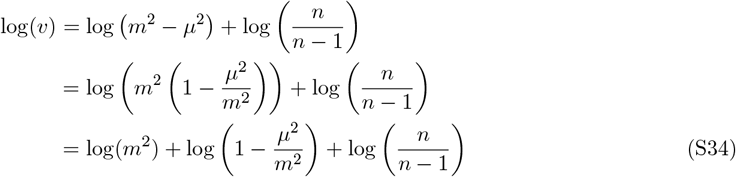

Detrending sets the temporal mean to zero, *µ*_*t*_ = 0; though, this can take any value in natural or physical processes prior to detrending. Thus, the data tends to follow Gaussian distribution, and the second term in Eq. (S34) will be

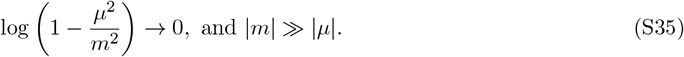

Eq. (S34) can be written as,

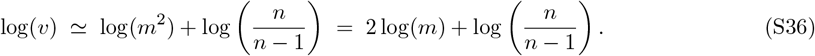

From the log-log fitting between the two quantities *m* and *v*, in a normally distributed data, the SL exponent is found as, *b* = 2. Thereby, we considered normal distribution as a baseline distribution corresponding to the contribution coming from data marginals, which is further utilized to remove marginal effect on the measurement in empirical data analysis.

The SL slope *b* was computed through ordinary linear regression of log(*v*) against log(*m*) using most common least-square fit. In this study, we used the least-square regression in all our linear fittings using MATLAB function ‘fitlm’, and Python function ‘stats.linregress’. For each block, residuals of the log(*v*)- versus-log(*m*) regression were computed and used to generate a root mean squared error (RMSE). The mean (across block) RMSE was plotted against average synchrony Ω.

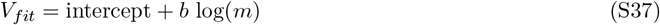

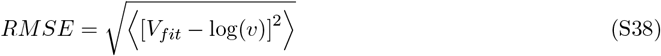

### 4 Simulation-based results

The multivariate time series is first generated by randomly drawn values from binomial distributions. The data size is *T* × *n*, where *T* and *n* represent total time points and total number of spaces, respectively. The space can be considered as total number of brain regions or total number of nodes in a network. We repeatedly generated the time series for 645 independent realizations for each distribution type.

#### 4.1 Effect of non-zero temporal mean on spatial scaling exponent: Independent and identically distributed

We have considered two cases, e.g., without and with detrending operations. We have detrended the multivariate time series. Detrending removes the additional bias from each time series data by setting temporal mean to zero. The detrended data now follow normal distribution. The spatial scaling slope is observed as *b* ≃ 2 in all experimental distributions after detrending.

##### 4.1.1 Independent and identically distributed multivariate data: Poisson distribution

Figure S2 shows numerical test on a spatio-temporal time series, size 260×116 (time points are 260 and number of locations are 116) has been generated by randomly drawing values from Poisson distribution with *λ*=50. We first generated a spatio-temporal data, histogram in red is plotted for case 1, the temporal mean at 50. Next, we have shifted the mean of the entire data, a constant value 20 to/from all is added or subtracted (temporal mean=70 for case 2 in black, and temporal mean =30 for case 3 in green) to the temporal of the original data (red) while freezing the variance and auto-correlation. Different random seeds are selected to construct time series for each locations. In all test cases we have considered an average correlation, between any pair of locations, close to zero, i.e., identical and independently distributed through time at each location.

**Figure S2.**
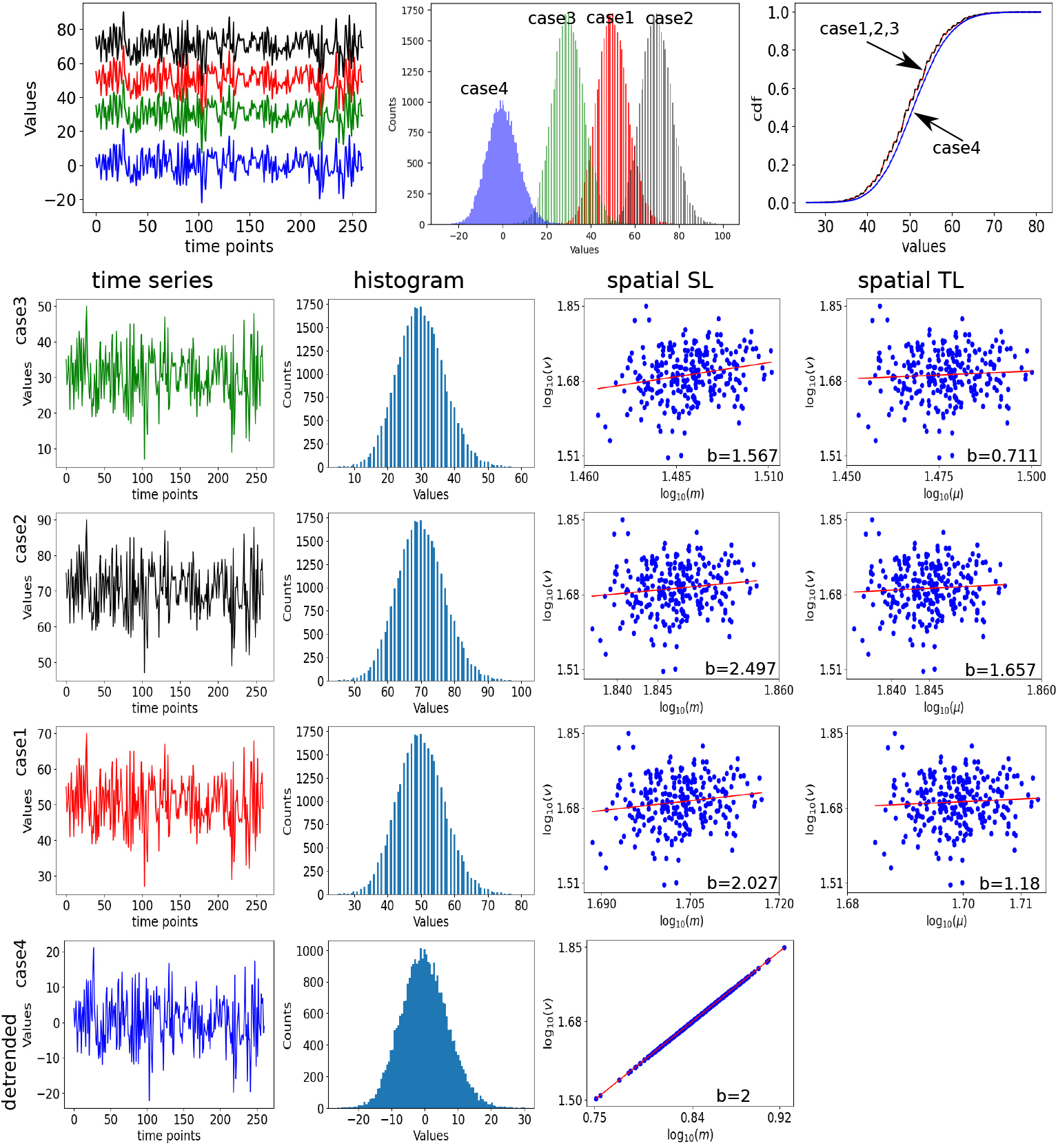
Test 1 for Poisson distribution. We have used Python function “numpy.random.poisson(50, size= (261, 116))” to create the spatio-temporal time series data. The seeds are varied randomly for different locations. Long-term correlation between any pair of locations are almost zero. Next, shifted the temporal mean of the entire data by adding and subtracting 50 from the previous case example to create case 2 (black) and case 3 (green) examples. The detrended scenario, labeled as case 4, shown in blue at the bottom row.

We have plotted single time series out of 116 locations for three different cases 1, 2, 3 in red, black and green, respectively. Time series, histogram plot of the data distribution and cumulative distributions are shown in top row in Fig. S2, respectively, arranged from left to right order. All the three cases 1, 2 and 3 are again plotted separately in fourth, third and second rows, respectively in Fig. S2. The spatial SL slope for case 1,2 and 3 are *b*=2.027, 2.497, 1.567, and the spatial TL slopes are *b*=1.18, 1.657, 0.711.

After detrending, the temporal mean is set to zero in all three cases. Thereafter, the sptio-temporal time series no longer follow Poisson distribution. The time series, histogram of the data distribution and cumulative distribution are visualized in blue referred as case 4 in top row in Fig. S2, and also separately plotted in bottom row. After detrending, the data tends to follow normal distribution, as seen in the cumulative distribution plot in blue for case 4. The spatial SL slope is found to be *b*=2 shownn in the bottom row in Fig. S2.

We have repeated the similar steps to generate the plots in Fig. S3 under the same statistical settings for Poisson (*λ* = 30, 50, 70) for another realization and derive the spatial SL and TL slopes. Compare to previous example shown in Fig. S2, for this test model, another realization of the time series are taken into consideration, when other properties are not disturbed. The average spatial correlation is close to zero. The spatial SL slope for case 1, 2 and 3 (red) are *b*=1.497, 1.729, 1.278 and the spatial TL slopes are *b*=0.593, 0.831, 0.355. We observed different SL exponent values because of the shifts in only the mean values of the distributions, which eventually alter the SL and TL slopes. These are different compared to the previous observations shown in Fig. S2. Next, we detrend the multivariate data, which sets temporal mean to zero in all the three cases. The time series of one location, the data distribution and the cumulative distribution are visualized in blue referred as case 4 in top row, in Fig. S3, the time series and distribution of the detrended data are plotted in bottom row first and second column, respectively, in Fig. S3. In the detrended cases, the spatial SL slope is found *b*=2.001 shown in bottom row third column in Fig. S3.

**Figure S3.**
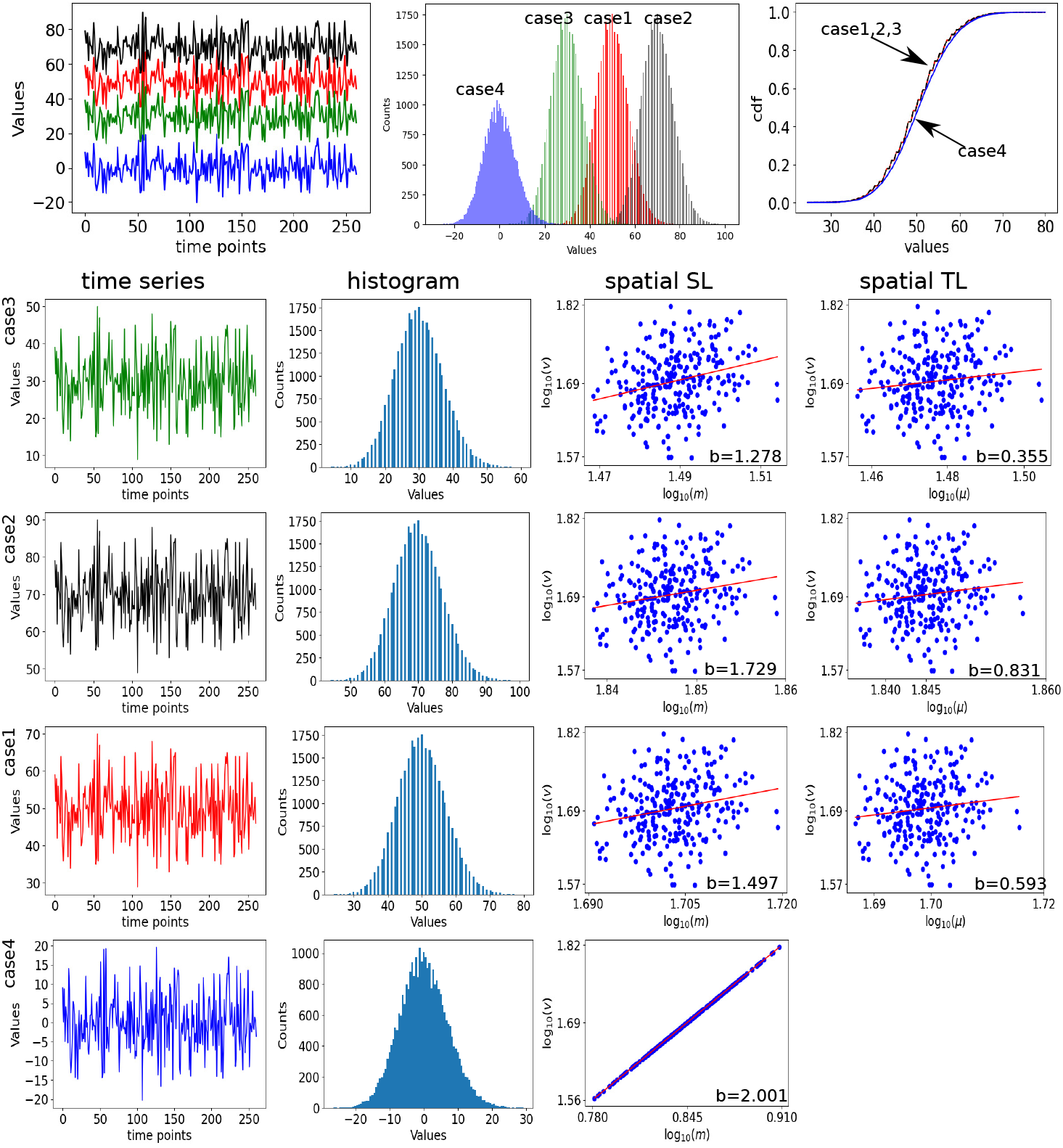
Test 2 for Poisson distribution. We have used Python function “numpy.random.poisson(50, size= (261, 116))” to create the spatio-temporal time series data shown in red. Next, shifted the temporal mean of the entire data by adding and subtracting 50 from the previous case example to create case 2 (black) and case 3 (green) examples. The detrended scenario, labeled as case 4, shown in blue at the bottom row.

##### 4.1.2 Independent and identically distributed data: Binomial distributions

Figure S4 shows numerical test on a spatio-temporal time series, size 260×116 (time points are 260 and number of locations are 116) has been generated by randomly drawing values from Binomial distribution with trials *n* = 100 and probability *p* = 0.5. We first generated a spatio-temporal data, histogram in red is plotted for case 1, the temporal mean at 125. Next, we have shifted the mean of the entire data, a constant value 50 and 100 are added to the temporal of the original time series data (red) while freezing the variance and auto-correlation, shown in black and green (case 2 and case 3). Different random seeds are selected to construct time series for each locations. In all test cases we have considered an average correlation, between any pair of locations, close to zero, i.e., identical and independently distributed through time at each location.

**Figure S4.**
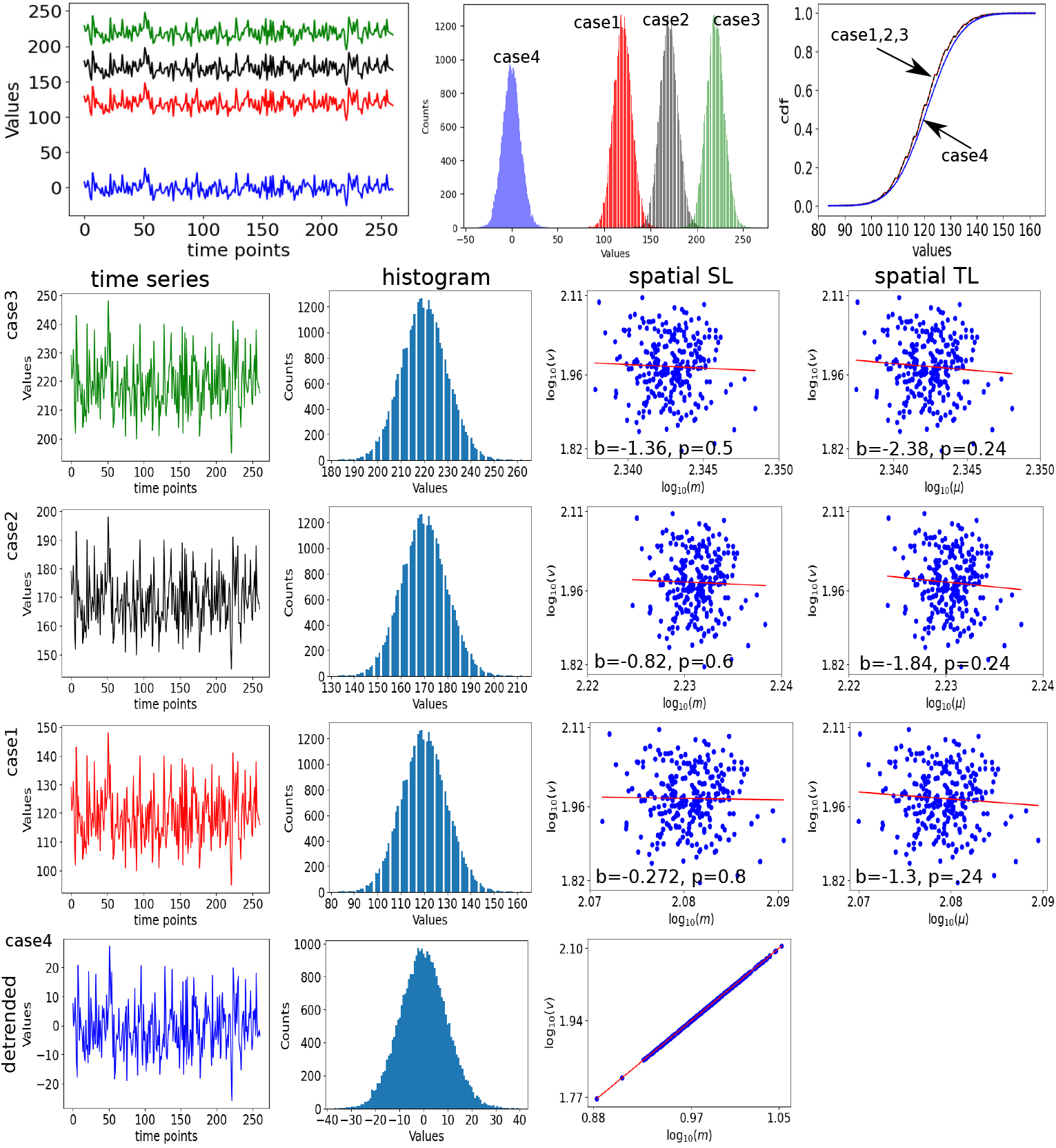
Test 1 for Binomial distribution. The Python function ‘numpy.random.binomial(*n*=100, *p*=0.5, size= (261, 116))’ has been used to construct the time series. The first distribution created with the temporal mean at 125, shown in red labeled as case 1. Next, we have shifted the mean of the entire data, a constant value 50 and 100 are added to the temporal of the original time series data (red) while freezing the variance and auto-correlation, shown in black and green (case 2 and case 3). Detrended data analysis is marked as case 4, shown in blue at the bottom row.

We have plotted single time series out of 116 locations, in green, red and black colors for three different cases, such as, temporal mean=125, 175 and 225 labeled as case 1, 2, 3, and corresponding colors are red, black and green, respectively. Time series, data distribution and cumulative distributions are shown in Fig. S4, respectively plotted from left to right order in top row. All the three cases 1, 2 and 3 are separately plotted in fourth, third and second rows, respectively in Fig. S4. The spatial SL slope for case 1,2 and 3 are *b*= -0.272, -0.82, -1.36, and the spatial TL slopes are *b*=-1.3, -1.84, -2.38.

After detrending, the temporal mean is set to zero in all three cases. Thereafter, the sptio-temporal time series no longer follow Poisson distribution. The time series, histogram of the data distribution and cumulative distribution are visualized in blue referred as case 4 in top row in Fig. S4, and also separately plotted in bottom row. After detrending, the data tends to follow normal distribution, as seen in the cumulative distribution plot in blue for case 4. The spatial SL slope is found to be *b*=2 shownn in the bottom row in Fig. S4.

We have repeated the similar steps followed in Fig. S4 and generated the plots in Fig. S5 under similar statistical settings for Binomial distribution (temporal mean=125, 175 and 225) for another realization and derive the spatial SL and TL slopes. Compare to previous example shown in Fig. S4, for this second test in Fig. S5, only time series fluctuations are different, but other properties are not disturbed. The average spatial correlation is close to zero. The spatial SL slope for case 1, 2 and 3 are *b*=4, 5.31, 6.62, 5.31 and the spatial TL slopes are *b*= 3.13, 4.43, 5.74. We observed different SL exponent values because of the shifts to the mean values of the distributions for three cases 1,2 and 3, which eventually alter the SL and TL slopes, as seen in Fig. S5. Next, we detrend the multivariate data, which sets temporal mean to zero in all the three cases. The time series of one location, the data distribution and the cumulative distribution are visualized in blue referred as case 4 in top row, in Fig. S5, the time series and distribution of the detrended data are plotted in bottom row first and second column, respectively, in Fig. S5. In the detrended cases, the spatial SL slope is found *b*=2.001 shown in bottom row third column in Fig. S5.

**Figure S5.**
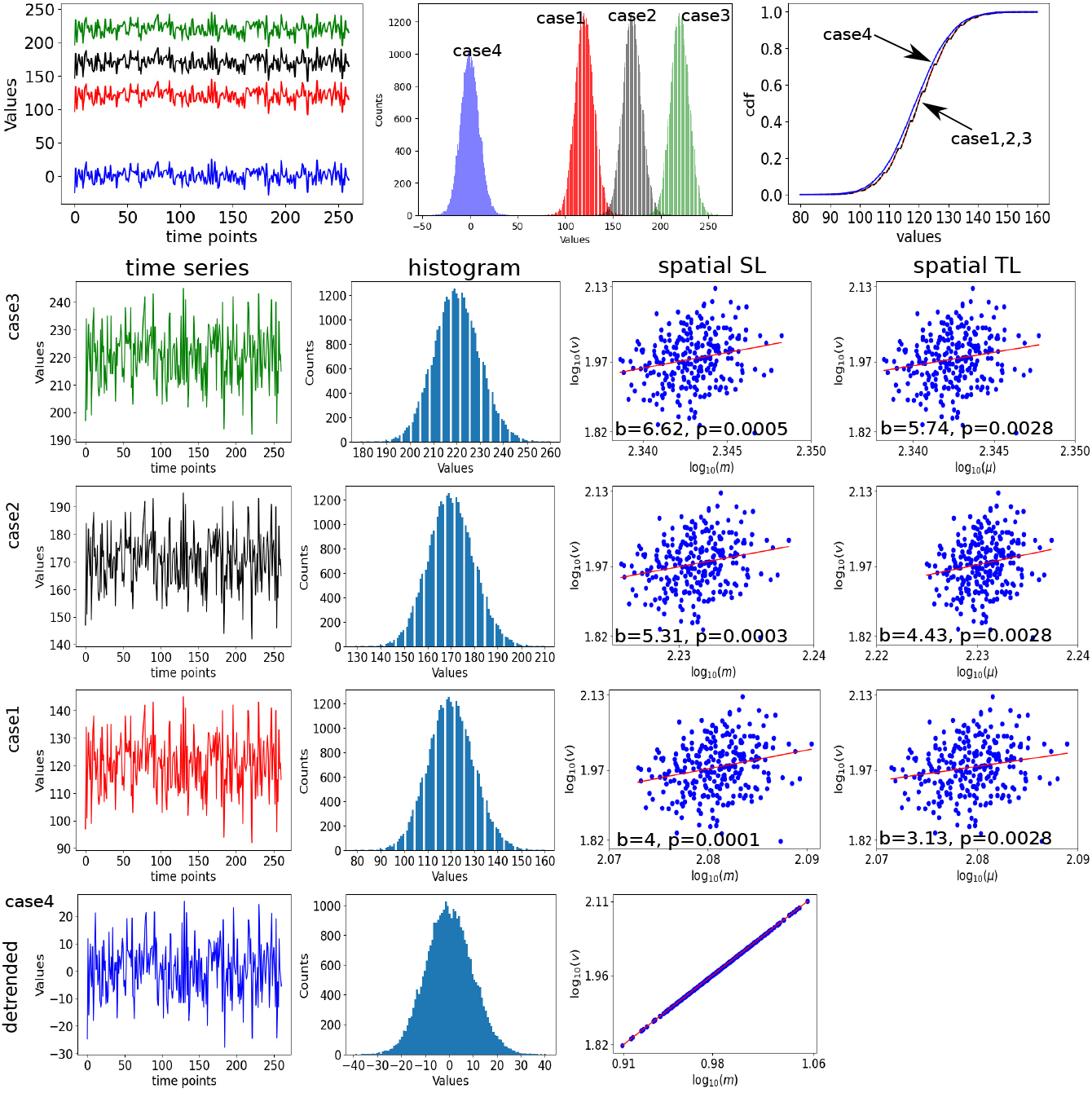
Test 2 for Binomial distribution. The Python function ‘numpy.random.binomial(*n*=100, *p*=0.5, size=(261, 116))’ has been used to construct the time series.

We have repeated the steps followed in previous two cases Fig. S4 and S5 to generated the plots in Fig. S6 for negative Binomial distribution (temporal mean=0.1, 0.6 and 1.1, labeled as case1, 2 and 3, corresponding colors are green, red and balck) and derive the spatial SL and TL slopes in three cases. The spatio-temporal time series constructed for case 2 (red) and case 3 (black) are generated by adding constant values 0.5 and 1 to the temporal mean of case 1 (green). In this test example, the average spatial correlation is close to zero. The spatial SL slope for case 1, 2 and 3 are *b*=1.28, 4.23, 2.66, and the spatial TL slopes are *b*=0.314, 3.47, 1.89 as seen in Fig. S6. We observed different SL exponent values because of the shifts to the mean values of the distributions for three cases 1,2 and 3, which eventually alter the SL and TL slopes, as seen in Fig. S6. Next, we detrend the spatio-temporal data, which sets temporal mean to zero in all the three cases. The time series of one location, the data distribution and the cumulative distribution are visualized in blue referred as case 4 in top row, in Fig. S6, the time series and distribution of the detrended data are plotted in bottom row first and second column, respectively, in Fig. S6. In the detrended cases, the spatial SL slope is found *b*=2 shown in bottom row third column in Fig. S6.

**Figure S6.**
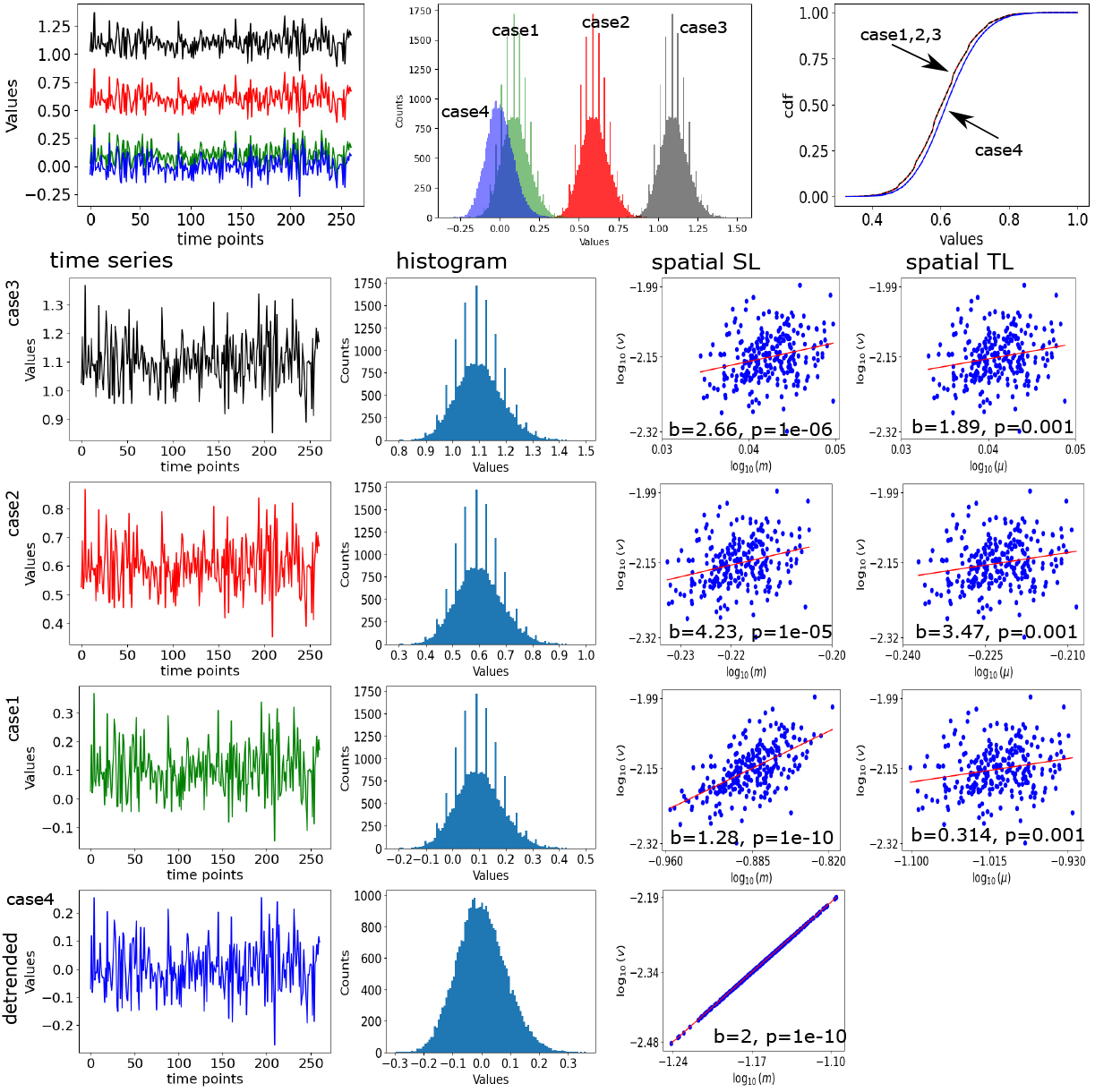
Test 3 for Negative Binomial distribution. The Python function ‘numpy.random.negative.binomial(n=100, p=0.5, size=(261, 116))’ has been used to construct the time series, and normalized by the maximum amplitude of each location.

##### 4.1.3 Independent and identically distributed data: Uniform distribution

Figure S7 shows numerical test on a spatio-temporal time series, size 260×116 (time points are 260 and number of locations are 116) has been generated by randomly drawing values from Uniform distribution. We first create a spatio-temporal data having the temporal mean at 0.5. The time series, the histogram and cumulative distributions are plotted in red marked as case 1. Next, we have shifted the mean of the entire data, where constant values 2 and 4 are added (temporal mean=2.5 for case 2 in black, and temporal mean =4.5 for case 3 in green) to the temporal of the original data (red) while freezing the variance and auto-correlation. Different random seeds are selected to construct time series for each locations. In all test cases we have considered an average correlation, between any pair of locations, close to zero, i.e., identical and independently distributed through time at each location.

**Figure S7.**
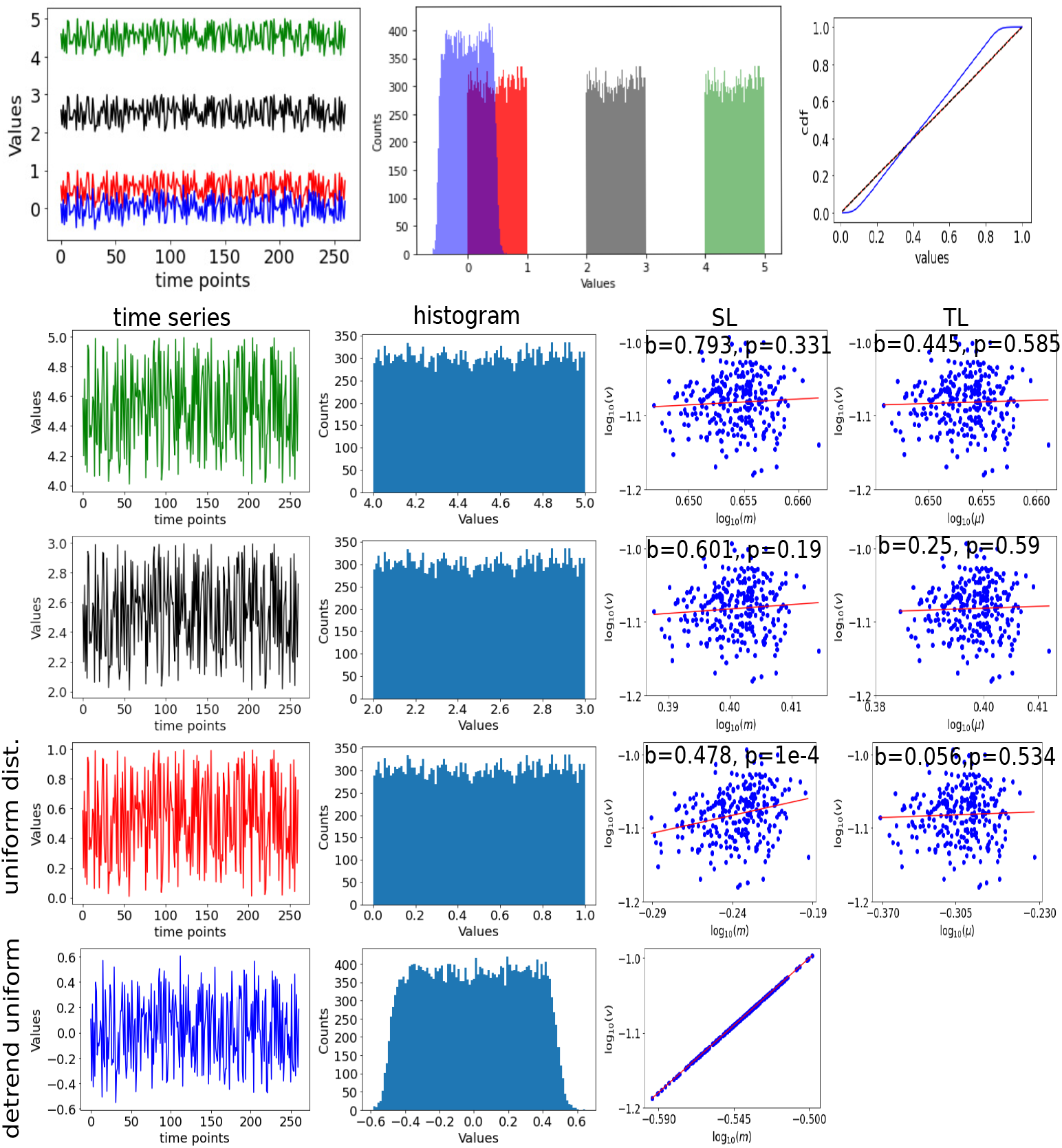
Uniform distribution. We have used Python built-in function ‘random.uniform(0,1,[261,116])’ to create the spatio-temporal data following uniform distribution, where pair-wise correlation between two locations are near zero. The temporal mean is 0.5 labeled as red. The constant values 2 and 4 are added to shift the temporal of the entire time series and generate two cases 2 and 3, corresponding colors are black and green. Detrended case is shown in blue in bottom row.

We have plotted single time series out of 116 locations for three different cases 1, 2, 3 in red, black and green, respectively. Time series, histogram plot of the data distribution and cumulative distributions are shown in top row in Fig. S7, respectively, arranged from left to right order. All the three cases 1, 2 and 3 are again plotted separately in fourth, third and second rows, respectively in Fig. S7. The spatial SL slope for case 1,2 and 3 are *b*=0.478, 0.601, 0.793, and the spatial TL slopes are *b*=0.056, 0.25, 0.445. The SL and TL exponents are not significantly emerged in these three cases.

After detrending, the temporal mean is set to zero in all three cases. Thereafter, the sptio-temporal time series no longer follow Poisson distribution. The time series, histogram of the data distribution and cumulative distribution are visualized in blue referred as case 4 in top row in Fig. S7, and also separately plotted in bottom row. After detrending, the data tends to follow normal distribution, as seen in the cumulative distribution plot in blue for case 4. The spatial SL slope is found to be *b*=2 shownn in the bottom row in Fig. S7.

##### 4.1.4 Independent and identically distributed multivariate data: Normal distribution

#### 4.2 Summary: Independent and identically distributed data

After detrending operation, we have observed that the data tends to follow normal distribution around zero mean. We have now conducted numerical tests on the spatio-temporal time series data constructed by randomly drawn values from a normal distribution. In the independent and identically distributed multivariate data (temporal mean *µ* = 0, and variance=1), we calculated the spatial RMS and spatial variance *v* and presented in Fig. S9. We considered Gaussian normal distribution to generate multivariate time series data of size, Time × locations (260 x 116). Figure S9(a) shows histgram of the data distribution. Log-normal distributions of the spatial RMS and variance are shown in Figs. S9(b) and (c), respectively. Exponent of the log-log fitting between spatial RMS and variance is found as *b*=1.988 (using analytical expression, *b*=2.002). (e) The exponents are estimated setting different random seeds for 645 independent observations. We generate total 645 spatio-temporal data to check the association between synchrony (Ω) and the exponent (*b*), when the pair-wise correlation between any two locations are near zero. We observe that synchrony does not influence the scaling exponent in independent and identically distributed normal distributions. The Sl exponent close to *b* = 2, is emerged, only when the temporal mean is set to zero and the average spatial synchrony is very weak. Irrespective of the original distribution in the multivariate data, after detrending the spatial SL exponent is always found near 2, when synchrony is very weak and underlying mechanism of origin is a random process. We consider the normal distribution as a null model, and is responsible for data marginal contribution for random processes in the detrended data. Under this background, we assume that if the neural activity is driven by the physical properties of given external stimuli, or spontaneous activity driven by internal stimuli, the scaling law may be influenced by synchrony differently in different brain states, rather than impacted by data marginals’ distribution structures. Thus, we can separate the two effects from our measurements.

**Figure S8.**
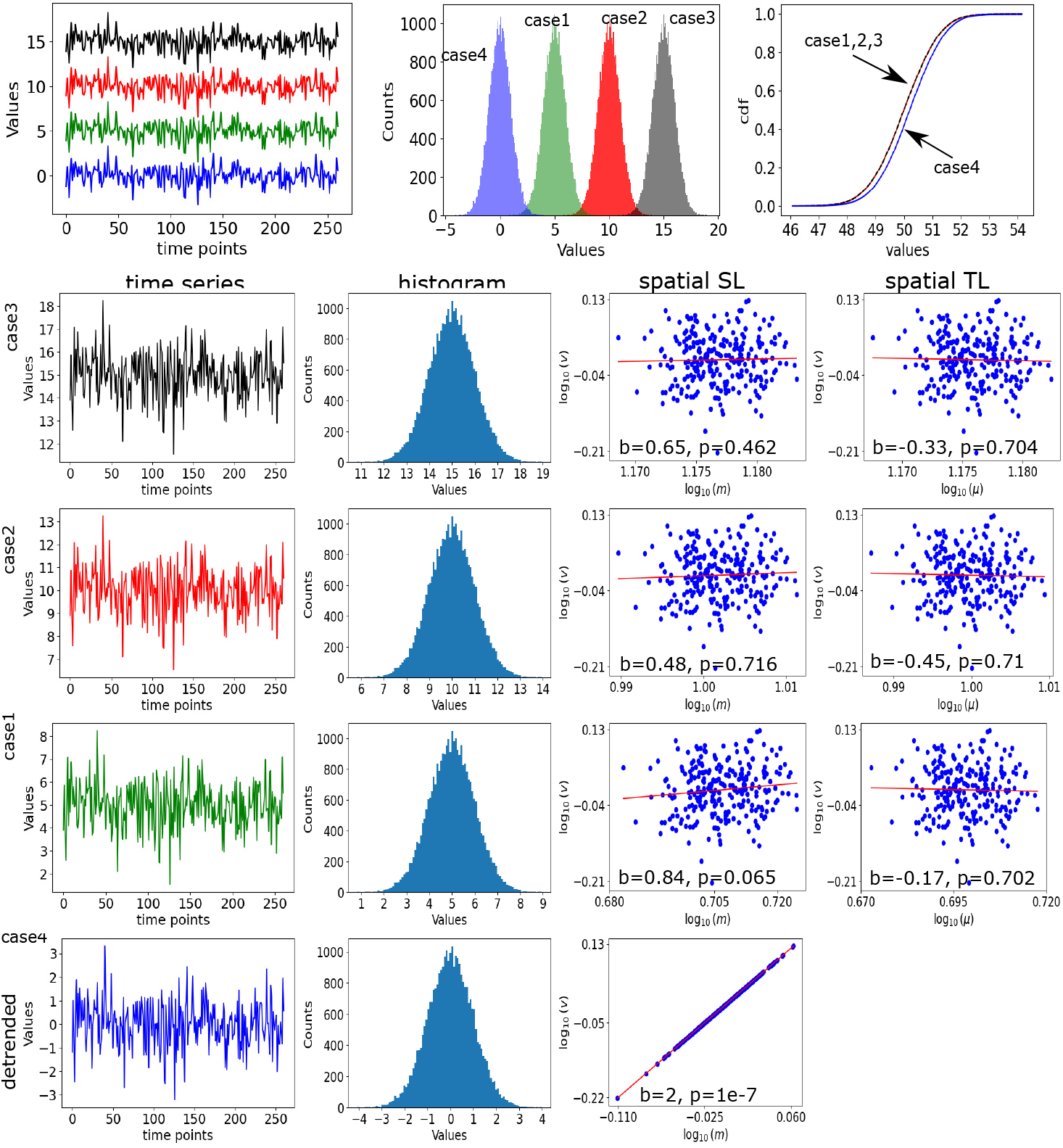
Normal distribution. These figures are generated by Python. The normally distributed spatio-temporal time series data was constructed using the built-in function “numpy.random.normal”. The algorithm is written in Algo. 4. The spatio-temporal time series in four cases are generated by changing the mean value in the Python function, as temporal mean=0, 5, 10, 15 (blue, green, red, black) shown in fifth, fourth, third and second rows, respectively.

**Figure S9.**
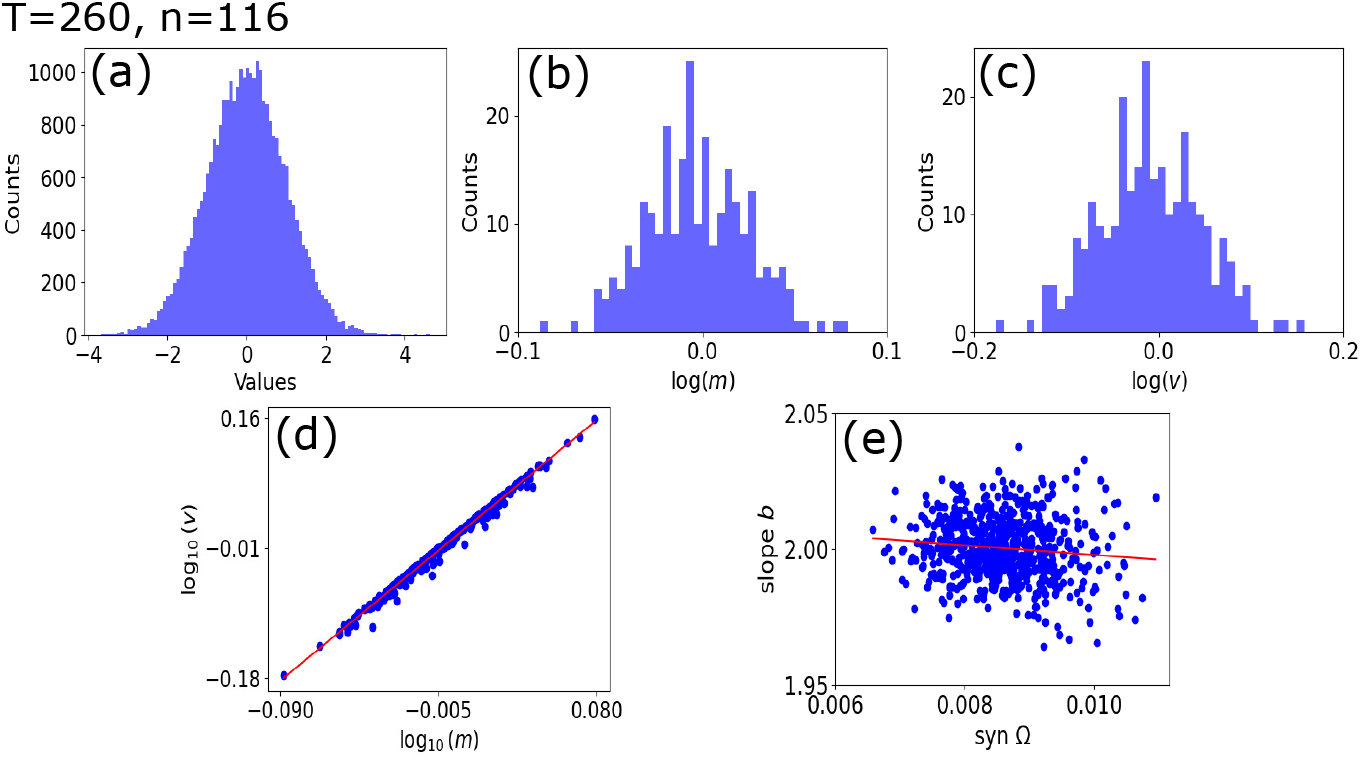
Normal distribution. (a-d) We used algorithm shown in Algo.4 and Python functions to create a spatio-temporal time series data with 260 time pints and 116 locations. (e) Total 645 realizations are made to construct iid data, where location-wise correlation is nearly zero. The SL slope is found nearly around 2 for the weak synchrony level.

#### 4.3 Identically distributed, but not necessarily independent

We have followed similar steps and mathematical expression proposed by DC Reuman et al. [2], using the well-known “common elements” method [13] of constructing a correlation-based multivariate random variable (*Y*_1_, …, *Y*_*n*_) such that the *Y*_*i*_ are identically distributed according to some specified distribution, and such that *cor*(*Y*_*i*_, *Y*_*j*_) = *ρ* for *i* ≠ *j* for Poisson, Negative Binomial, Gamma and Normal distributions.

##### 4.3.1 Poisson distribution

For identically distributed time series data *Y*_*i*_, drawn from Poisson distribution, using the common element method are presented here. Assume, a single time series variable *X* added to the spatio-temporal time series *X*_*i*_ to construct the random variable *Y*_*i*_. The statistical properties of the data *X* and *X*_*i*_ are modified by the average correlation *ρ*, as shown bellow.

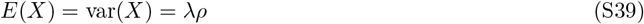

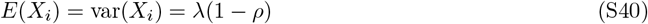

Defining *Y*_*i*_ = *X* + *X*_*i*_, it is well-known that *Y*_*i*_ is Poisson distributed, because the summands are independent, and *E*(*Y*_*i*_) = *E*(*X*) + *E*(*X*_*i*_) = *λρ* + *λ*(1 − *ρ*) = *λ* as desired. Also, for *i* ≠ *j*,

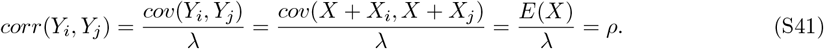

For more detailed descriptions and analytical derivations, please refer to Ref. [2]. Simulation results are shown in Fig. S10. We derived synchrony using Eq. (S15), found to be almost similar to the *ρ* value, and the statistical properties are numerically calculated, and then put to the Eq. S31 to derive the spatial SL exponent *b*. We have stored both the fitted slope and semi-analytical exponent values for different average synchrony values.

**Figure S10.**
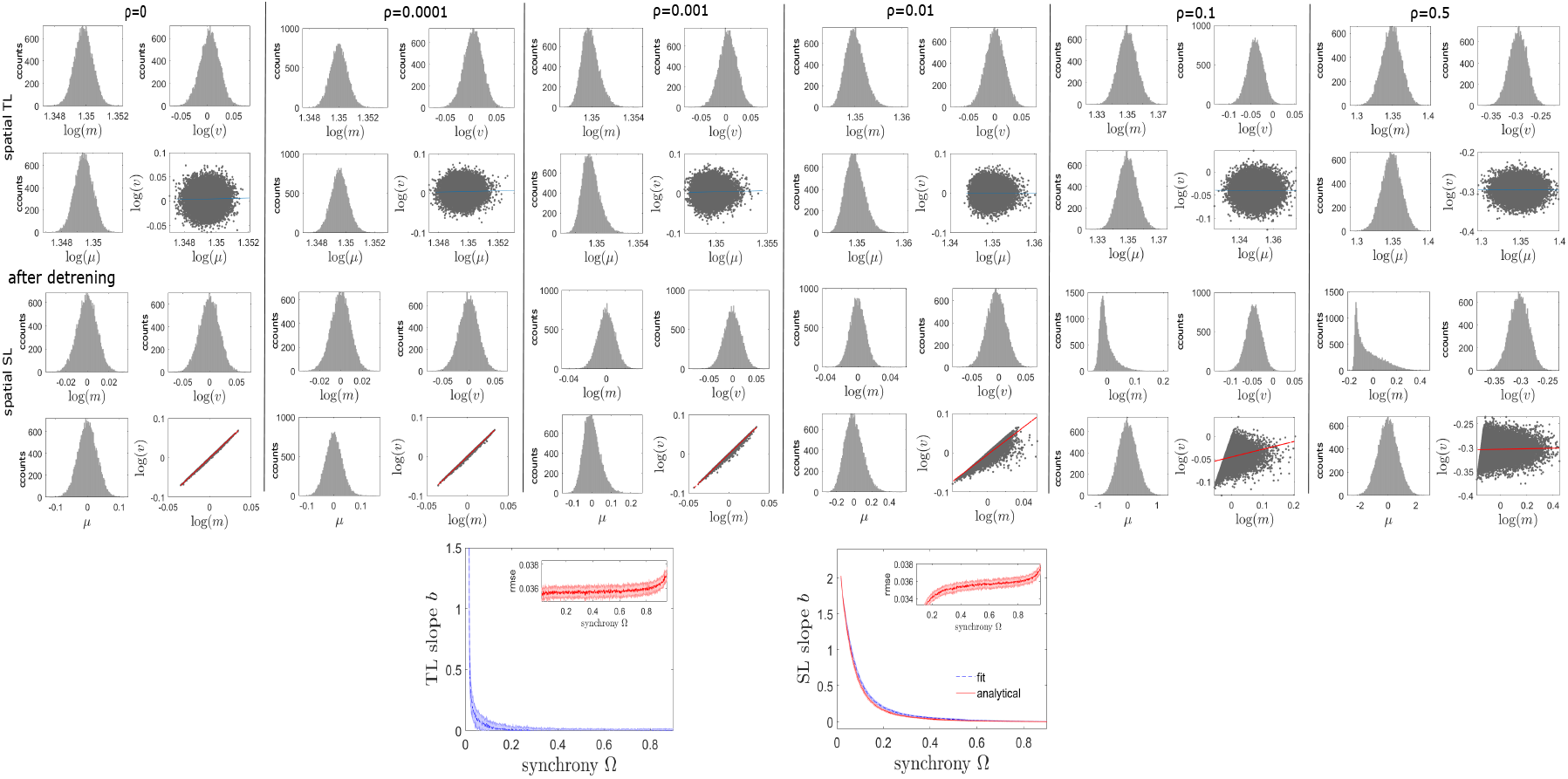
Influence of synchrony on the emergent of SL slope in synthetically generated multivariate random variable (*Y*_1_, …, *Y*_*n*_) has been tested. The random variable *Y*_*i*_ are identically distributed but not necessarily independent, constructed by randomly drawn values for a given Poisson distribution (*λ* = 50), using common element method [13], while controlling the correlation between *Y*_*i*_ and *Y*_*j*_ for all *i* ≠ *j*, with the parameter, *ρ*. The inset shows RMSE, calculated using Eq. S38. After detrending, the results are produced to visualize the spatial SL for various *ρ* values and also the spatial SL exponent against the synchrony are shown. As the average correlation increased from zero, the Sl exponent *b* almost always decreased quite sharply from 2.

##### 4.3.2 Negative Binomial distribution

We employed similar strategy to generate multivariate data drawn from negative binomial distributions, NB(*r, p*), where *p* denotes the probability of a success or of a failure, and *r* represents success or failure. Let X be a time series drawn from negative binomial distribution with parameters *r*_*X*_ = *ρr* and *p*, and let the *X*_*i*_ be independent (with respect to each other and *X*) negative binomials with parameters *r*_*i*_ = *r*−*r*_*X*_ = *r*(1−*ρ*) and *p*. Defining *Y*_*i*_ = *X* + *X*_*i*_, it is known that *Y*_*i*_ is negative binomial with parameters *r* = *r*_*X*_ + *r*_*i*_ and *p*, as desired; it is also easy to show, since the variance of a negative binomial with parameters *r* and *p* is *r*(1 − *p*)*/p*^2^, that cor(*Y*_*i*_, *Y*_*j*_) = *ρ*, as desired. Simulating draws from (*Y*_1_, …, *Y*_*n*_) is straightforward and computationally efficient. Standard formulas for negative binomial moments provide, *M* = *r*(1 − *p*)*/p*, and *V* = *r*(1 − *p*)*/p*^2^, for other moments and detailed analytical derivations refer to Ref. [2].

**Figure S11.**
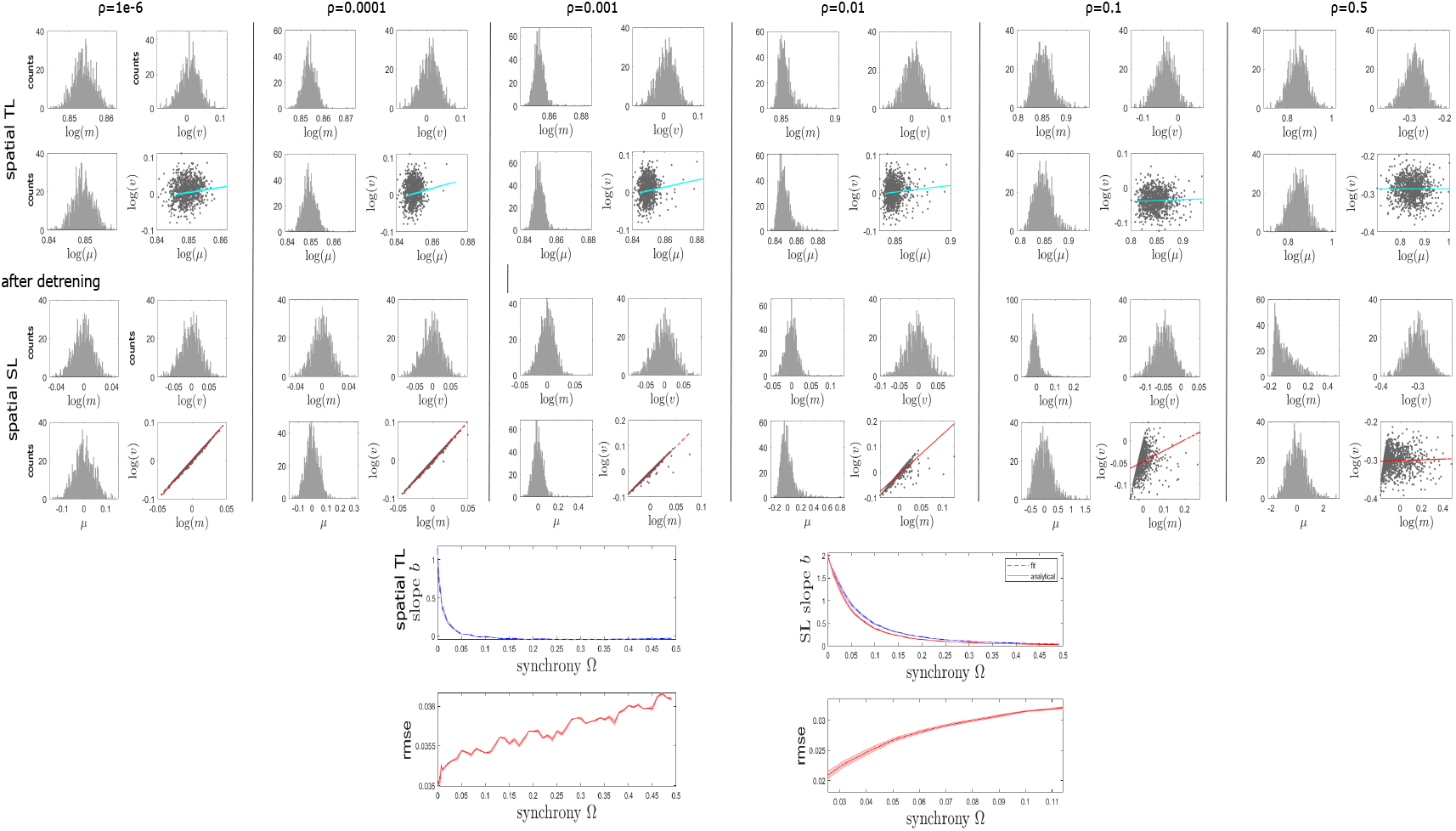
Influence of synchrony on the emergent of SL slope in synthetically generated multivariate random variable (*Y*_1_, …, *Y*_*n*_) has been tested. The random variable *Y*_*i*_ are identically distributed but not necessarily independent, constructed by randomly drawn values for a given negative binomial distribution and used common element method [13] to create *ρ* dependent spatio-temporal data, where corr(*Y*_*i*_,*Y*_*j*_)=*ρ* for all *i*≠ *j*. The inset shows RMSE, calculated using Eq. S38. After detrending, the results are produced to visualize the spatial SL for various *ρ* values and also the spatial SL exponent against the synchrony are shown. As the average correlation increased from zero, the Sl exponent *b* almost always decreased quite sharply from 2.

##### 4.3.3 Gamma Distribution

The same analytic strategy was applied to the gamma distribution Γ(*α, β*), where *α* is shape parameter and rate parameter is *β*. Now, let *ρ* is the induced correlation, then assume *X* follows gamma distributed with shape and rate parameters *ρα, β*. The *X*_*i*_ is independent to each other and with *X*, and is drawn from gamma distribution with parameters *α*(1 − *ρ*) and *β*. It is known that *Y*_*i*_ = *X* + *X*_*i*_ is gamma distributed, with shape parameter equal to the sum of the shape parameters of *X* and *X*_*i*_, *α* and rate parameter equal to the common rate parameter for the summands, *β*. A straightforward check can give cor(*Y*_*i*_, *Y*_*j*_)=*ρ* for *i* ≠ *j*, as desired. Simulating draws from (*Y*_1_, …, *Y*_*n*_) is again straightforward and computationally efficient. The first two moments were from standard formulas or were computed by inserting *Y*_*i*_ = *X* + *X*_*i*_ into the moment definitions and computing: *M* = *α/β*, and *V* = *α/β*^2^.

**Figure S12.**
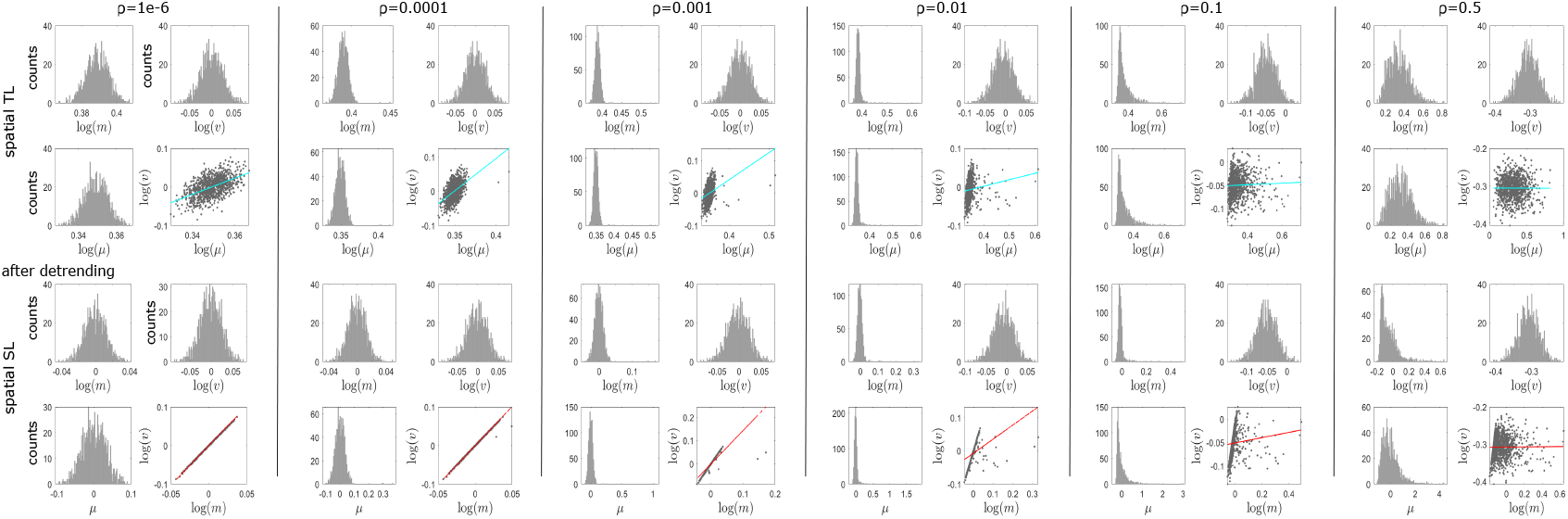
Variation in synchrony-induced SL exponents for different *ρ* in Gamma distributions.

##### 4.3.4 Normal Distribution

The same analytic strategy was applied to the normal distribution *N* (*µ, σ*), where *µ* is mean and *σ* is standard deviation. Now, let *ρ* is the induced correlation, then assume *X* is single time series dependent on *ρ*, which follows normal distributed, and let *X*_*i*_ is independent to each other and with *X. X*_*i*_ is drawn from normal distribution with dependency on the parameters (1 − *ρ*). It is known that *Y*_*i*_ = *X* + *X*_*i*_ is normal distributed. A straightforward numerical check can give cor(*Y*_*i*_, *Y*_*j*_)=*ρ* for *i* ≠ *j*, as desired. Simulating draws from (*Y*_1_, …, *Y*_*n*_) is again straightforward and computationally efficient.

**Figure S13.**
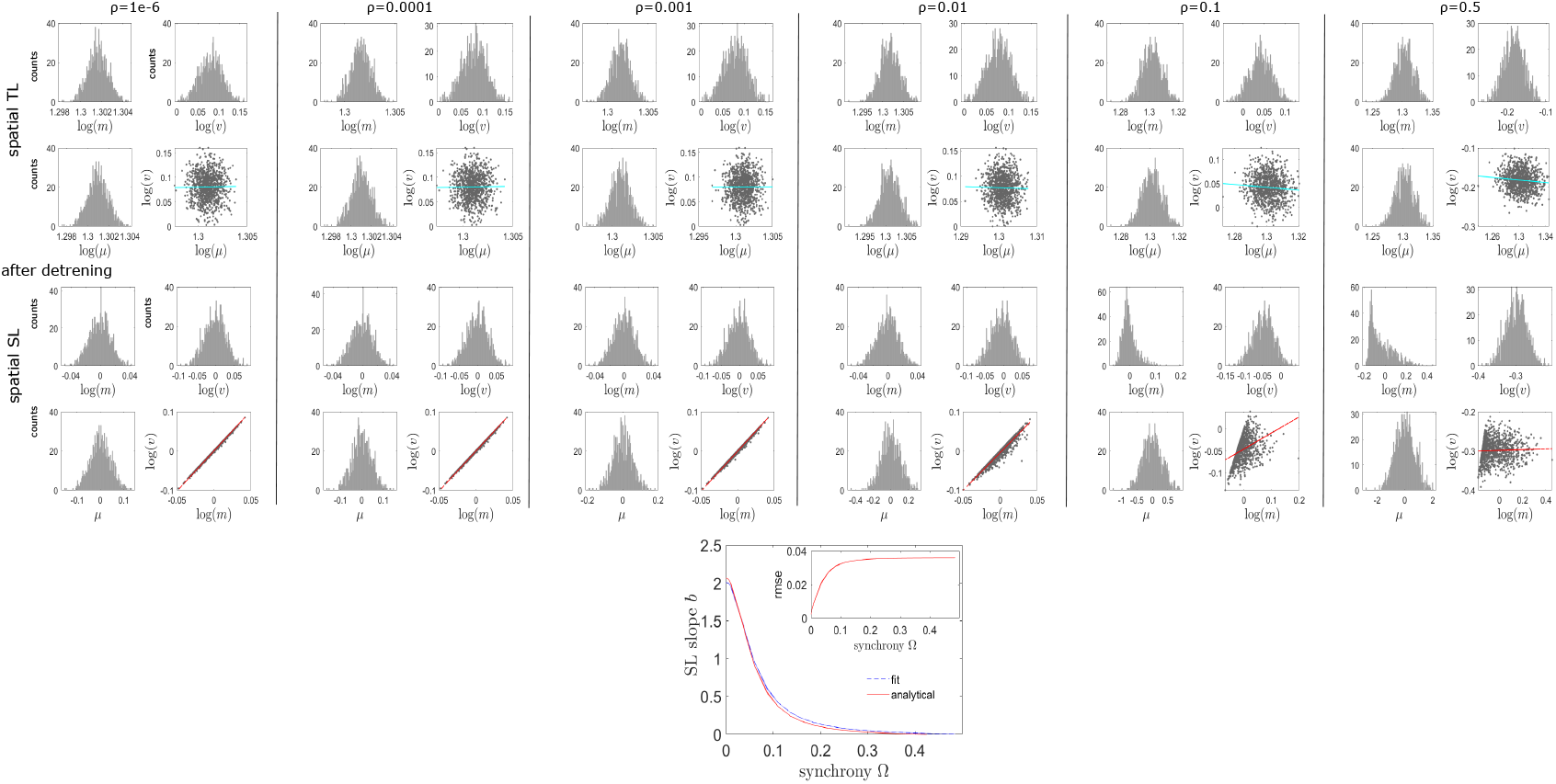
Variation in spatial SL exponents against synchrony in case of Normal distribution. We have shifted the mean values above zero for the cases of TL derivations and then, either detrended the data or generated it with zero mean using algorithm shown in Algo. 5.

### 5 Algorithm to construct multivariate time series

#### 5.1 Construct spatio-temporal time series: identically independent distributions

We used MATLAB default random generator, Mersenne ‘twister’ algorithm with ‘mt19937ar’ generator keyword and variable seeds for different observations in the three test cases, binomial, uniform and Gaussian normal distributions.

##### 5.1.1 Poisson distribution

###### Algorithm 1

Test time series have been generated by drawing values from Poisson distribution

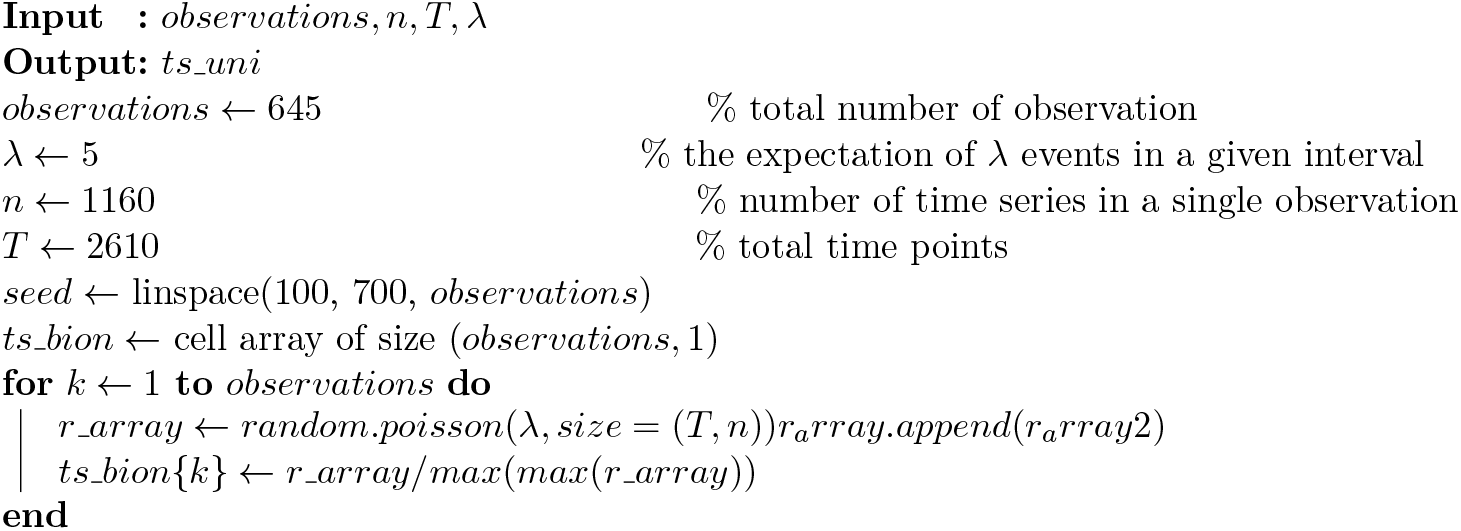

##### 5.1.2 Binomial distribution

###### Algorithm 2

Test time series have been generated by drawing values from Binomial distribution

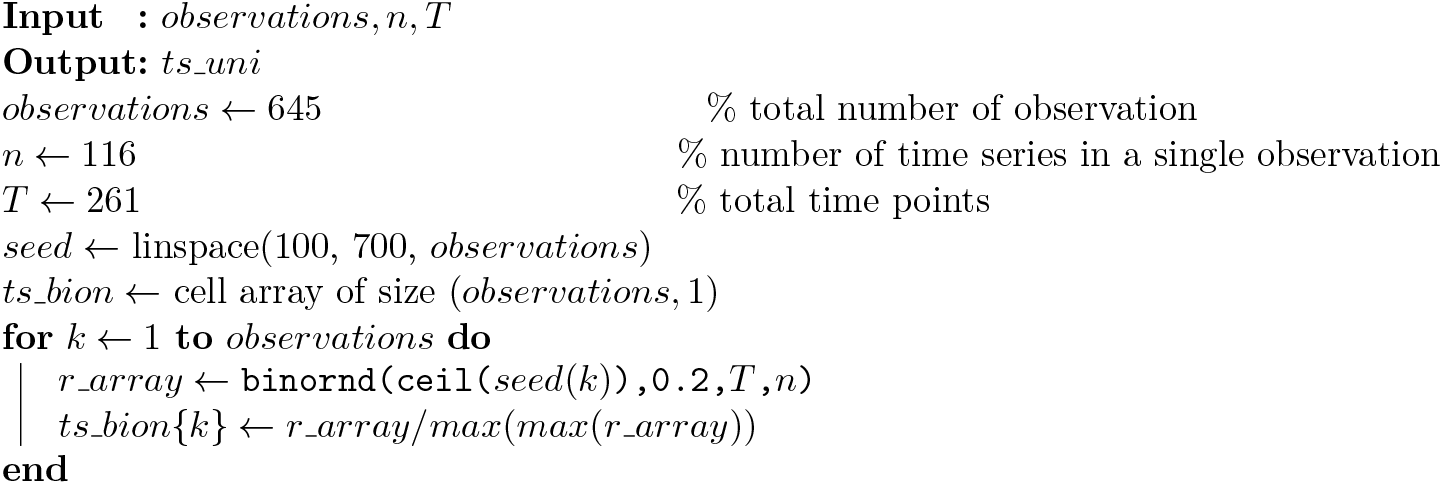

##### 5.1.3 Uniform distribution

###### Algorithm 3

Time series generated from uniform distribution

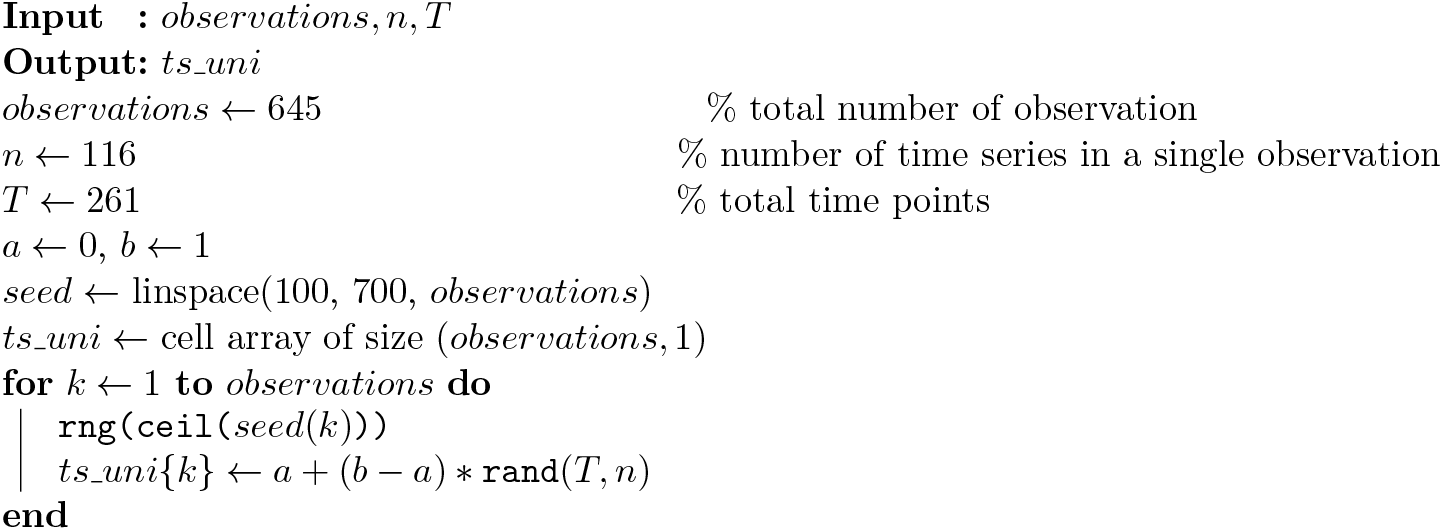

##### 5.1.4 Normal distribution

###### Algorithm 4

Time series generated (iid) from normal distributions in Python

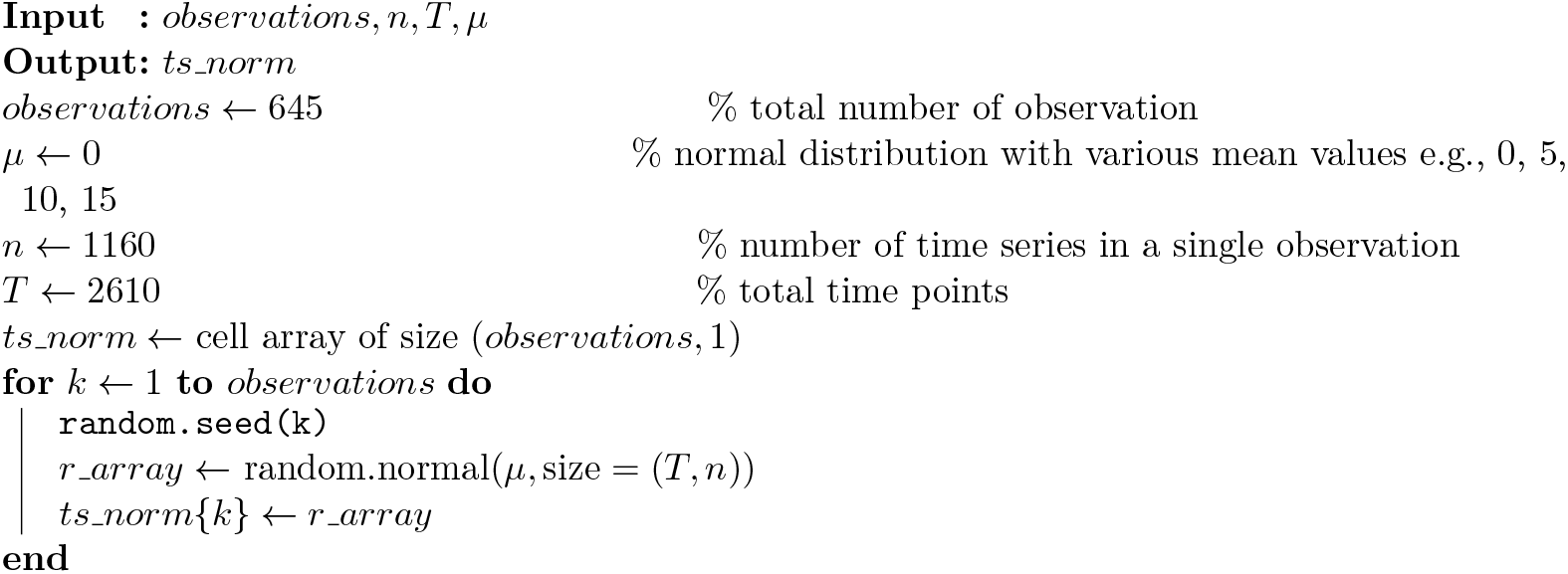

##### 5.1.5 Construct spatio-temporal time series: identically distributed, but not necessarily independent

###### Algorithm 5

The spatio-temporal time series is generated from normal distributions, using common element method.

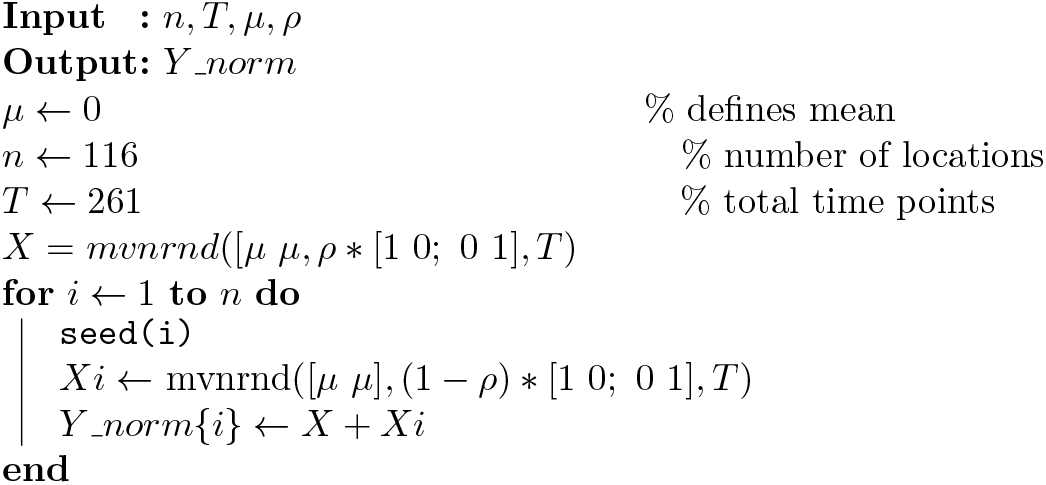

### 6 Empirical data analysis using CamCAN dataset

#### 6.1 Data sources and participants

We have used the data from http://www.mrc-cbu.cam.ac.uk/datasets/camcan. The data were collected as part of stage 2 of the Cambridge Centre for Aging and Neuroscience (CamCAN) project [14, 15]. The fMRI data in resting state (rest), naturalistic movie watching (mov) and sensorimotor task (smt) from 645 healthy participants were used in our study. The data is made publicly available and can be downloaded from their website.

**Table S1.**
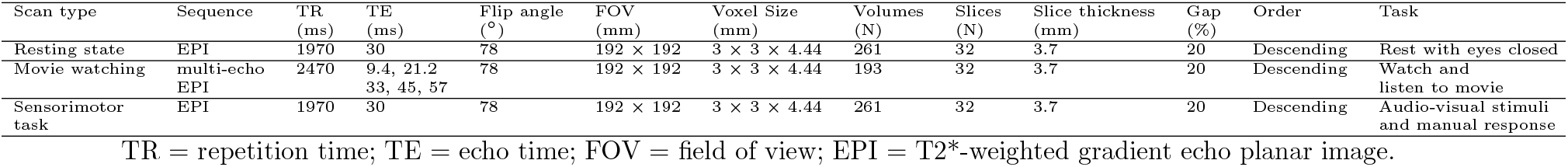
Functional MRI scans.

#### 6.2 Data preprocessing

The fMRI data for each functional run (resting state, movie watching, and SMT) were unwrapped using fieldmap images, realigned to correct for motion and, slicetime was corrected. EPI data were coregistered to the T1 image, transformed to montreal neurological institute (MNI) space using the warps and affine transformation from structural image (estimated using DARTEL). For region of interest (ROI) analysis, mean regional BOLD time series were estimated in 116 parcellated brain areas using Anatomical Automatic Labelling atlas (AAL) [8] (http://www.gin.cnrs.fr/tools/aal). Preprocessed data were provided by Cam-CAN research consortium. Details of preprocessing pipeline can be found in [14].

For group-level analysis, we divided the data of 645 participants into three cohorts, young adults (YA) age range 18-27 years, middle-aged adults (MA) age 46-55 years, and older adults (OA) age 75-84 years. Each participant’s BOLD time series in the resting state, naturalistic movie watching, and SMTs were used. We did region-wise detrending for each subject, resulted in zero ‘temporal mean’ [2, 16], i.e., the fluctuation of the regional time series is around zero, which also implies that the signals contain both positive and negative values.

#### 6.3 Surrogate time series: phase-randomization

We randomized phases of the empirical BOLD series to produce surrogated data utilized to examine marginal’s contributions to measurement. Given a time series *Y* (*t*), the complex valued Fourier transform can be written as,

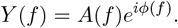

*A*(*f*) and *ϕ*(*f*) are amplitude and phase, respectively. We considered a univariate phase randomization using fast Fourier-transform surrogate data [17]. A ‘phase-randomized’ Fourier transform 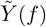 is made by randomly rotating the phase *ϕ* at each frequency *f* by an independent random variable *φ* which is chosen from uniform distribution in the range [0,*π*]. The *φ*’s are not completely independent, but instead are constrained to satisfy the symmetry property *φ*(*f*)=−*φ*(−*f*) to insure that the inverse Fourier transform is a time series of real values. That is,

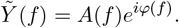

From this surrogate time series is given by the inverse Fourier transform by,

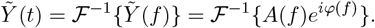

By construction, 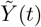 will have the same power spectrum as the original data set *Y* (*t*), and by the Weiner-Khintchine theorem the same autocorrelation function, i.e., the circular autocorrelation is preserved [17], when *Y* (*t*) has a zero temporal mean.

#### 6.4 Empirical Results

Neural encoding process [18] involves changes in firing rates and timing of neural population within certain brain region, and in synchrony, an emerged co-activation pattern across and within few regions in response to internal or external stimuli. Conventional studies leave answers to many important questions unclear, e.g., how accurately and efficiently can a mental state be inferred? We argued that if the process of neural encoding does indeed translate physical properties of a stimulus into patterns of brain activity that represents stimulus characteristics, then the signature can be extracted from the attributes of spatial fluctuation scaling of a time series.

Decoding a person’s current mental state by learning to recognize characteristic spatial patterns of brain activity associated with different mental states takes into account not just activity at single locations but the full spatial pattern of activity [18]. Such decoding method reveals that substantially more information is encoded in fMRI signals [18]. Three major constrains may mask non-linearity in the time series and can cause measurement error. For example, a recent paper argued in favor of linearity in the data, showing how macroscopic resting-state brain dynamics are best described by linear models [19]. They mentioned that although more than a few (around 100-10^4^) neurons are required, spatial averaging still dissolves the nonlinear aspects of the dynamics while mostly sparing the linear ones [19]. Secondly, there is an inevitably large time-scale separation between the observed slow varying blood oxygen level dependent activity (0.001Hz-0.1Hz) and underlying neural activity at different spatial brain scale in response to a given stimulus or spontaneous activity. The BOLD activity resulted from much faster neuronal activity at the microscopic level (e.g., single neuron action-potential), or mesoscopic level activity, averaging over an ensemble of neuronal population displaying higher frequency fluctuations (∼4Hz). Finally, microscopic nonlinear dynamics can be counteracted or masked by four factors associated with macroscopic dynamics: averaging over space and over time, which are inherent to aggregated macroscopic brain activity, observation noise and limited data sample, which stem from technological limitations [19].

The conventional approaches have proven successful in elucidating many aspects of the relationship between cognitive and mental states, and brain activity. However, recent advances in data analysis procedures raise the possibility of deciphering additional and complementary information from neuroimaging data [18]. Under this background, we envisaged a scale invariant linear relationship as a quantitative measure of spatial aggregation, while masking the effects from temporal mean and observation noise. In contrast to the strictly single location-based conventional analyses, such as temporal TL [16] or temporal autocorrelation [20], we have utilized spatial scaling law. The spatial SL holding statistical property of a multivariate time series is defined by the linear relationship when the spatial variance is approximately a power law scaling of spatial RMS upon a logarithmic transformation. Thus, the amplitudes of BOLD signals (*Y*_*i*_(*t*)) in all the brain regions include both positive and negative values evolving around zero temporal-mean [16,21]. Thereby, our analysis is focused on the attributes from spatial properties of BOLD signals while suppressing contributions from temporal features.

The study proposes a revised spatial scaling law (SL) derived from spatial distribution of individual’s detrended BOLD signals during different encoding processes for different brain states, including resting state, movie watching and sensorimotor task. Our primary aim is to find a feature less influenced by local temporal mean and data marginals. Next, we examine correlation between the spatial feature and synchrony, quantifying coordinated neural dynamics under different brain states, and finally, aging effect on the synchrony-scaling relationship.

#### 6.5 Empirical data visualization

Empirical data distributions before and after detrending operation for three brain states (rest, movie and sensorimotor task) are shown.

**Figure S14.**
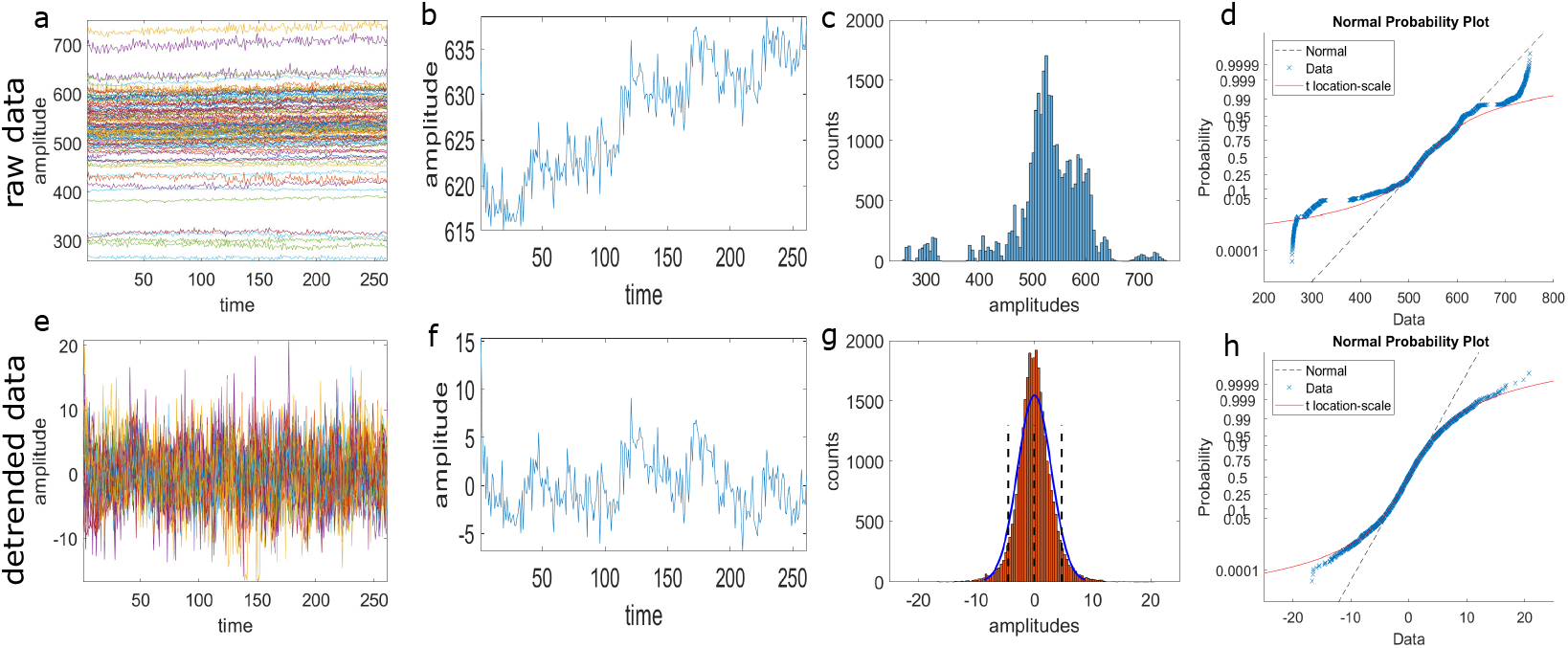
Empirical data and detrending operation in resting state. (a-d) Raw data is shown. (a) Empirical BOLD signal of one participant. Data size is 261 × 116, brain regions = 116 (AAL atlas), time points = 261. (b) BOLD series of region-111. (c) Distribution of the raw data. (d) Probability plot of the raw data to test the normal distribution. Detrended data is shown in (e-f). (g) Detrended data distribution. Normal fit is shown in red line. (f) Probability plot of the normally distributed data after detrending operation.

**Figure S15.**
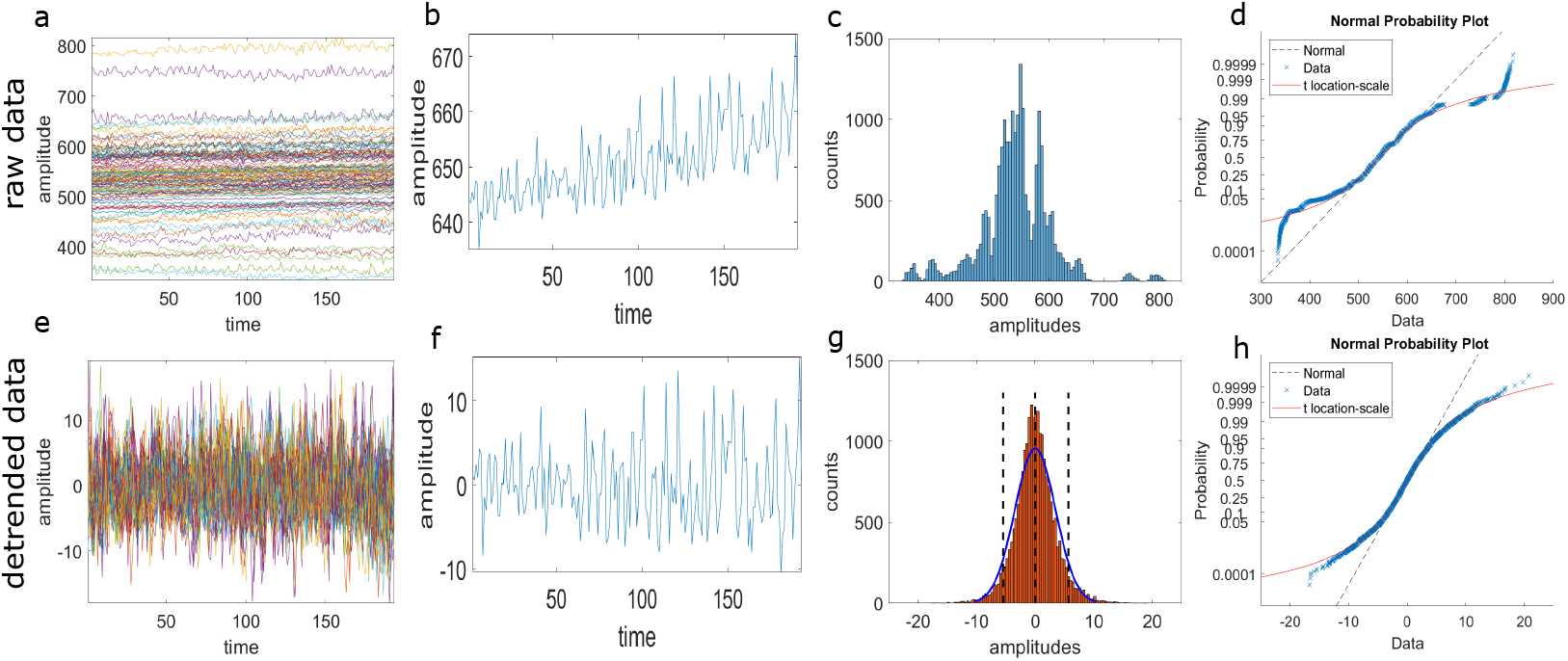
Empirical data and detrending operation in movie watching task. (a) Empirical BOLD series of one participant. Data size is 261 × 116; brain regions = 116 (AAL atlas), time points = 261. (b) BOLD signal of left cerebellum-10 (one location). (c) Histogram plot of the raw data. (d) Normal probability plot shows that neither the normal line (black dashed line) nor the location-scale curve (red) fits very well. After detrending, BOLD series of all the brain regions are shown in (e). (f) Suppressed trend of the BOLD signal of left cerebellum-10, compare (b). (g) Data distribution with normal fit in blue curve. The vertical black dashed lines indicate 95% CI. (h) Normal Probability plot of the detrended data shows that it fits well to the normal distribution for the data range within 95% CI.

**Figure S16.**
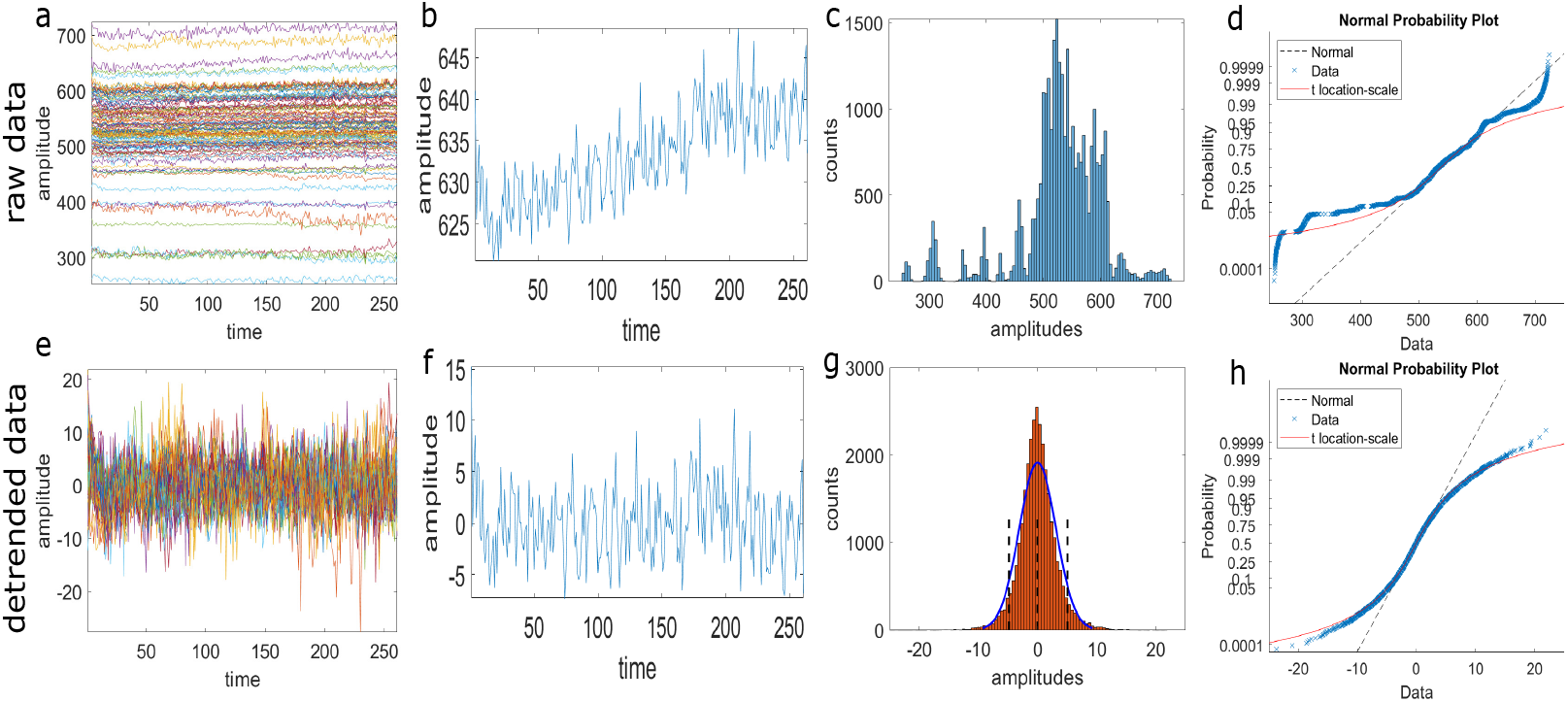
Empirical data and detrending operation in sensorymotor task. (a) Empirical BOLD series of one participant. Data size is 261 × 116; brain regions = 116 (AAL atlas), time points = 261. (b) BOLD signal of left cerebellum-10 (one location). (c) Histogram plot of the raw data. (d) Normal probability plot shows that neither the normal line (black dashed line) nor the location-scale curve (red) fits very well. After detrending, BOLD series of all the brain regions are shown in (e). (f) Suppressed trend of the BOLD signal of left cerebellum-10, compare (b). (g) Data distribution with normal fit in blue curve. The vertical black dashed lines indicate 95% CI. (h) Normal Probability plot of the detrended data shows that it fits well to the normal distribution for the data range within 95% CI.

**Figure S17.**
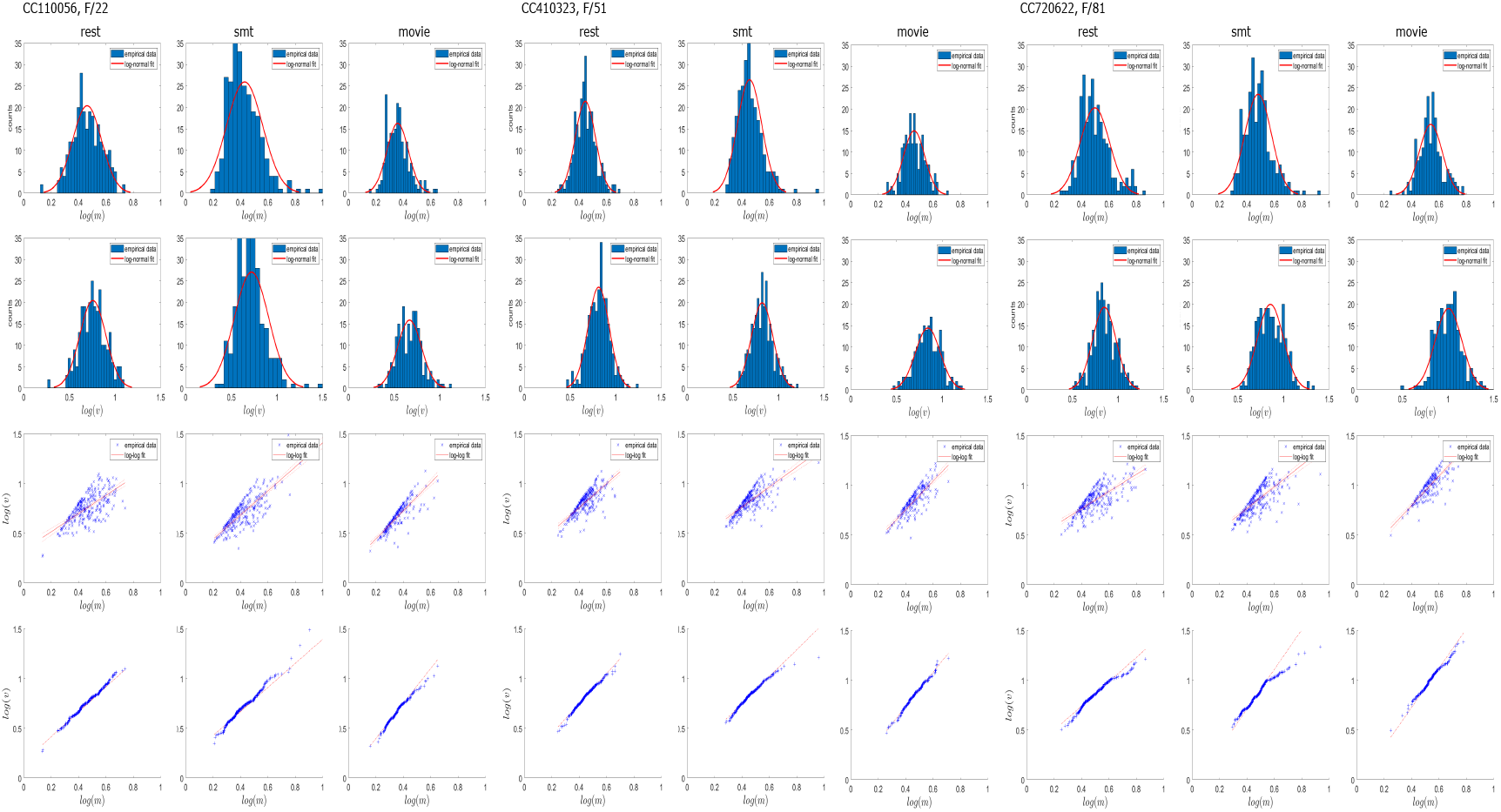
Functional MRI scans from three participants are chosen under three different brain states, such as rest, somatosensory task (smt), and movie. Top two rows show distributions of the spatial RMS and spatial variance with log normal fit. Third row from top shows a power law fitting between spatial RMS and variance on log-log scale. Red line indicates spatial SL exponent. Bottom row shows quantile-quantile plot.

**Figure S18.**
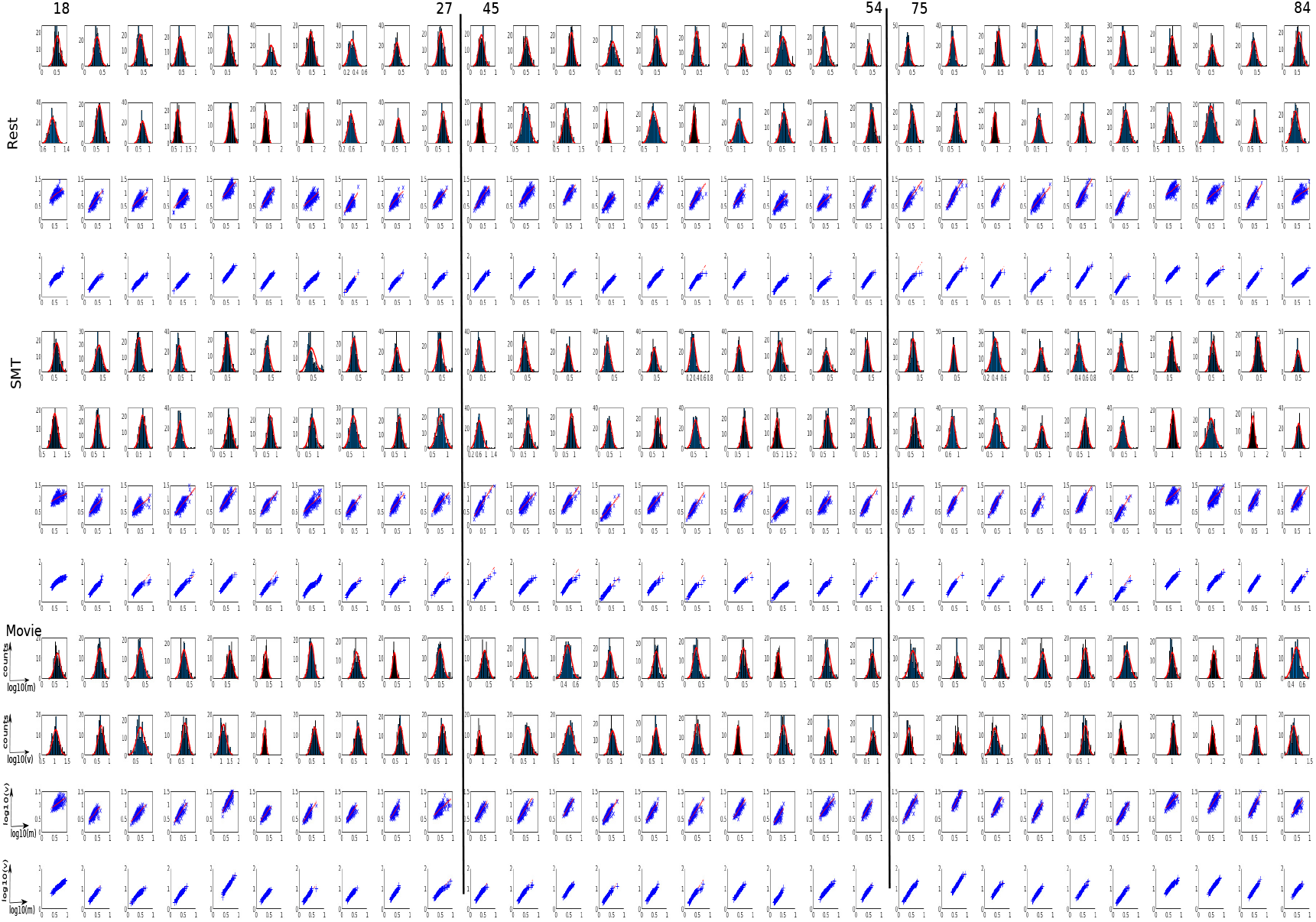
Ten participants are chosen from young, middle-aged and elder groups for three brain states. For each brain state, there are four rows. Top two rows show the log-normal distributions of the spatial RMS log(*m*(*t*)) and spatial variance log(*v*(*t*)). Third row shows log-log scatter plot between spatial RMS and variance. The red line indicates spatial SL slope. Bottom row shows quantile-quantile plot.

From the numerical simulations it is evidenced that any given distribution after detrending starts following normal-like distribution, which is set as a null model underlying random process or as a data marginal when spatial scaling exponent is not driven by the coordinated activity (at a very weaker strength of synchrony the spatial scaling 2). This has been utilized to remove the contributions or the effects of randomness on the empirical observations and measurements, so that we can obtain results solely driven by the internal or external stimuli. Thereby, We are able to decompose the spatial SL exponent *b* into two parts: the SL driven by random process *b*_*marg*_, and SL driven by the synchrony *b*_*syn*_.

#### 6.6 Task driven coordinated dynamics and spatial scaling law

We examine the influence of synchrony for both the resting state and active tasks using fMRI data from a large cross-section of adult lifespan (18-84 years old) subjects, *N* =645 participants, BOLD time series of size 260 time points×116 brain locations.

**Figure S19.**
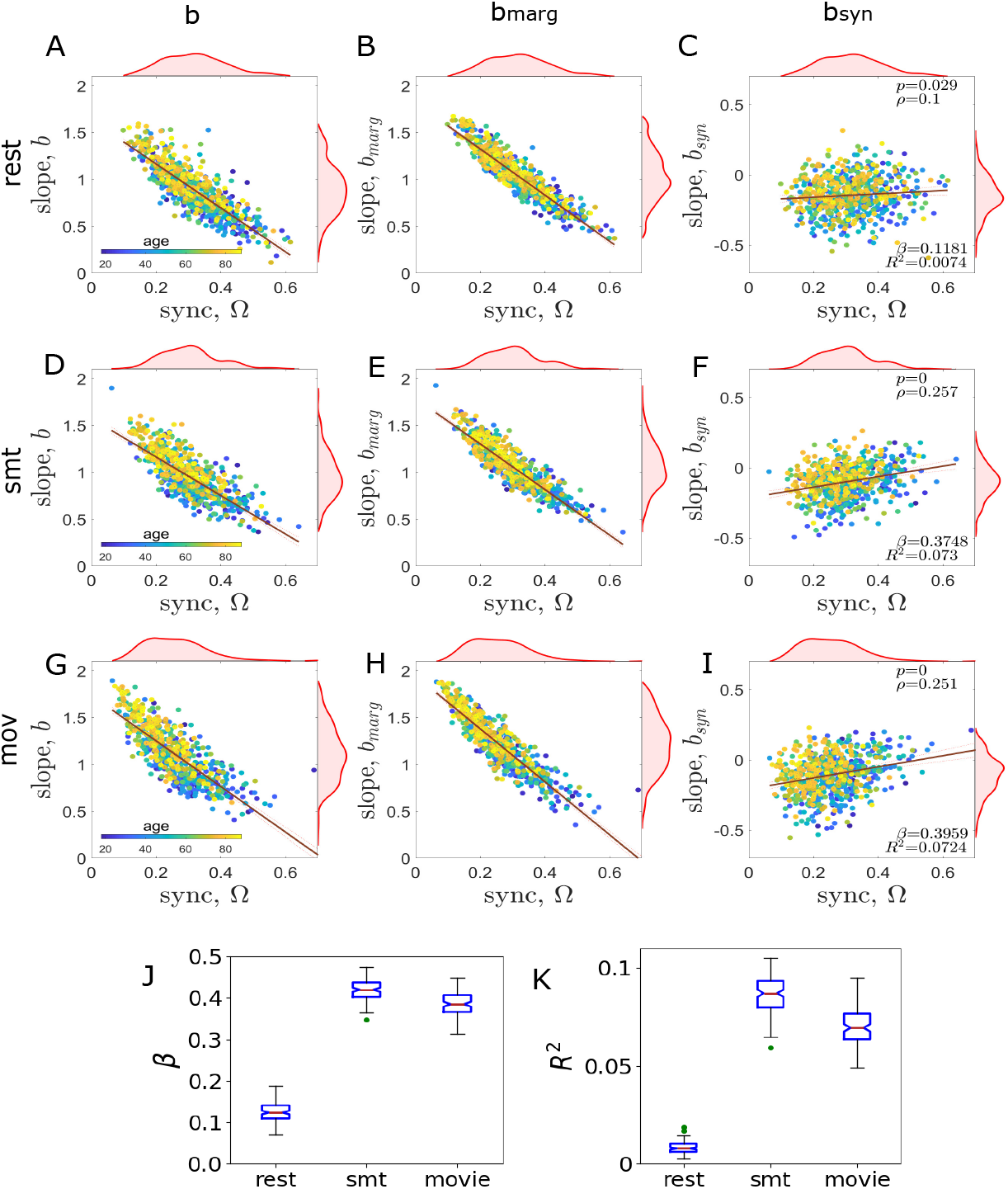
Plots of the SL exponent *b* and synchrony Ω for resting state (A-C), movie watch (D-F) and sensorimotor task (G-I) in CamCAN aging cohorts. First column (A,D,G) shows the scatter plot between synchrony (Ω) and the SL exponent (*b*) measured from the empirical data. Second column (B,E,H) shows scatter plot between Ω and the slope *b*_*marg*_, contributed by the data marginals (emerged from random process). The slope *b*_*marg*_ has been derived from the data constructed by randomizing the phases of empirical BOLD activity for three brain states. Third column (C,F,I) shows scatter plot fit between Ω and the slope *b*_*syn*_, derived by subtracting slope *b*_*marg*_, due to marginal distribution, from the original slope *b*, such that the synchrony induced SL slope can be clearly observed, when the contribution from the marginal distribution is separated from the measurement using *b*_*syn*_=*b*−*b*_*marg*_. (J) The degree of correlation is measured by the fitted slope (*β*) between Ω and *b*_*syn*_ for the three brain states, and corresponding *R*- squared values are shown in (K). We have measured the association slope (*β*) by phase randomization of the empirical data for 1000 independent realizations.

#### 6.7 Aging effects on synchrony and spatial SL in three brain states

We examine the effect of healthy aging on the spatial scaling. The SL slope is plotted against age as a continuous independent variable in Fig. S20; the linear regression fitting is shown by the red straight lines; blue curves show the slope by averaging over a window of five years. The synchrony-driven exponent *b*_*syn*_ increases (middle column) with increasing age in all three brain states. In a resting state, aging has no significant effect on the slope *b*_*syn*_, whereas the aging trajectory shows a significant increase in the slope *b*_*syn*_ against age for *smt* and *mov* cases. In other words, aging effects are prominently visible in the movie-watching and sensory-motor tasks but not in the resting state (rest). The third column in Fig. S20 shows a significant decrease in synchrony across the lifespan trajectory in all three brain states. The error bars (grey shades) are obtained for a number of participants present in each age value.

**Figure S20.**
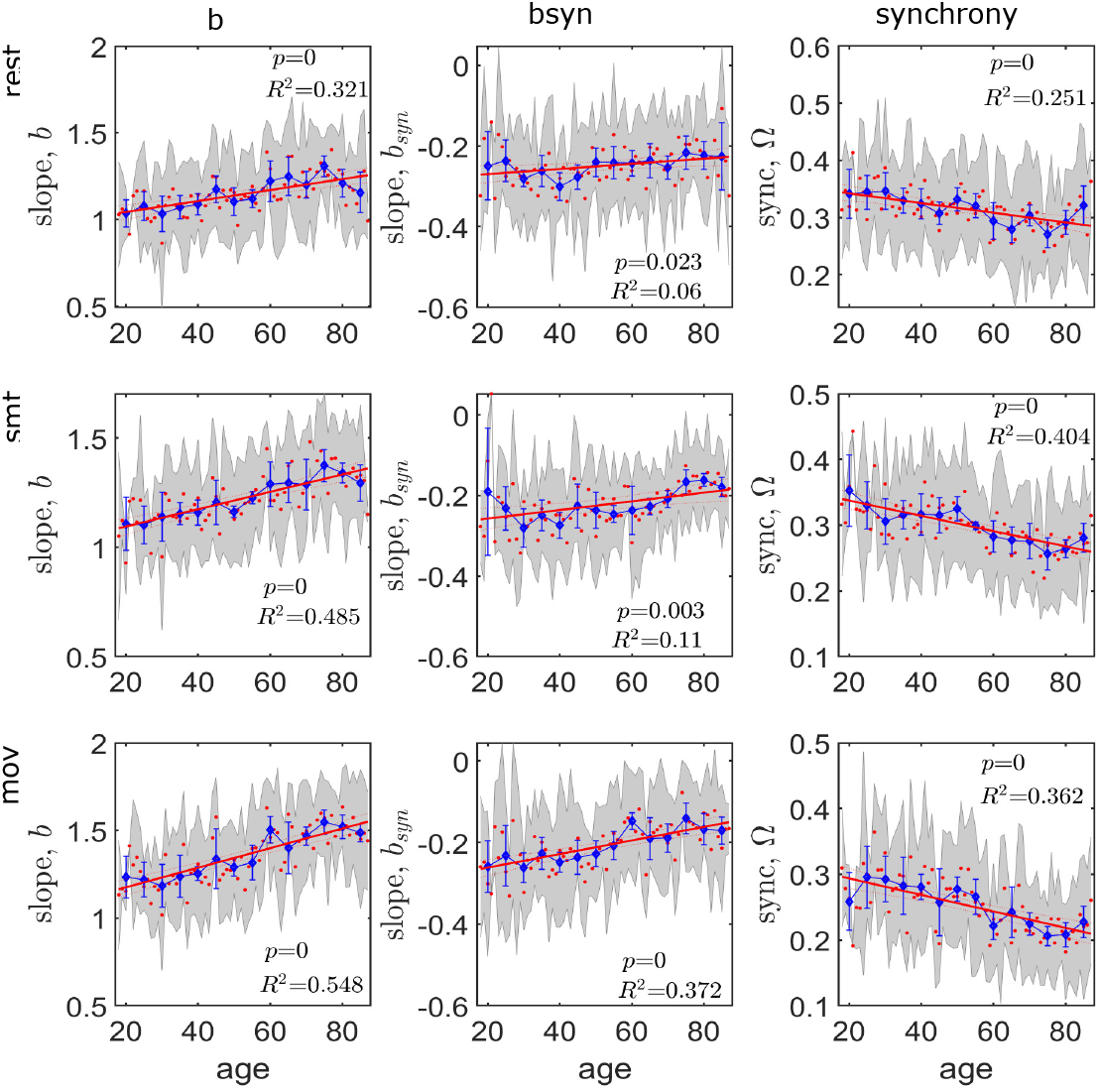
Effects of age on the synchrony-driven spatial SL for three mental states. We use least-square linear regression (red line) to fit *b*_*syn*_ and age, where age is the continuous independent variable. Blue curve is the slope value derived by averaging over five years bin. Year-wise standard deviation is shown by gray error shade. Details of the stats are given in the legend of each figure. Significance threshold is set at *p<*0.005.

#### 6.8 Group-wise analysis: Age-related alterations in the synchrony-induced SL exponent under three brain states

Associations between the spatial SL (*b*_*syn*_) and synchrony (Ω) are shown for three age groups in *rest* (A-C), *mov* (D-F) and smt (G-I) states.

**Figure S21.**
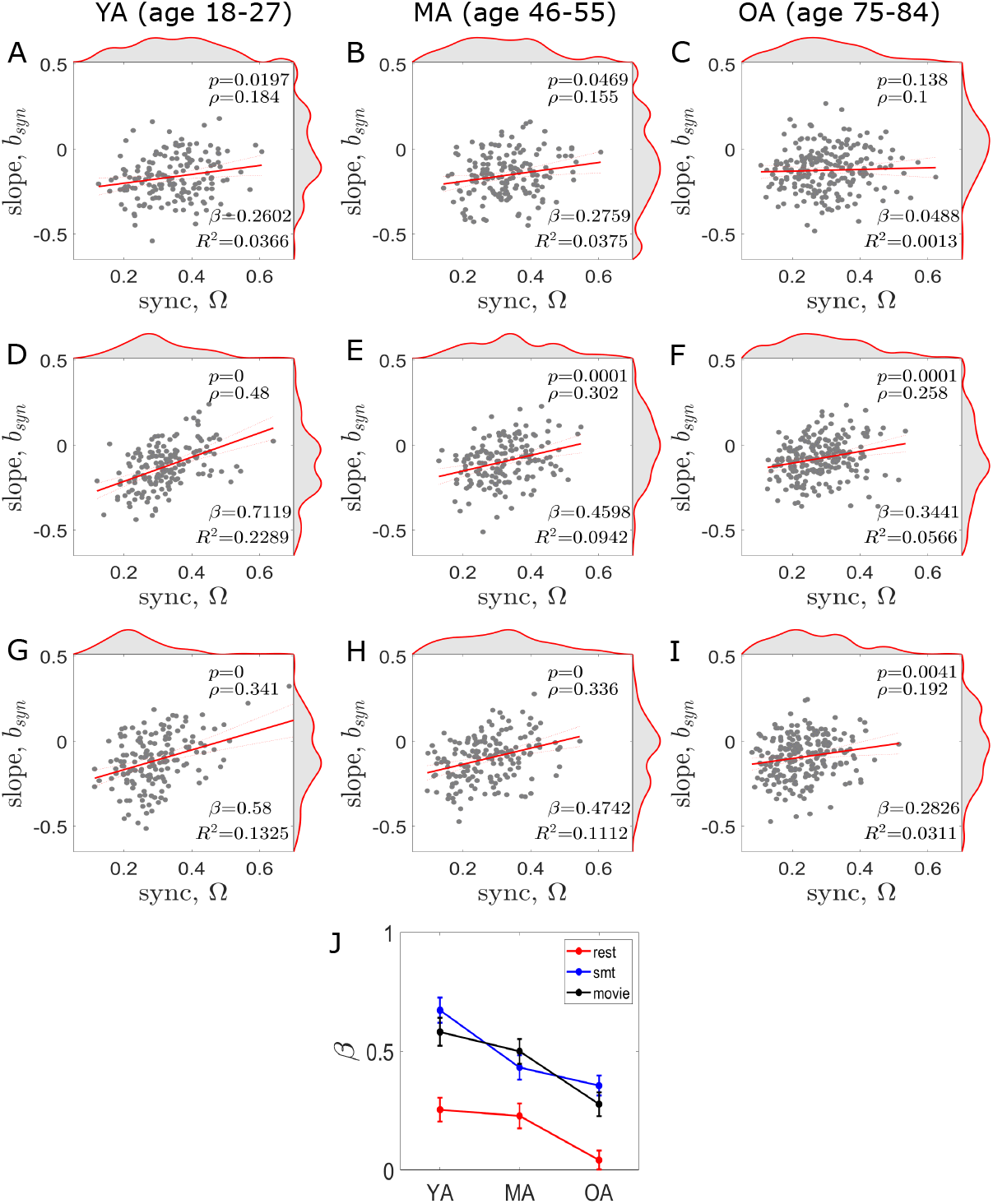
Age-related changes in the spatial SL, i.e., association between the spatial SL (*b*_*syn*_) and synchrony (Ω) are shown for three age groups in rest (A-C), movie (D-F) and smt (G-I) states. Three age groups are created, taking 10 years of bin. (A,D,G) In young adults (YA), the slope (*β*) remains positive and significant in all three conditions. (B,E,H) In middle-aged adults (MA), the association slope between the spatial scaling exponent and synchrony becomes flat, whereas, (C,F,I) in older adults (OA), the slop tends to negative. The age-related changes are much prominent in the movie watching task. However, it can be observed that the slope, *β* has a tendency to rotate clockwise with increasing age in all three brain states. (J) The slopes the plotted for three age cohorts in all three conditions. We conducted a repetitive simulations of 1000 times to check consistency in the association between synchrony and slope. The error bars indicate variations over the independent realizations.

### 7 Cross validation in NKI dataset: Healthy aging cohort

#### Data description and preprocessing

Resting-state BOLD activity is processed using AAL parcellation (116 ROIs) in Nathan Kline Institute (NKI)/Rockland dataset, (available in the UCLA Multimodal Connectivity Database (UMCD) [22] from total *N* =89 healthy participants, young: *N* =45 (22 Female), and old: *N* =44 (28 Female). Young group with age range 18-30 years, mean age 22.6222 ± 3.3864. Elderly group with age range 59-83 years, mean age 67.4545 ± 7.1315. Functional MRI (fMRI) for resting state have been preprocessed using the CONN toolbox [23] (https://web.conn-toolbox.org/), a Matlab/SPM-based software. Data were preprocessed using a default preprocessing pipeline of CONN. The preprocessing methods were unwrapping using field-map images, realignment to correct for motion, slice timing correction, segmentation, normalisation to the MNI template, outlier rejection and functional smoothing. A Gaussian kernel with of 2 mm and TR=2.5 sec are used to perform spatial smoothing. Denoising helped remove signal changes related to white matter, cerebrospinal fluid, motion, breathing and cardiac pulsations. Band-pass filter with 0.004-0.1 Hz pass-band frequency is applied. Linear detrending are performed. Mean regional BOLD time series were estimated in 116 brain areas of AAL atlas [8]. BOLD signal has size: Time Points×ROIs = 900×116.

**Figure S22.**
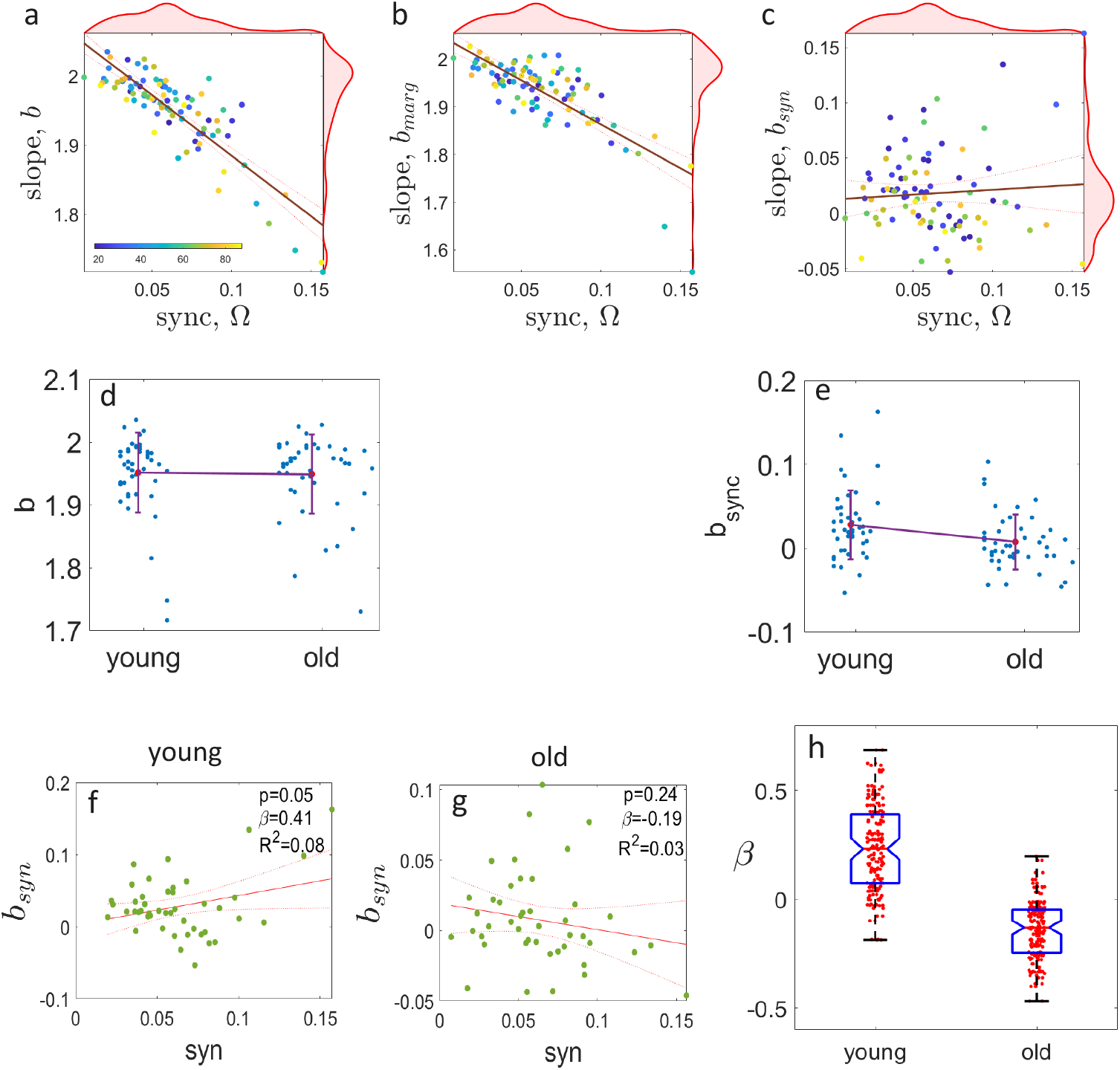
Cross validation in NKI dataset of healthy aging cohorts. (a,b,c) Spatial SL slope versus synchrony is derived from resting state BOLD activity. First panel shows raw slope, middle panel shows result for surrogate data by phase randomization, last panel shows synchrony-induced SL slope for entire the participants (*N* =45 (22 Female) young, and *N* =44 (28 Female) old). (d, e) The raw and synchrony-induced SL slopes are shown to compare between young and elderly adults. (f, g) Synchrony versus SL slope (*b*_*syn*_) is depicted for the two aging groups. With increasing age, the association between synchrony and Sl slope turns towards clock-wise direction, aligned to the results shown in Fig. S21(a) and (c) for YA and OA, respectively. (h) The effect of the synchrony on scaling behavior is derived for 100 independent realizations, which is aligned to the results for CamCAN dataset shown in red curve comparing YA and OA in Fig. S21(J). At the advanced age, the *β* value reduced significantly compared to the young cohort.

### 8 Synchrony-induced scaling in neural disorder in UCLA dataset

#### Data description and preprocessing

We have randomly selected *N* =66 healthy controls (HC) (33 Female) age range 21-50 years, mean age 31.12 ± 8.62, and Attention-Deficit Hyperactivity Disorder (ADHD) (*N* =40 (19 Female)) age range 21-50 years, mean age 32.05 ± 10.41 from publicly available dataset shared by the Consortium for Neuropsychiatric Phenomics (CNP) (<https://f1000research.com/articles/6-1262/v2>) [24]. This is a part of the large study funded by the NIH Roadmap Initiative that aims to facilitate discovery of the genetic and environmental bases of variation in psychological and neural system phenotypes, to elucidate the mechanisms that link the human genome to complex psychological syndromes, and to foster breakthroughs in the development of novel treatments for neuropsychiatric disorders [24]. The study includes imaging of a large group of healthy individuals from the community (66 subjects),and individuals diagnosed with ADHD (45). The participants, ages 21-50, were recruited by community advertisements from the Los Angeles area and completed extensive neuropsychogical testing, in addition to fMRI scanning. Participants were screened for neurological disease, history of head injury with loss of consciousness, use of psychoactive medications, substance dependence within past 6 months, history of major mental illness or ADHD, and current mood or anxiety disorder. More details can be found in Ref. [24].

Resting state BOLD signal of size: Time Points × ROIs = *T* ×*n*= 152×116, and AAL atlas of *n*=116 ROIs are used. Spatial scaling exponent (*b*) is derived first from the preprocessed BOLD signals of individual participants, where the violin plots presented in Fig. S23(a) are shown for HC and ADHD groups in blue and red, respectively. Figure S23(b) depicts SL exponents after removing marginals contributions. Figures S23(c, d) show the effect of synchrony on the SL exponent for HC and ADHD groups. The association between synchronization and the spatial scaling, captured by the slope *β* through least square linear regression fit, is found to be insignificant within the people diagnosed with ADHD, but very prominent in healthy controls. Essentially, the brain-wide coordinated activity has no effect on the scaling law in case of ADHD. In other words, the synchrony-scaling relationship breaks down in neural disorder. Further, we have performed 1000 independent realizations to confirm our observations, and derived the association slope *β*, which depicts a sharp contrast between the healthy and ADHD groups, see Fig. S23(e).

**Figure S23.**
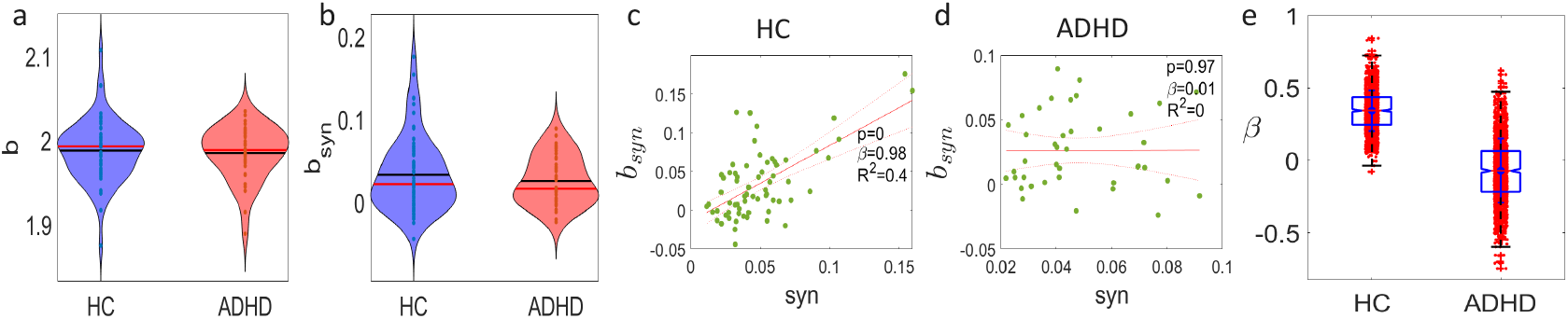
Synchrony-induced scaling law in neurodevelopmental disorder. (a, b) Synchrony effects on the SL exponent (raw and synchrony-induced) for healthy participants and ADHD subjects. (c, d) The influence of synchrony on the SL is shown for HC and ADHD groups is obtained for a single realization using linear regression fit, where the association slope is *β*. It is observed that the effect of synchrony on the SL becomes not significant (p*>*0.05) in ADHD compared to the HC group. (e) Furthermore, from 1000 independent realizations, the the value of *β* is derived for the HC and ADHD groups. It seems that the neural disorder has reduced the values of *β* profoundly, in comparison to the healthy controls.

### 9 Details of the Datasets

**Table S2.** Details of the empirical data from three independent cohorts. Total participants, *N* =840. Task conditions in CamCAN are resting state, movie watching and sensorimotor task. For NKI-Rockland and UCLA datasets, BOLD signals are obtained at eye open resting state.

| Dataset | Total | Group | Age range (years) | Mean age ± Std |
| --- | --- | --- | --- | --- |
| CamCAN | N=645 (326 Female) | Continuous age, N=645 | 18-88 | 54.24 ± 18.47 |
| | | Young, N=160 | 18-38 | 30.01 ± 5.51 |
| | | Middle, N=164 | 45-60 | 52.3 ± 4.68 |
| | | Old, N=223 | 65-88 | 75.08 ± 6.16 |
| NKI | N=89 (50 Female) | Young, N=45 | 18-30 | 22.62 ± 3.39 |
| | | Old, N=44 | 59-83 | 67.46 ± 7.13 |
| UCLA | N=106 (52 Female) | HC, N=66 | 21-50 | 31.12 ± 8.62 |
| | | ADHD, N=40 | 21-50 | 32.05 ± 10.41 |

### 10 Comparison with other metrics : Metastability, phase synchrony, entropy of functional connectivity dynamics

In this section, we employ three distinct global metrics widely used in neuroscience to investigate age-related changes in brain dynamics. This analysis enables us to identify specific characteristic changes and compare them with our main findings regarding synchrony-induced spatial SL exponents.

#### 10.0.1Metastability and Phase Synchrony

Recent brain network computational models suggest that the human brain achieves maximum metastability during resting states [25, 26] . This manifests as optimal information processing capability and switching behavior [25, 27–29]. Furthermore, the concept of metastability to measure brain dynamics enhances our understanding of cognitive aging theories [29–31] and help explore physiological substrate changes during aging [31, 32]. We have selected metastability as one of the global metrics to capture alterations in the brain-wide dynamics by comparing among different brain states across the adulthood aging trajectory. Metastability quantifies the continuous transitional states observed in brain signals without settling into specific attractor states [33–35] and serves as a metric for capturing the brain’s dynamic operating point during rest [25, 30, 36]. Metastability captures temporal fluctuation by estimating brain regions’ tendencies to deviate from coherent states. This provides an effective measurement unit for observing spatial cohesion during short periods (hours) and may serve as a key component for observing gradual changes in lifespan brain neural dynamics. We must mention here that the Kuramoto order parameter is a valid measure in rhythmic oscillations, thereby, the rhythmicity in BOLD signal is achieved by applying a narrow band filter that allows pass-band frequency ranging from 0.001 Hz to 0.1 Hz, prior to compute metastability and synchrony measure. The measurement begins with calculating the Kuramoto order parameter [37], *r*(*t*), using,

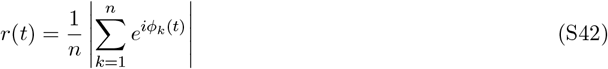

where, *ϕ*_*k*_(*t*) is the instantaneous phase of *k*^*th*^ brain region. The value of *r*(*t*) lies between 0 ≤ *r*(*t*) ≤ 1; *r*(*t*) = 1 representing complete synchronous, *r*(*t*) = 0 for desynchrony, and (0 *< r*(*t*) *<* 1) representing partial synchrony among all brain regions. Hence, mean of *r*(*t*) indexes the global coherence among a system of oscillating brain regions. Metastability (meta) can be calculated by taking the standard deviation of *r*(*t*) over the time:

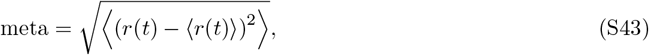

where ⟨.⟩ denotes average over time. The global phase synchronization (syn) is measured by taking average over time as,

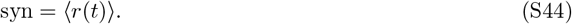

The dynamical measure of metastability offers several advantages: it quantifies neural systems’ tendencies to operate in both segregative and integrative modes [35], characterizes criticality states where dynamical systems remain at the edge of phase transitions with enhanced information processing capabilities [25, 38], and enables investigation through generative neural dynamics models, providing mechanistic insights into network processes generating brain signals [36, 39, 40]. Based on these insights, we employ metastability as the dynamical measure to capture brain dynamics help compare against our proposed metric, the synchrony-induced scaling exponent, while capturing alteration patterns in different brain states along the adulthood aging.

#### 10.0.2 Entropy of functional connectivity dynamics

Quantifying the temporal stability of dFC patterns is of immediate concern to studies investigating the relationship between resting state and task-related brain dynamics (Li, Lu, & Yan, 2019). Recent studies show that the correlations among brain regions are not stationary, but evolve over time, and have refocused the study of spontaneous brain activity on characterizing these time-varying functional interactions. the synchrony between the BOLD activities of different brain regions displays global slow fluctuations that reflect the dynamical association and dissociation of functional synchronized clusters. Emerging evidence suggests that FC dynamics FC may charaterize alterations in macroscopic neural activity patterns underlying critical aspects of cognition and behavior. Temporal similarity is captured by measuring entropy of the functional connectivity dynamics (FCD) [41–43], the time-time correlation matrix. This helps us to examine dynamical pattern of invariance or alterations in the temporal stability across slow time scale, i.e., lifespan aging. Static FC is quantified with metrics such as correlation, covariance, and mutual information between the time series of different regions, wherein the temporal and spatial scales examined are determined by the question of interest (Bressler and Menon, 2010; Bullmore and Sporns, 2009; Friston, 2011). It therefore represents an empirical characterization of the temporal relationship between regions, without indicating how the temporal covariation is mediated (Friston, 2011; Friston and Buchel, 2007).

Instantaneous phases *θ*(*i, t*) of the BOLD signals of *i* = 1, …, *n* (*n* = 116) regions are derived using Hilbert transform. Phase similarity between *i*^th^ and *j*^th^ brain regions at *t* time point is captured by the stream of FCs, computed by

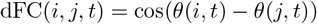

where *i, j* = 1, …, *n*. Principal component analysis (PCA) reduces the state-space dimension into a much lesser dimension, e.g., the dFC matrix of size *n* × *n* is reduced to size 3 × *n*, considering top three principle components, three largest eigen values. We express the original vector space as, **dFC**_*t*_=**V DV** at time point *t*, where **V** is the eigenvector of size *n*×*n*. The diagonal of **D** contains sorted eigen values, e.g., *λ*_1_ *> λ*_2_ *>* … *> λ*_*n*_, then, 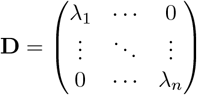.

If *k* is the number of principal components chosen for the analysis, and reduced subspace at time point *t* is **FCD**_*t*_; the reduced-dimension eigen matrix is 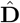, corresponding eigenvector is 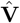, then we have

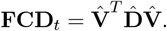

Temporal stability of the dominant subspace **FCD**(*k, n, t*) is estimated by the similarity between two FCDs at any two given time points (say, *t*_*x*_ and *t*_*y*_) is measured using angular distance. The principal angle between the **FCD** subspaces from different time points is calculated by the temporal stability matrix [44, 45], say Φ of size *T* × *T*, where *T* is the total time points, then the angular distance between two time points, *t*_*x*_ and *t*_*y*_ is derived by,

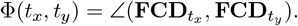

We have quantified complexity of temporal stability matrices by measuring entropy for all three categories, rest, movie viewing and sensorimotor task across the adulthood aging trajectory. The entropy is defined as,

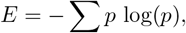

where *p* contains the normalized histogram counts of the temporal stability matrix.

#### 10.0.3 Results

The dynamical measure of metastability presented in Figs. S24(a-c). No significant changes are observed in resting state [Fig. S24(a)], but significantly decreased in the sensory motor and movie watching tasks; as seen in Figs. S24(b,c). The synchrony derived by taking time average of the Kuramoto order parameter is used here to effectively track how global coherence changes with age, and also help compare the results against the metric, spatial synchrony Ω. Significant decreasing patterns are observed in the global phase synchrony with increasing age, as seen from Figs. S24(d-f) in all three brain states, which are consistent with the results presented in the main text, Fig. 5 (third column). Both the metrics have captured significant decline in the global coherence level with age. Figures S24(g-i) illustrate changes in the functional connectivity dynamics (FCD) against age. No significant changes are observed in resting state; see Fig. S24(g), where as entropy significantly increased with age in the sensory motor and movie watching tasks; as seen in Figs. S24(h,i). In summary, it can be concluded that age-related changes are prominent in task evoked activity rather than resting state, whereas age profoundly affects brain-wide coordinated activity measured by synchrony.

**Figure S24.**
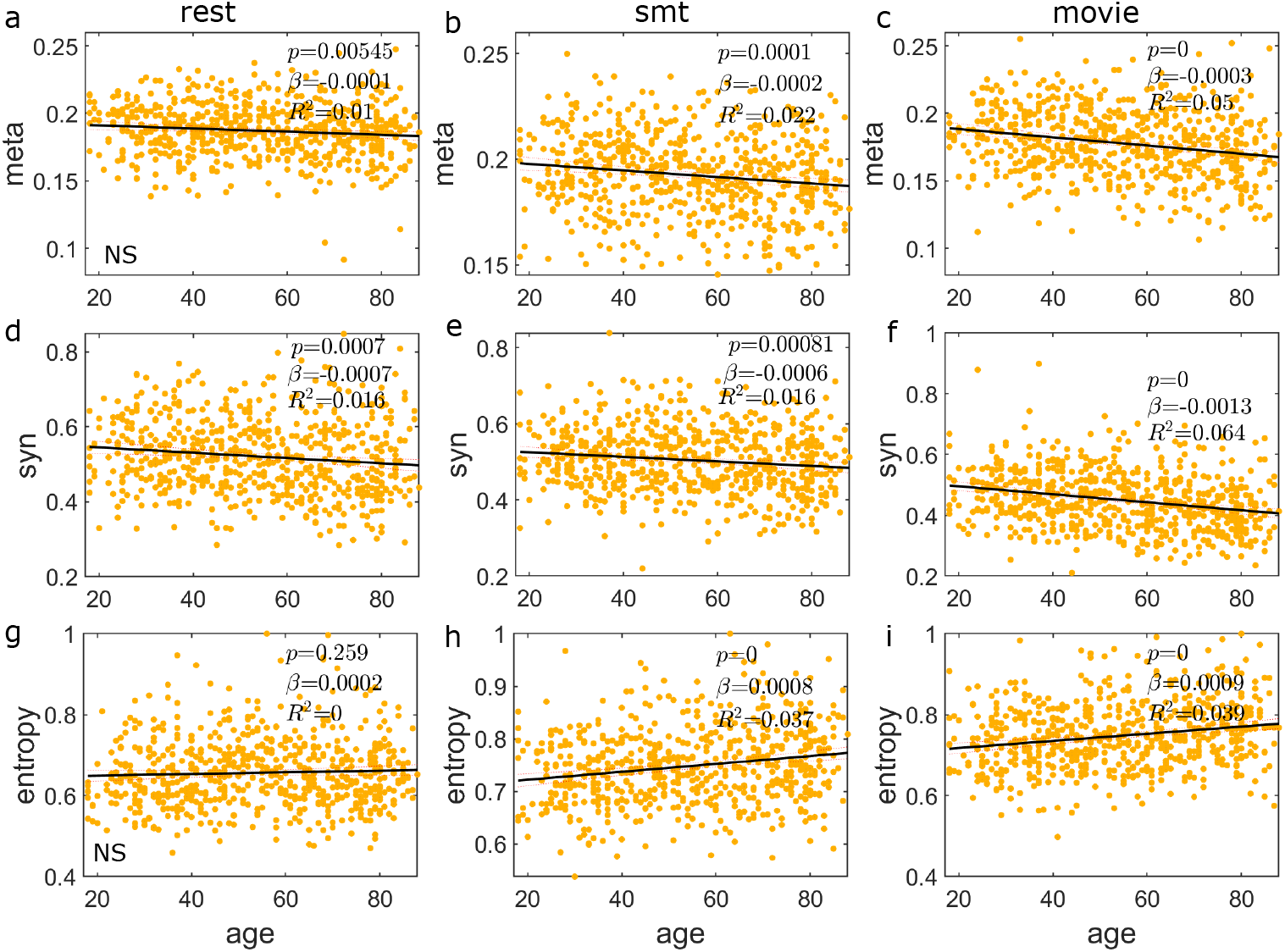
(a-c) Metastability, (d-f) entropy of functional connectivity dynamics, and (g-i) global phase synchrony are plotted against age, arranged from top to bottom rows, where columns from left to right represent resting state (a,d,g), sensory motor (b,e,h) and movie watching task (c,f,i) based results. Statistical significance level is defined by *p <* 0.005, otherwise marked as not significant (NS). We have fitted the scatter plots using least square linear regression model, where age is the independent variable.

